# Multi-Subject Stochastic Blockmodels for Adaptive Analysis of Individual Differences in Human Brain Network Cluster Structure

**DOI:** 10.1101/672071

**Authors:** Dragana M. Pavlović, Bryan R. L. Guillaume, Emma K. Towlson, Nicole M. Y. Kuek, Soroosh Afyouni, Petra E. Vértes, Thomas B. T. Yeo, Edward T. Bullmore, Thomas E. Nichols

## Abstract

There is great interest in elucidating the cluster structure of brain networks in terms of modules, blocks or clusters of similar nodes. However, it is currently challenging to handle data on multiple subjects since most of the existing methods are applicable only on a subject-by-subject basis or for analysis of a group average network. The main limitation of per-subject models is that there is no obvious way to combine the results for group comparisons, and of group-averaged models that they do not reflect the variability between subjects. Here, we propose two novel extensions of the classical Stochastic Blockmodel (SBM) that use a mixture model to estimate blocks or clusters of connected nodes, combined with a regression model to capture the effects on cluster structure of individual differences on subject-level covariates. Multi-subject Stochastic Blockmodels (MS-SBM) can flexibly account for between-subject variability in terms of a homogenous or heterogeneous effect on connectivity of covariates such as age or diagnostic status. Using synthetic data, representing a range of block sizes and cluster structures, we investigate the accuracy of the estimated MS-SBM parameters as well as the validity of inference procedures based on Wald, likelihood ratio and Monte Carlo permutation tests. We show that multi-subject SBMs recover the true cluster structure of synthetic networks more accurately and adaptively than standard methods for modular decomposition. Permutation tests of MS-SBM parameters were more robustly valid for statistical inference and Type I error control than tests based on standard asymptotic assumptions. Applied to analysis of multi-subject resting state fMRI networks (13 healthy volunteers; 12 people with schizophrenia; *N* = 268 brain regions), we show that the Heterogeneous Stochastic Blockmodel estimates ‘core-on-modules’ architecture. The intra-block and inter-block connection weights vary between individual participants and can be modelled as a logistic function of subject-level covariates like age or diagnostic status. Multi-subject Stochastic Blockmodels are likely to be useful tools for statistical analysis of individual differences in human brain graphs and other networks whose prior cluster structure needs to be estimated from the data.

## 1. Introduction

Network-like representations of brain connectivity (e.g., functional, structural or causal associations) allow us to explore the link between the architecture of the brain and the way it facilitates (a) specialised segregative processes and complex integrative processes. Two network markers, ‘modular structure’ and ‘rich-club’ have shaped our understanding of brain networks. The ‘Modular structure’ represents a decomposition of the brain network into clusters of densely connected nodes, whose ties to the other clusters in the network are much sparser, and, as suggested by Rubinov and Sporns (2010), this feature is informative of specialised segregative processes. Traditionally, a modular structure is detected with the Newman Spectral (Newman, 2006) or the Fast Louvain algorithms (Blondel et al., 2008). On the other hand, rich-club is a concept such that the high degree nodes tend to be more densely connected to each other than to the lower degree nodes and it is believed to be informative of complex integrative processes (see, e.g., van den Heuvel and Sporns (2013) for a review). Nevertheless, as demonstrated by Pavlovic et al. (2014) in the brain network of round worm *Caenorhabditis elegans* (*C. elegans*), both aspects of this organisation can simultaneously be captured by a more general concept of ‘cluster structure’, which is defined as the average proportion of edges; within each cluster, and between each cluster pair. This fitted cluster structure typically provides the individual cluster sizes, the total number of clusters *Q* and a *Q* × *Q* matrix of within-cluster and between-cluster connectivity averages ***π***. The variations in mean connectivity values among the cell elements of ***π***, traditionally called ‘blocks’, allow us to characterise the overall network organisation and quantify the portions that are modular and those that are hub-like. In the case of *C. elegans*, the cluster structure turned out to be a combination of 2 rich club clusters, comprising sensory neurons which regulate forward and backward locomotions while the reminder of the clusters were found to be modular. Thus, from a purely methodological view, the cluster structure can be thought of as a rich and elegant tool which marries many isolated graph-theoretical measures, yet remains general enough to capture a catalog of network organisations, including ‘core-periphery’, ‘disassortative’,‘star-patterned’ structures and more (see, e.g., Matias and Robin (2014) for a review), whose occurrence and functional relevances in brain networks are open for discussion.

Beyond *C. elegans*, the analysis of human brain connectivity data is a rapidly growing field of study with a current focus on between-group comparisons between brains of patients and healthy controls in multi-subject datasets (Van Den Heuvel and Pol, 2010). Despite the need for comparative network studies, there has not been much progress in extending single network data analyses of cluster structure to multi-subject data analyses and between-group comparisons. The reasons for this can be found in a set of challenges brought about by the multi-subject nature of the data, including:

i. The need to estimate a common network decomposition over subjects while accounting for between-subject variability in connectivity rates.
ii. How to use such a network decomposition to infer differences between populations (e.g., cases vs. controls) or effects of covariates.

Furthermore, ground truth about cluster structure in human neuroimaging data is generally unknown and assumptions on its form may not be prudent. Hence, there is a pressing need for a statistical framework that provides estimates of cluster labels and general cluster structure, independent from prior assumptions about its form (e.g., assuming that the cluster structure can only be various shades of modular). In line with the multi-subject nature of research on human brain networks, this framework should ideally be rich enough to accommodate for the above mention set of challenges. To address this gap, we follow the class of probabilistic network clustering models originated in the work of Snijders and Nowicki (Nowicki and Snijders, 2001; Snijders and Nowicki, 1997), called Stochastic Blockmodel (SBM). The SBM uses a framework of mixture models to describe heterogeneity in a distribution of network edges. Essentially, one type of distribution is selected for the entire network, such as the Bernoulli distribution, and each cluster and cluster pair has its own Bernoulli parameter. These parameters represent the mean values in the cluster structure ***π*** or, in this specific example, they represent the probability that there is an edge between any two of the nodes which can either be in (a) the same cluster or (b) two different clusters. However, as the clusters can be different sizes, we therefore need a parameter ***α*** to indicate the proportion of each Bernoulli cluster in the overall mixture of distributions. Hence, the larger the cluster, the more likely it is that a randomly selected node falls into it and, thus is expected to contribute more to the mixture.

To obtained the estimates of ***π***, ***α*** and the latent cluster labels, Snijders and Nowicki (1997) considered maximum likelihood estimates (ML) (based on Gibbs sampling and Expectation Maximisation (EM) algorithm) and showcased some computational challenges in the optimisation that limited these techniques to small networks (e.g., < 100 nodes). A trivial change of network density and reparametrisation of the SBM of Snijders and Nowicki (1997) was also considered in the work Newman and Leicht (2007), in which it was referred to it as the ‘Newman and Leicht model’. More recently, Daudin et al. (2008) introduced frequentist variational approximation as an optimisation strategy of Snijders and Nowicki’s SBM, and derived a criterion (Biernacki et al., 1998) that measures the amount of variation in the data explained by a particular network partition. This allowed not only to estimate the optimal number of clusters, but also to quantify the overall goodness of fit of each cluster structure in the dataset. Since then, the SBMs have been adapted for various biological datasets, including the Overlapping SBM (Latouche et al., 2011, 2014), the closely related Mixed-membership SBM (Airoldi et al., 2009), the bayesian SBM (Latouche et al., 2012; Côme and Latouche, 2015), online SBM (Zanghi et al., 2008), the SBM with nodal features (Zanghi et al., 2010) and the SBM with edge features by Mariadassou et al. (2010). SBMs have been applied in the analysis of macaque anatomical cortex (Picard et al., 2009), *C. elegans* brain network (Pavlovic et al., 2014) and human brain networks (Hinne et al., 2015; Moyer et al., 2015; Pavlovic, 2015). Of note, SBM hybrids have also been considered in the analysis of dynamic connectivity (Matias and Miele, 2017) and in neuroimaging (Robinson et al., 2015). Theoretical properties of the SBMs have additionally been studied in the work of Bickel et al. (2013), Choi et al. (2012), Ambroise and Matias (2012), Wolfe and Olhede (2013), Olhede and Wolfe (2014) and Gao et al. (2015) to mention a few.

From this literature, we specifically highlight the work of Mariadassou et al. (2010) who developed the generalised SBM for the analysis of a single network. We refer to this model as generalised SBM for two reasons. First, the variational fitting procedure of SBMs holds for any distribution from the exponential family, which makes it applicable to a wide range of datasets. Second, the authors used generalised linear models to statistically link edge-based features (or covariates) and network cluster structure ***π***. The elegance of this model is that it allows us to study edge-covariate effects, which either affect connectivity in each element of the cluster structure with the same intensity (homogeneous effects), or affect each element of the cluster structure with different intensity (heterogeneous effects). It is also interesting to note that, in both their real data analyses and simulation experiments, the authors only investigated the ‘homogenous effect’ version of their model and, while the ‘heterogenous effect’ model was defined, the behaviour of this model and its domain of application has not been explored.

Inspired by Mariadassou et al. (2010), we consider three multi-subject SBMs (MS-SBMs), Binomial SBM (Bin-SBM), Homogeneous SBM (Hom-SBM) and Heterogenous SBM (Het-SBM). In order to understand the conceptual differences between these multi-subject SBMs, we simulate three simple examples, each of which features three binary and undirected networks with 90 nodes, decomposed into three blocks of equal size. The assignment of nodes is standardised across all subjects, aged 20, 40 and 90 years. The cluster labels of edge probability per each block (i.e. intercept) and age regression coefficients were supplied to the MS-SBMs. Subject-specific connectivity matrices (i.e. per block edge probabilities) were fitted according to each specific MS-SBM. Subsequently, edges for each block according to the Bernoulli density are systematically generated until the entire network is completed. As shown in Figure 1 (a), Bin-SBM assumes common rates of connectivity within each block for all subjects, regardless of age, hence, there is almost no variation in the connectivity between the subjects. For Hom-SBM, the effect of age was set to −0.06 on a logit scale and hence, with increasing age, there is a decrease in connectivity over the entire cluster structure for all subjects (see Figure 1 (b)). Consequently, variability of the cluster structure is tuned to tolerate only minor variations across subjects, as only unidirectional covariate effects can be modelled. In contrast to this, Het-SBM is much more flexible as it allows the effect of age in each block to decrease or increase independently and, thus, different types of cluster structures can be observed across subjects (see Figure 1 (c)). For example, the first subject has a modular structure while the third subject has an overall loss of modular structure in Block 1 as there is an increase of connectivity patterns in Block (1, 2) and (1, 3) which is ‘disassortative’ (Hu and Wang, 2009) (inter-block connections > intra-block connections). In this particular instance, the effect of age in Block (1, 1) was set to −0.06 and in Block (1, 2) to 0.001 on a logit scale, leading to a decrease of connectivity in Block (1, 1) and an increase of connectivity Block (1, 2) with increasing age. Although it is very unlikely that such an extreme variation between subjects is present in real brain data, it is noteworthy that the model is rich and flexible enough to handle this degree of structural variation. For these reasons, we mainly focus on the Het-SBM in this work. As shown in Figure 1, these models assume that the cluster labels are fixed across all subjects but there is an inter-subject variation through subject covariates in a logistic regression model. To our knowledge, this combination of constrained block structure and inter-subject regression model is novel. Unlike in Mariadassou et al. (2010) that utilises edge-based covariates, we propose an entirely new model class that utilises covariates that only vary by subject and not by node or by edge. This is due to the high explanatory power of basic subject covariates, like age and gender (Dosenbach et al., 2010). Furthermore, the inclusion of subject variables allows for inferences between subject populations, such as testing for differences between patient and control groups. In that regard, not much literature about hypothesis testing for multi-subject networks exists. One such work, Alexander-Bloch et al. (2012) discussed how statistical inference can be utilised to compare some global network statistics or some nodal statistics between two groups of networks. However, a significant group difference in a global network statistic does not allow us to locate where the group difference occurs within the multi-subject network. While this shortcoming can be alleviated using nodal statistics, these statistics have a high degree of noise and may yield a large number of tests (at least, one per node). In this paper instead, we utilise our proposed MS-SBMs to make statistical inference on the cluster-wise regression parameters. Such test statistics (1) have a lower level of noise than nodal statistics (due to the pooling effect of the nodes in each cluster), (2) are localised within the network (i.e. by noting the nodes in each involved cluster) and (3) may potentially yield less tests than nodal statistics. Although we do not dismiss the potential value of including edge/node-based covariates (e.g., one could consider a distance-based edge covariate) in our proposed MS-SBMs, we view these to be secondary to the value of an inter-subject regression framework and beyond the scope of the present work. Although we do not dismiss the potential value of including edge/node-based covariates (e.g., one could consider a distance-based edge covariate) in our proposed MS-SBMs, we view these to be secondary to the value of an inter-subject regression framework and beyond the scope of the present work.

**Figure 1:**
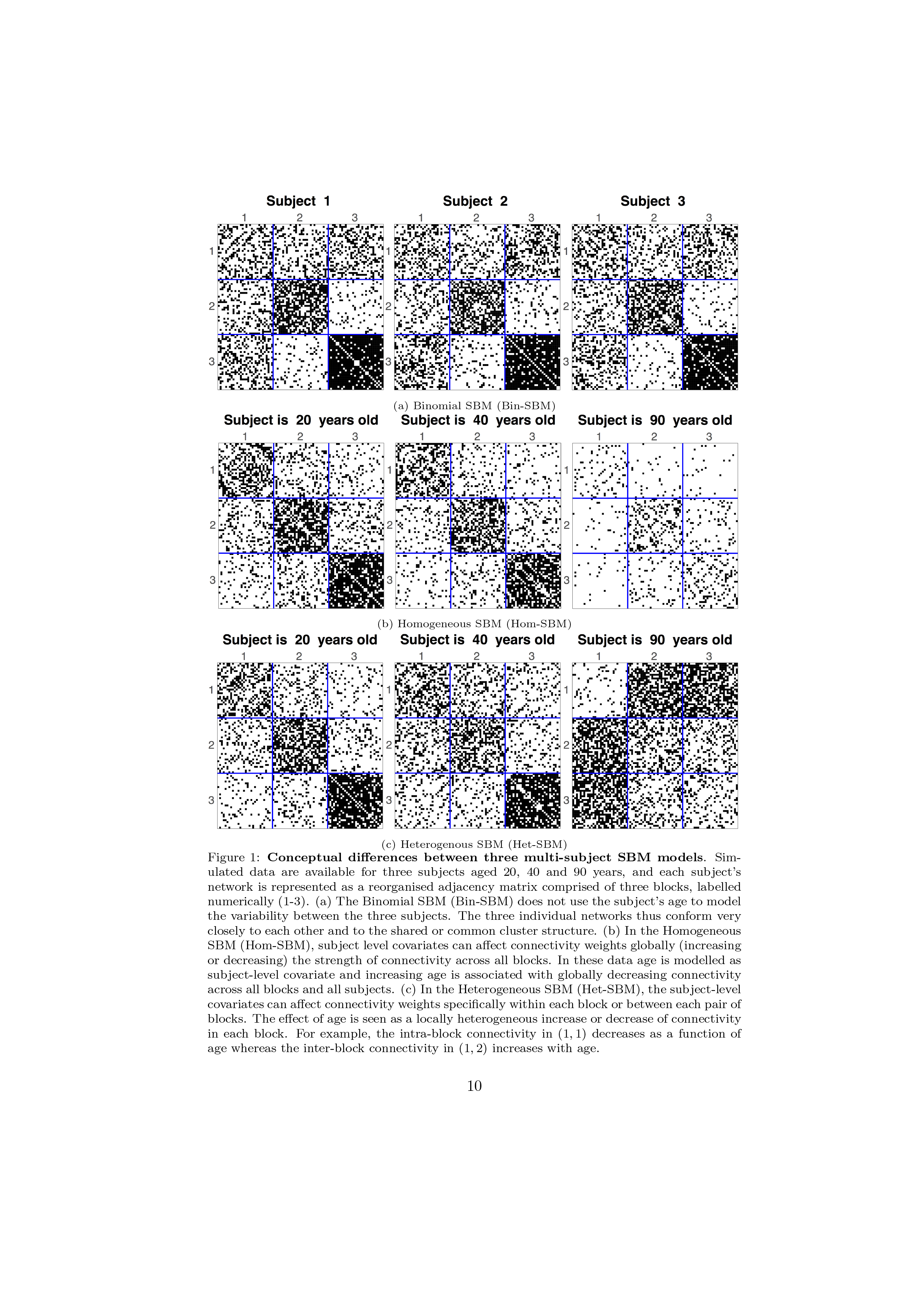
Conceptual differences between three multi-subject SBM models. Simulated data are available for three subjects aged 20, 40 and 90 years, and each subject’s network is represented as a reorganised adjacency matrix comprised of three blocks, labelled numerically (1-3). (a) The Binomial SBM (Bin-SBM) does not use the subject’s age to model the variability between the three subjects. The three individual networks thus conform very closely to each other and to the shared or common cluster structure. (b) In the Homogeneous SBM (Hom-SBM), subject level covariates can affect connectivity weights globally (increasing or decreasing) the strength of connectivity across all blocks. In these data age is modelled as subject-level covariate and increasing age is associated with globally decreasing connectivity across all blocks and all subjects. (c) In the Heterogeneous SBM (Het-SBM), the subject-level covariates can affect connectivity weights specifically within each block or between each pair of blocks. The effect of age is seen as a locally heterogeneous increase or decrease of connectivity in each block. For example, the intra-block connectivity in (1, 1) decreases as a function of age whereas the inter-block connectivity in (1, 2) increases with age.

The remainder of this paper will be organised as follows. First, we will use Bin-SBM to review the variational approximation and the model selection criterion (ICL). Second, we will define Het-SBM for which we discuss an estimation procedure based on a variational approximation and Firth regularisation. Similar derivations for Hom-SBM can be found in Supplemental Information (**SI**) **E**. Third, hypothesis testing (or equivalently inference) procedures based on parametric and non-parametric tests like Wald, likelihood ratio and permutation tests will be described. Fourth, Monte Carlo simulations will be used to evaluate the accuracy of the cluster structure estimated by Bin-SBM and Het-SBM, compared to two prior methods for modular decomposition: Newman’s Spectral algorithm and the Fast Louvain algorithm. As an extra comparison, we will also consider a block-to-node parametrisation of the SBM, given in the work of Newman and Leicht (2007). This SBM will be assessed only on a slice of our simulation because this is a single-network model and as the authors did not implement a strategy to estimate *Q*, we benchmarked this model only on the ground truth *Q*. Fifth, advantages of using Het-SBM over Bin-SBM will be shown, and an example in which the between-subject variation strongly impacts the estimated cluster structure and cluster labels will be raised. Sixth, using simulations, we will assess the performance of parametric and non-parametric tests in terms of their control of false positives (Type I error). Seventh and lastly, we will apply Het-SBM to a resting state fMRI study with 25 subjects split into two groups: healthy subjects (controls) and subjects diagnosed with schizophrenia (cases).

## 2. Methods

In this section, we first setup the preliminary notation, and then we describe the MS-SBMs followed by their associated hypothesis testing procedures based on the Wald, likelihood ratio and permutation tests.

### 2.1 Notation

We will be employing the usual statistical convention of capital Roman letters to denote random variables and lower case letters to denote their observed realisations. Scalar and non-scalar values are denoted by light and bold face fonts, respectively. For example, a non-scalar random variable will be denoted as ***X*** and its realisation as ***x***. If ***X*** is a discrete random variable, *f*_***X***_(***x***) ≡ *f* (***x***) ≡ P(***X*** = ***x***) are three equivalent notations for its probability mass function or discrete density. Densities conditional on a random variable are written using a vertical bar, as in *f*(***x***|***z***). Furthermore, we will be using the notation 𝔼_*f*\*_ [*g*(*X*)] with *f*\* indicating the density of *X*. That is, 𝔼_*f*\*_ [*g*(*X*)] = Σ_*x*∊*S*_*g*(*x*)*f*\* (*x*), where *S* is the sample space of *X*. As shown in Eq. (5), the notation is utilised to establish the variational nature of the underlying distribution.

In both models, the random variable *X*_*ijk*_ represents the possibility of an edge between the nodes *V*_*i*_ and *V*_*j*_ in the *k*-th subject, and *x*_*ijk*_ denotes a binary realisation of *X*_*ijk*_ with 1 being an edge and 0 no edge. Therefore, for the *k*-th subject, ***X***_*k*_ = ((*X*_*ijk*_))_1≤*i*≠*j*≤*n*_ denotes an *n* × *n* random, symmetric adjacency matrix with elements *X*_*ijk*_, and, for a total of *K* subjects, ***X*** denotes the set of independently distributed random matrices ***X*** = {***X***_1_, …, ***X***_*K*_}. The individual matrices ***x***_*k*_ = ((*x_ijk_*))_1≤*i*≠*j*≤*n*_ are assumed to be undirected, without self-connected nodes and without multiple-edges between the nodes. Hence, they are binary and symmetric matrices with 0s on their principal diagonal and a total of *n*(*n* − 1)/2 data points.

The goal of each model is to estimate a common cluster structure among *K* subjects. Thus, both multi-subject models assume that the set of nodes, labelled as {*V*_1_, …, *V*_*n*_}, are divided into *Q* unknown (latent) blocks or clusters. Block membership of a particular node *V*_*i*_, will be indicated by a random vector ***Z***_*i*_ = (*Z*_*i*1_, …, *Z*_*iQ*_) whose elements *Z*_*iq*_ take the value 1 if *V*_*i*_ ∈ *q*-th group and 0 otherwise. Pooling this information across nodes, the *n* × *Q* random matrix ***Z*** can be defined such that the vectors ***Z***_*i*_ are mutually independent and follow a categorical density with *Q* possible outcomes

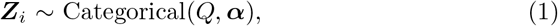

where ***α*** is a 1 × *Q* dimensional vector of success probabilities ***α*** = (*α*_1_, …, *α*_*Q*_) and 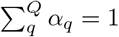. Specifically, if we assume that the cluster labels for each node are given, we interpret an individual *α*_*q*_ as the probability that a randomly selected node falls into the *q*-th block. Here, it is important to note that the choice of categorical (or equivalently single trial multinomial) density implies that fitted blocks form a partition of all nodes, in which each node belongs to only one block (i.e. disjoint blocks). This is formally noted as 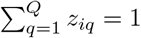.

### 2.2 Binomial Stochastic Blockmodel (Bin-SBM)

In this model, all subjects are assumed to have the same variation, such that the distribution of edges is not influenced by subject-specific information (covariates) but instead is solely dependent on observed edges. Consequently, for the *k*-th subject, the edges are assumed to follow a Bernoulli distribution

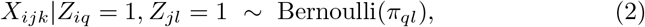

where *π*_*ql*_ is the connectivity rate that expresses the probability that nodes are connected between (*q, l*)-th blocks. For all blocks, connectivity rates are compiled into a *Q* × *Q* matrix ***π*** wherein the within-block rates are located on the diagonal, and between-block rates are located on the off-diagonal of this matrix. As the edges are undirected, the ***π*** matrix is symmetrical (i.e. *π*_*ql*_ = *π*_*lq*_). In particular, each block-specific connectivity rate *π*_*ql*_ is the expected mean of its edges (i.e. *π*_*ql*_ = 𝔼(*X*_*ij*_|*Z*_*iq*_ = 1, *Z*_*jl*_ = 1)).Thus, by parameterising each block component separately, the Bin-SBM can represent a host of different network topologies, including various degrees of modular organisation and core-periphery structures.

Block-specific connectivity rates are assumed to be constant across all subjects, suggesting that random variables 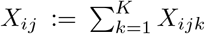 follow a binomial distribution,

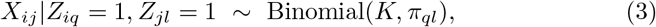

where realisations across subjects can be compiled into a symmetric connectivity matrix, denoted by the random variable ***X*** = ((*X*_*ij*_))_1≤*i*≠*j*≤*n*_. Notably, this trivial model matches the case of binomially distributed edges in a single network in Mariadassou et al. (2010), where edges between node pairs represent the total number of observed edges across *K* subjects.

#### Estimation

Closely following Daudin et al. (2008) and Mariadassou et al. (2010), the framework of variational approximation is utilised to estimate the model’s parameters. The standard approach formulates the statistical model in terms of *complete data*, represented by the joint discrete density *f* (***x***, ***z***; ***π***, ***α***) whose likelihood is given as

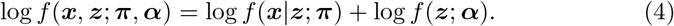

The marginal of the complete data likelihood is noted as a *incomplete data* likelihood, log *f* (***x***; ***π***, ***α***) and has the same parameters as the complete data likelihood. Ideally, the estimation of the model’s parameters should be based on *f* (***x***; ***π***, ***α***). However, as the explicit calculation of this density is computationally challenging, we use a variational approach.

In the variational approach, the model’s parameters are estimated by optimising the variational bound 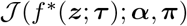, defined as

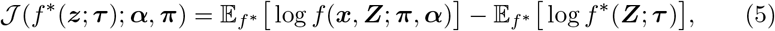

where *f*\*(***z***; ***τ***) depends on the variational parameter ***τ*** (*n* × *Q* matrix of posterior probabilities), and it denotes the parametric family that is closest (in the Kullback-Leibler sense) to *f* (***z***|***x***; ***π***, ***α***). The complete derivation of this bound and its relationship to *f* (***x***; ***π***, ***α***) can be found in **SI** A. Based on Eq. (5), computation of the variational bound requires taking expectations of ***Z*** with respect to their variational density *f*\*(***z***; ***τ***). To make this computationally feasible, *f*\*(***z***; ***τ***) is taken to be a product of individual densities of ***Z***_*i*_. Each density is categorical with block-specific probabilities, that are independent in each node

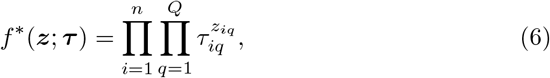

where Σ_*q*=1_ *τ*_*iq*_ = 1, and 𝔼_*f*_* (*Z*_*iq*_) = *τ*_*iq*_, 𝔼_*f*_* (*Z*_*iq*_*Z*_*jl*_) = *τ*_*iq*_*τ*_*jl*_. In particular, *τ*_*iq*_ is the strength of evidence that a node *V*_*i*_ is a member of block *q* having observed the data. For example, the node *V*_*i*_ in a three cluster structure has a vector 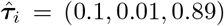. Based on the ‘maximum a posteriori’ (MAP) estimate shown in Eq. (11), this node has the strongest affiliation to the third block as its posterior probability is the highest (i.e. 0.89).

Taking the expectations stated in Eq. (5), the variational bound can more concisely be written as

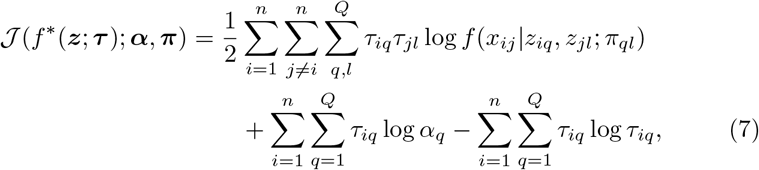

where 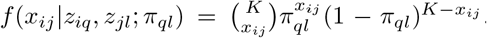. Optimising Eq. (7) with respect to the variational parameter ***τ*** and the model’s parameters ***α*** and ***π***, according to the constraints 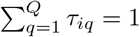 and 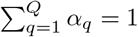, we find the following fixed point relations

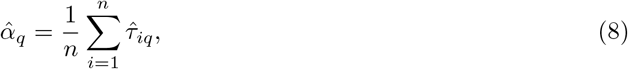

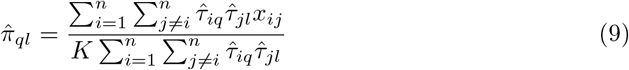

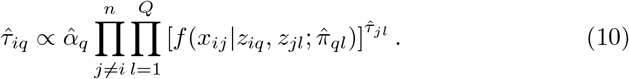

Detailed derivations of the above mentioned equations can be found in **SI** B. An estimate of the probability that a randomly selected node is assigned to the *q*-th block 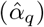 in Eq. (8) is calculated as the average of posterior probabilities of all nodes in block *q*. An estimate of within- or between-block connection probability 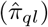 in Eq. (9) is calculated as the expected number of edges between block *q* and *l*, taking into account the posterior probability of nodes belonging to block *q* and *l*. The estimate of 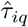 (Eq. (10)) is proportional to (1) the estimate of the node belonging to the *q*-th block 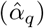 and (2) the probability of observing connectivity of node 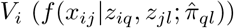, taking into account posterior block assignment probabilities of other nodes 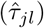.

Finally, estimates of the classification vector 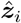 are obtained from 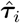 such that

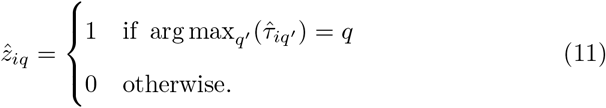

#### Model Selection

To estimate an optimal model with *Q* clusters, denoted by m_*Q*_, we use the Integrated Classification Likelihood (ICL) criterion proposed by Biernacki et al. (1998) and its adaptation presented in Daudin et al. (2008), (see **SI** C for details). The notation m_*Q*_ refers to all the parameter estimates of Bin-SBM 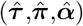 for *Q* clusters. The ICL criterion is defined on a complete data log likelihood (see Eq. (4)) such that

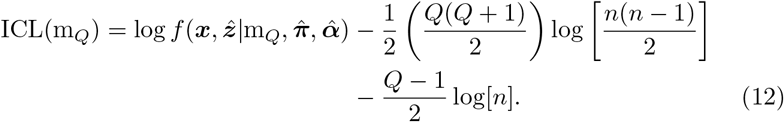

Thus, ICL serves as a measure of goodness of model fit, mediating the trade off between the fitted complete data likelihood (first term in Eq. (12)) and the complexity of the model (second and third terms in Eq. (12)). As *Q* increase, the second and the third terms in Eq. (12) progressively penalise the overall ICL scores with increasing severity. For example, a network with 100 nodes and 3 clusters, has a penalisation term of 30, but, if the number of clusters is increased to 30, the penalisation term increases to 2045. Selecting a model with 30 clusters over a model with 3 clusters can only be justified when the difference between their complete data likelihoods is large enough to outweigh the heavy penalisation term.

#### Implementation Details

Beginning with the initial values for ***τ***^(0)^ (see **SI** D.2 for details), model parameters are updated in two iterative steps:

1. 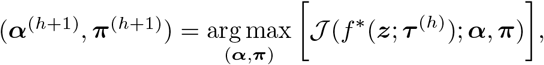
2. 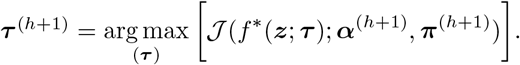

First, in the Maximisation Step or (M-Step), we use Eq. (8) and Eq. (9) to update ***α*** and ***π***, respectively. Second, in the Expectation Step (or E-Step), we use Eq. (10) to update ***τ***. Both steps are iterated until parameter estimates reach convergence. Technical details of the algorithm and a pseudocode can be found in **SI** D.2.

As shown in Daudin et al. (2008), this fitting procedure ensures that the algorithm iteratively climbs the variational bound (Eq. (7)), which in turn maximises the incomplete data log likelihood log *f* (***x***; ***α***, ***π***) without the need to calculate it directly. This guarantees that the algorithm will converge at a local maximum typically in the neighbourhood of the initial estimate ***τ***^(0)^, but it does not guarantee convergence at the global maximum; identical limitations are found in the classical EM-algorithm (Wu, 1983). It is worth noting that, while selecting a local maximum solution may not be ideal, the algorithm should improve upon the starting point, obtained through other clustering algorithms (unless, of course, the starting point is already a local maximum). Hence, the procedure may be expected to provide an improvement upon the fit of any other clustering algorithms.

For a given network dataset, the likelihood surface can be heavily spiked, and, therefore, an informative starting point should yield a better likelihood fit than a non-informative one. The quality of parameter estimates depends on how informative ***τ*** is. Therefore, instances in which many nodes can be equally assigned to all clusters will tend to produce poor estimates of ***π***, yielding a very small ICL score. However, selecting an informative starting point is often very difficult in practice (Scrucca and Raftery, 2015) as the large number of possible choices (*Q*^*n*^) make it impossible to explore every possible starting point. In our application, hierarchical clustering (Murtagh and Legendre, 2014) was utilised to generate an initial estimate of ***τ***^0^ as it was found to perform relatively well in practice and had been used successfully by Mariadassou et al. (2010). Nevertheless, other approaches may also be suitable, and, thus, depending on time constraints and dimensionality of the data (i.e. the number of nodes in the studied network), it is possible to consider a random initialisation whereby the nodes labels are sampled from *Q* number of blocks. In this particular instance, one would also need to consider multiple initialisations so that the sample space of all possible node classification is searched more effectively. Finally, it is worth mentioning that similar issues are found in many graphical models using variational algorithms (e.g., Mixed Membership SBM, Overlapping SBM, Latent Dirichlet Association model, Hidden Markov models and more).

### 2.3 Homogeneous Stochastic Blockmodel (Hom-SBM)

In this model, variability between subjects is assumed to be a global feature of the multi-subject networks. Thus, conditional on its node assignments, each edge is assumed to follow a Bernoulli distribution with a probability of connection as a function of the subject covariates via a logistic regression model

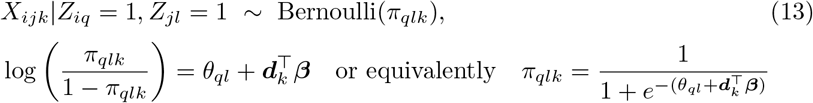

where *π*_*qlk*_ is the *k*-th subject’s connectivity rate in block (*q, l*), and *θ*_*ql*_ is the common intercept in block (*q, l*) across all subjects (i.e. baseline block probability on the logit scale). In particular, 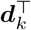 is the *k*-th subject’s 1 × *P* dimensional vector of covariates and ***β*** is a *P* × 1 vector of regression coefficients. This model has a block specific intercept for all subjects (*θ*_*ql*_), and covariates that relate globally to the overall block structure but not individual blocks.

In contrast to Het-SBM, where each covariate was allowed to interact with the cluster structure in a block-wise manner (see Figure 1 (c)), the covariates in Hom-SBM only engage with the entire cluster structure in a global fashion (see Figure 1 (b)). This suggests a smaller degree of variation in the cluster structure across subjects in Hom-SBM compared to Het-SBM. Hom-SBM may be useful in neuroimaging applications (e.g. accounting for a global nuisance effect during preprocessing), and we have provide a detailed derivation of the estimation and inference procedures of in **SI** E. However, Het-SBM is preferable for the purposes of testing varying covariate effects which may increase the number of connections between some brain areas, but decrease it between other brain areas. Thus, in this paper, we have chosen to focus on Het-SBM.

### 2.4 Heterogenous Stochastic Blockmodel (Het-SBM)

In this model, variability between subjects is assumed to influence within- and between-block connectivity. Thus, conditional on its node assignments, each edge is assumed to follow a Bernoulli density, whose rates of connection depend on the subject covariates via a logistic regression model. Formally,

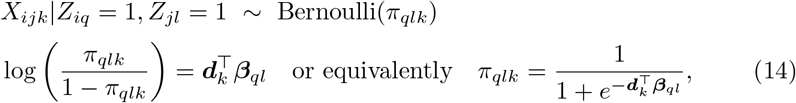

where *π*_*qlk*_ is the connectivity of block (*q, l*) associated with subject *k*, 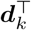 is the 1 × *P* vector of covariates associated with subject *k* (typically the first element will be 1, representing the intercept), and ***β***_*ql*_ is a *P* × 1 vector of regression parameters. The *s*-th component of ***d***_*k*_ and ***β***_*ql*_ are denoted as ***d***_*ks*_ and ***β***_*qls*_, respectively, while the total of *Q*(*Q* + 1)/2 individual regression vectors ***β***_*ql*_ are collectively denoted as ***β***.

#### Estimation

In this model, the variational optimisation strategy and Fisher’s scoring algorithm are used to estimate the parameters ***τ***, ***α*** and ***β***. When estimating ***β***, circumstances may arise where the inferences about ***β*** may be biased due to small or sparse samples (i.e. rare events^1^) or due to a perfect matching between covariate and binary scores (i.e. complete separation^2^). To prevent such biases, a Firth type estimation (Firth, 1993; Kosmidis, 2014) is used.

Similar to the estimation strategy used for the Bin-SBM (see Section 2.2), we begin by defining the lower bound. Since ***β***_*ql*_ = ***β***_*lq*_, we use the notation *γ*_*ijql*_ to denote the following cases

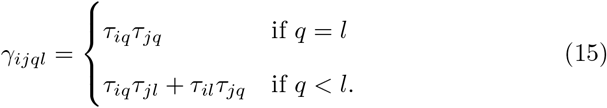

Thus, the variational bound can be stated as

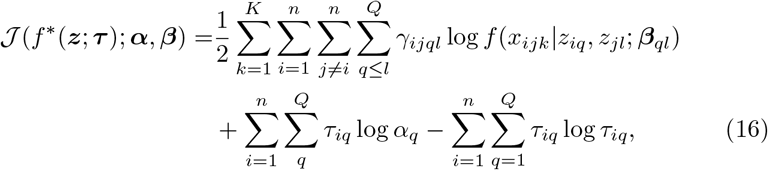

where 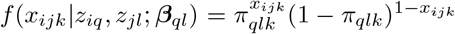 and *π*_*qlk*_ depends on ***β***_*ql*_ (Eq. (14)). Optimising Eq. (16) for ***τ*** and ***α*** yields the following point estimating equations:

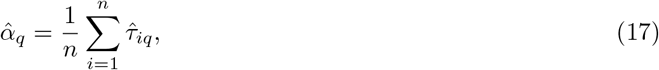

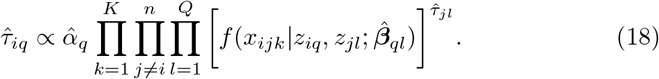

The Firth type estimates for ***β***_*ql*_ are generated from optimising the modified variational bound 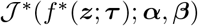 such that

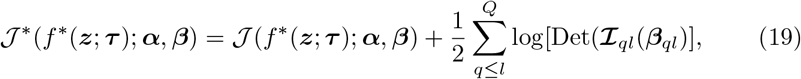

where 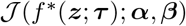 is given in Eq. (16) and the term 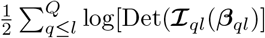 is a penalisation term. The first order partial derivates of 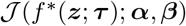 with respect to ***β***_*ql*_ can be written as **U**_*ql*_(***β***_*ql*_) such that

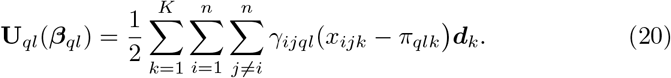

Similarly, negative second order partial derivatives with respect to ***β***_*ql*_ yield a Fisher Information matrix 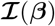, expressed as a *Q*(*Q* + 1)/2*P* × *Q*(*Q* + 1)/2*P* block diagonal matrix of individual sub-matrices 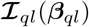. This can be defined as

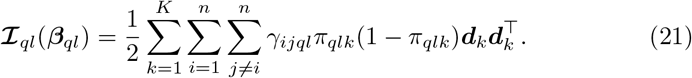

To find estimates of ***β***_*ql*_, the modified bound given in Eq. (19) is optimised according to Fisher’s scoring formula

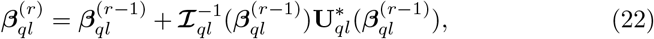

where (*r*) is the *r*-th iteration and 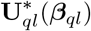 is the modified score vector. The *s*-th element of 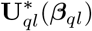 is

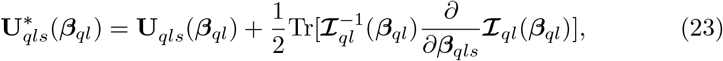

where

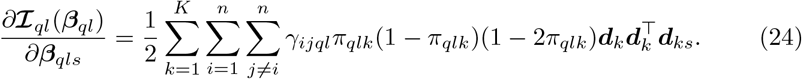

#### Model Selection

Similar to Bin-SBM, the ICL criterion for Het-SBM is given as

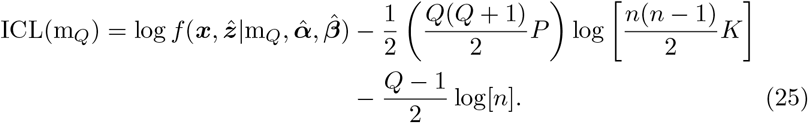

#### Implementation Details

Beginning with some initial values for ***τ***^0^, the model parameters are iteratively updated according to a two-step procedure

1. 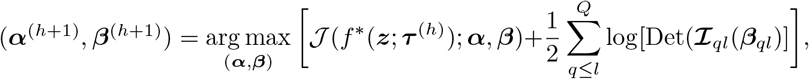
2. 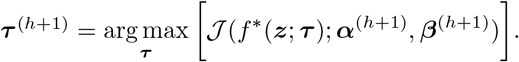

In the first step, Eq. (17) and (22) are used to update ***α*** and ***β***, respectively. The part of the variational bound related to ***β*** is calculated by the sum of products between the non-negative weights ***γ*** and the conditional log likelihoods of the logistic regression model (see the first term in Eq. (16)). As the conditional log likelihood of the logistic regression is globally concave, and the individual components of ***γ*** are non-negative, then the sum of their product is also globally concave. This implies that there is at most one global maximum and, if the global maximum exists, the algorithm will converge on it. However, in extreme cases of complete or quasi separation, the maximum likelihood solution will not exist. In this situation, the algorithm will typically return high values for the regression coefficients but their variances would diverge to infinity, rendering inferences inaccurate. However, an optimisation with Firth penalisation typically ensures the existence of a global maximum, high quality estimates of regression coefficients and a numerically stable algorithm even in samples with rare events and extreme circumstances of separation. The latter property was confirmed by Heinze and Schemper (2002) who wrote the package `logistf` in `R` software which implements Firth regression for binary data. In the well-behaved datasets (i.e. without any of the above mentioned peculiarities), there is a perfect agreement between the maximum likelihood and Firth estimates. Fisher’s scoring algorithm is initialised with zero as the starting values for the parameters (***β***). For each iteration of Fisher’s scoring algorithm, a maximal absolute change of value 5 is set for all the parameters (e.g., if the change for a parameter is estimated to be −7.2, it is forced to be −5). This ensures that changes in the parameters are not too large, which might cause the algorithm to exceed the maximum. When the weighted log likelihoods (i.e. block-wise log likelihoods) are smaller than values obtained with previous parameter estimates, a step-halving procedure is executed. Parameter updates are reduced by half until an improvement is observed or until a maximal number of halving steps is reached, wherein the previous parameter estimates are retained. This strategy is consistent with the implementation in `logistf`.

In the second step, ***τ*** is updated according to Eq. (18). These two steps are iterated until the convergence is obtained in terms of the relative changes of the parameter estimates. The pseudocode for these algorithms can be found in **SI** F.

In this context, several strategies can be used for parametric (i.e. Wald and likelihood ratio test) or non-parametric (i.e. permutation test) inference conditional on ***τ*** and ***α*** (i.e. the block assignments) such that sampling distributions of the variational estimator coincide with the maximum likelihood estimator (MLE).

### 2.5 Inference in Multi-subject Stochastic Blockmodels

Multi-subject models estimate a common cluster structure across subjects, serving as the common ground for making comparisons between subjects. Since Het-SBM imposes a logistic regression model on each element of the cluster structure, the methodological framework of logistic regression can be used to estimate differences between groups of subjects or the effects of covariates on the connectivity rate at each block. While these could simply be used as network summary metrics, the logistic regression model offers the additional possibility to determine if linear combinations of these quantities are statistically different from a specified constant (e.g., 0). In this context, several strategies can be used for parametric (i.e. Wald and likelihood ratio test) or non-parametric (i.e. permutation test) inferences, and each such test is assumed to be conditional on ***τ***. Due to the joint optimisation in the variational algorithm of Het-SBM, cluster labels estimates depend on the covariates in the logistic regression model. Once the model with the highest clustering evidence is fitted, inferences are carried out without accounting for the variance of ***τ***. This notably ensures the interpretability of results for non-parametric tests as it prevents scenarios where the permuted clusters do not overlap with the original cluster.

#### Wald test

The Wald test has commonly been used in logistic regression analysis to make inferences on the estimates of regression coefficients. In the context of Het-SBM, due to special block diagonal structure of the Fisher Information matrix, the null hypothesis can be written as 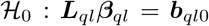. The Wald statistic takes the form

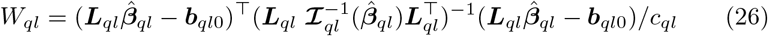

where ***L***_*ql*_ is a matrix (or a vector) that defines the combination of the parameters (or contrast) tested and *c*_*ql*_ denotes the rank of ***L***_*ql*_. Asymptotically, *W*_*ql*_ follows a 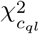 distribution. If ***L***_*ql*_ is a vector, then

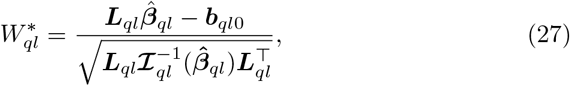

asymptotically follows a Standard Normal distribution. As performed in both the Ordinary and Forth MLE approaches (Firth, 1993; Heinze and Schemper, 2002), standard errors of model parameters are estimated with the Fisher Information matrix (see Eq. (21)).

#### Likelihood ratio (LR) test

An alternative to the Wald test for the inference on the combination of parameters 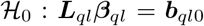 is the likelihood ratio (LR) test. Conceptually, the likelihood ratio test compares the full (or alternative) model against the restricted (or null) model. Thus, the likelihood ratio statistic is formulated as

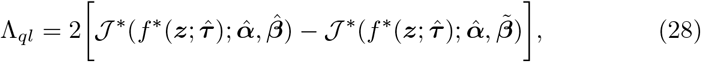

where 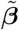 is the set of all parameters estimated under the null hypothesis. More precisely, 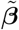 is estimated from the modified variational bound of the restricted model which has a different penalisation term from the one of the full model. While 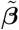 is obtained using the modified variational bound of the null model, it is substituted into the modified variational bound of the full model to finally get the likelihood ratio statistic.

Similarly to the Wald test, under the null hypothesis, Λ_*ql*_ is assumed to follow a 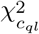 distribution, where *c*_*ql*_ is the rank of ***L***_*ql*_. Note that for Ordinary MLEs, the likelihood ratio is based on the non-modified variational bound given by Eq. (16).

#### Multiple testing

Inference procedures for the block level parameters effectively make *Q*(*Q* + 1)/2 individual tests and this presents multiple comparison problem. To control the family-wise error rate (FWE), defined as the probability of making at least one Type I error, the Bonferroni correction (Holm, 1979) is utilised. This correction is valid for any dependence structure and is easy to apply: Instead of using a nominal *α*_0_ significance value (e.g., 0.05), *α*_0_/*n*_*T*_ is used instead, where *n*_*T*_ is the number of tests (here, *n*_*T*_ = *Q*(*Q* + 1)/2).

#### Permutation test

In addition to Wald and likelihood ratio (LR) tests, that depend on asymptotic sampling distributions, permutation tests are also considered (Good, 2000). Permutation tests are based on the premise that, under the null hypothesis, the data can be exchanged without altering its distribution. This implies that the distribution of any test statistic can be found empirically through repeated evaluations of randomly rearranged (or permuted) data. In the context of Het-SBM, permutation tests are used to make inferences on the parameter vector ***β***_*ql*_ under the null hypothesis, in which there is no association between edge occurrence and the covariate tested. In our simulations, only tests of the entire parameter vector ***β***_*ql*_ are considered. However, it is important to note that such an approach would not be possible in real fMRI as the model depends on more than one covariate. To account for this, a simple permutation strategy proposed by Potter (2005) is used. In this approach, the covariate of interest is regressed on the remaining nuisance covariates (using a linear regression model), and the residuals from this model are used in place of the original covariate. Following the permutation of these residuals, logistic regressions are fitted and *P*-values are computed by comparing the resulting test statistics against their original scores.

The *P*-value of the observed Wald test statistic *w*_*ql*0_ is computed by a Monte Carlo sampling scheme such that for a sequence of random permutations indexed by *t* (*t* = 1, …, *M*), there is *M* Wald statistics labelled as *w*_*ql*1_ …, *w*_*qlM*_. Including the observed Wald statistic *w*_*ql*0_ in the permuted scores, the Monte Carlo *P*-value is computed as

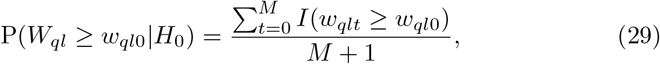

where *I*(·) is the indicator function. The same approach can be used with the likelihood ratio test to obtain *P*-values without assuming an asymptotic *χ*^2^ distribution.

#### Improved Multiple Testing Procedures with Permutation

The Bonferroni method for controlling the FWE is conservative in the presence of dependence. Thus, there is a need to consider an alternative permutation based procedure (Westfall and Young, 1993). For the Wald statistic *w*_*ql*0_, the FWE corrected *P*-value is given as

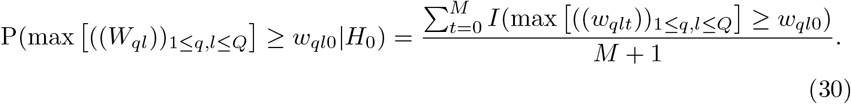

The FWE corrected *P*-values for the likelihood ratio (LR) statistics are computed in the same way.

## 3. Methodology of Simulations and Results

This section details the methodology and results of three sets of simulations labelled *Simulation I-III*. To make the cross-referencing between initial simulation parameters and results more convenient for the reader, this section first details the goals and setups of each simulation, and then their corresponding results.

### 3.1 Simulation I Goals and Setup

#### Goal of Simulation I

The goal of *Simulation I* is to assess the overall performance of Bin-SBM and Het-SBM in a range of different cluster structures which exhibit important deviations from a classically modular organisation^3^. The between subject variations in the cluster structures are assumed to be independent from subject covariates. Since this setup does not consider covariate effects, it allows for the comparison between multi-subject SBMs and modular decomposition methods such as the Fast Louvain (FL) and Newman Spectral (NS) algorithms. The modular algorithms are utilised in terms of their stand alone implementations as well as in combination with consensus clustering.

#### Setup of Simulation I

In this simulation, the total number of blocks, subjects and nodes has been fixed to 10, 30 and 200, respectively (i.e. *Q* = 10, *K* = 30 and *n* = 200). The range of block sizes was determined according to three designs, which we will refer to as Balanced, Mildly Unbalanced (M. Unbalanced) and Unbalanced designs. Each design is characterised by varying degrees of heterogeneity of block sizes, detailed in Table 1. Given the selected network size, a *Q* of 10 was selected in order to yield similar block sizes to those already reported in applied literature (Picard et al., 2009). While one could also vary *n* and *Q*, and perhaps even study entirely different experimental questions like small sample behaviours (Pavlovic, 2015), our prior experience with different *n* and *Q* have been found to yield comparable outcomes to what was obtained.

**Table 1:** Block sizes for three block size designs. In the Balanced design, all blocks have the same size while, in the Mildly Unbalanced (M. Unbalanced) and Unbalanced designs, there is a systematic decrease in the overall number of nodes such that the last blocks are the smallest. In the M. Unbalanced design, the block sizes are mildly varying while, in the Unbalanced case, the block sizes are changing more rapidly.

| Design Label | Block Labels |  |  |  |  |  |  |  |  |  |
| --- | --- | --- | --- | --- | --- | --- | --- | --- | --- | --- |
|  | 1 | 2 | 3 | 4 | 5 | 6 | 7 | 8 | 9 | 10 |
| Balanced | 20 | 20 | 20 | 20 | 20 | 20 | 20 | 20 | 20 | 20 |
| M. Unbalanced | 38 | 32 | 27 | 23 | 19 | 16 | 14 | 12 | 10 | 9 |
| Unbalanced | 60 | 41 | 29 | 20 | 15 | 11 | 8 | 6 | 5 | 5 |

The connectivity rates ***π*** have been chosen according to three types of cluster structures: *Homogeneous-Modular* (Hom-Modular), *Heterogeneous-Modular* (Het-Modular) and *Core-Modular*. As shown in Figure 2, within-block and between-block connectivity rates in the Hom-Modular cluster structure have constant values in both the principal diagonal and off-diagonal parts of the matrix. In contrast, these values vary randomly in the Het-Modular structure. Both connectivity matrices have larger connectivity rates on the principal diagonal than on the off-diagonal elements of the matrix and, as such, they fit into a general definition of modular network structure. Finally, the Core-Modular structure in Figure 2 is a combination between the heterogeneous modular structure (Blocks 1-8) and a densely inter- and intra-connected core (Blocks 9 & 10). The first two cluster structures (Hom-Modular and Het-Modular) were chosen because of the extensive evidence that brain networks demonstrate at least some degree of modularity (Meunier et al., 2009) across many different species, including the *C. elegans*, macaque and human brains (Bullmore and Sporns, 2009; Rubinov and Sporns, 2010; Pan et al., 2010; Sporns et al., 2004; Hilgetag et al., 2000; Achard et al., 2006; Bassett and Bullmore, 2009). However, there is also strong evidence that the cluster structure of a brain also consists of a core of rich club nodes. Previous work on Stochastic Blockmodel analysis of *C. elegans* connectome (Pavlovic et al., 2014) revealed a hybrid structure or ‘core-on-modules’ structure which deviated from a purely modular organisation by the inclusion of densely inter- and intra-connected core blocks. This kind of structure was captured by the Core-Modular structure in Figure 2, where two cores (Blocks 9 &10) are densely connected to all other blocks in an otherwise heterogeneous modular network.

**Figure 2:**
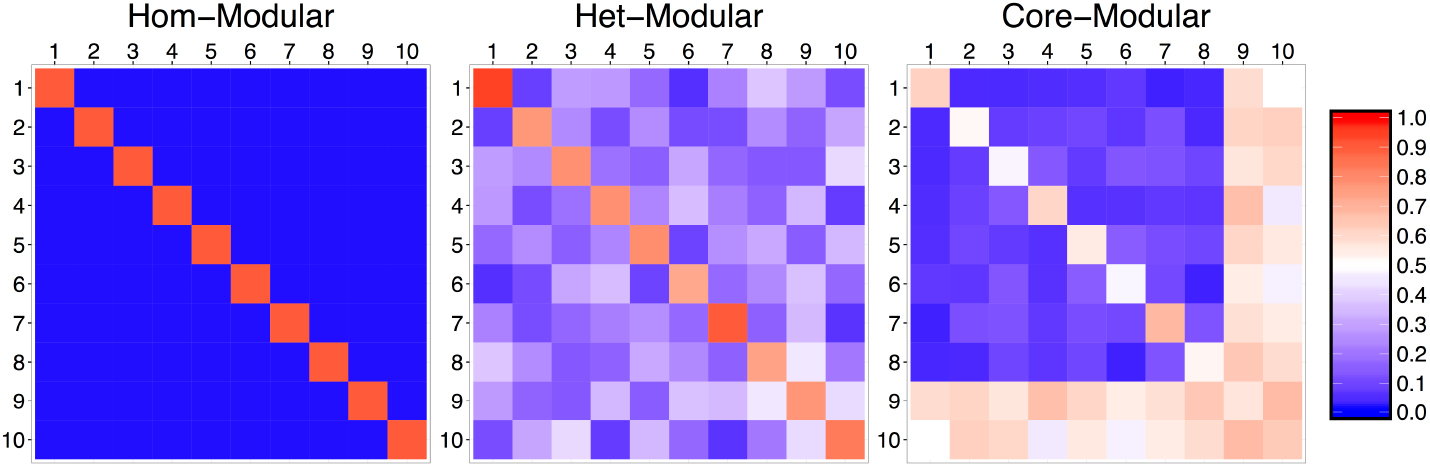
Connectivity matrices (*π*) for three different cluster structures. Each cluster structure is characterised by differing degrees of modular organisation, defined as a relatively higher within-blocks connectivity rates compared to the between-blocks connectivity rates. The homogeneous modular (Hom-Modular) case represents perfect, classical modularity characterised by high within-block and low between-block connectivity rates that are common across all blocks. The heterogeneous modular (Het-Modular) case retains the modular structure but it allows for varying rates of connectivity within and between different blocks. The Core-Modular structure is a hybrid organisation characterised by (a) a heterogenous modular structure (Blocks 1-8) and (b) two core blocks with dense within- and between block connections (Block 9 & 10).

For each of these three cluster structures (i.e. Hom-Modular, Het-Modular and Core-Modular) and for each of the block designs (Balanced, Mildly Unbalanced and Unbalanced), we generate Bernoulli realisations according to the connectivity rates in Figure 2 in order to systematically build a network. For example, to generate a network with a Hom-Modular structure where Block 1 has 20 nodes and a within-block connectivity rate of 0.9, we perform (20 × 19/2: maximum number of edges in Block 1) Bernoulli trials where the probability of seeing an edge (i.e. outcome 1) is 0.9. Similarly, in order to generate connections between 20 nodes in Block 1 and 20 nodes in Block 2, we perform (20 × 20: maximum number of edges between Block 1 & 2) Bernoulli trials using the probability specified by the connection rate between these two blocks (e.g., 0.05). In this block-wise manner, an entire network can be systematically built, and 100 individual networks for each of these combination of parameters (i.e. block design and cluster structure) can be generated. It is important to note that firstly, the block assignment of each node is already known and, therefore, the ground truth can be used to benchmark the ability of a model to correctly estimate true cluster assignments. Secondly, our data generating process does not take into account between-subject variation.

Each of the simulated multi-subject networks are fitted with Bin-SBM, Het-SBM, the Newman Spectral (NS) and the Fast Louvain (FL) algorithms to study their abilities to retrieve the true clustering. Specific details of each fitting procedure are given below.

#### Fitting procedure for MS-SBMs

For a given network and each value of 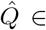 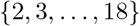 blocks, Bin-SBM and Het-SBM are fitted 30 times with different initialisations obtained from k-means, random label sampling and hierarchical clustering (hclust). For k-means, 10 initialisations are generated using the function `Kmeans` (from the R package `amap` (Lucas, 2014)) on the average adjacency matrix, which allowed the restarts to vary through to the stochastic nature of k-means. For random label sampling, 10 starting points are generated through uniform sampling of labels between one and 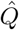. For hclust, which is deterministic in nature, 10 initialisations are generated by applying the `hclust` function (R package `stats` (R Core Team, 2017)) on the average adjacency matrix, and on nine randomly selected subject-specific adjacency matrices. The best solution is selected as the one with the highest ICL score both with respect to different initialisations and values of *Q*.

#### Fitting procedure for NS-A and FL-A

The Newman Spectral (NS) and Fast Louvain (FL) algorithms are run on the average adjacency matrices with 510 restarts^4^. The corresponding results are referred to as NS-A and FL-A where the first two letters indicate the algorithm and the last letter indicates that the method is applied on the averaged data. For the Fast Louvain algorithm, the same 510 initialisations used by Bin-SBM and Het-SBM are utilised. No specific initialisation is used for the Newman Spectral algorithm as the function `modularity_und` does not allow the setting of the initialisation. Each of these modular algorithms is used with 11 different values as their resolution parameter *γ* ranging from 0.5 to 1.5 with a step size of 0.1, to show the influence of the tuning parameter on their fits. The best solution with respect to each resolution parameter value is selected as the one with the highest modularity score. Unfortunately, the modularity function heavily depends on the value of the resolution parameter *γ* so that the larger the value of *γ*, the larger the modularity score is and, as such, it is not possible to use it to compare the fits across different values of *γ*.The functions for the Newman Spectral and Fast Louvain algorithms can be found respectively as `modularity_und` and `community_louvain` in the `Matlab Brain Connectivity Toolbox` (Rubinov and Sporns, 2010, http://www.brain-connectivity-toolbox.net/, last accessed 01 May 2018).

#### Fitting procedure for NS-C and FL-C

A consensus strategy is also used for each of the two modular algorithms (Lancichinetti and Fortunato, 2012). The corresponding results are referred to as NS-C and FL-C where the first two letters indicate the algorithms and the last letter indicates that the method is applied with a consensus algorithm. Within this approach, each clustering algorithm is run 10 times on each subject-specific adjacency matrix to compute a within-subject agreement matrix, that indicates the number of times each node pair was assigned to the same cluster. Each of the subject-specific agreement matrices is then normalised by dividing it by its maximum value, and further filtered by setting all values smaller than an arbitrary cutoff value to zero (here, 0.8 as suggested by one of the reviewers). Each of the resulting matrices is then fitted by the given clustering algorithm to yield a within-subject consensus clustering for each subject. Finally, a between-subject agreement matrix of within-subject consensus clustering assignments is computed. It is also divided by its maximum value and filtered using the same cutoff value of 0.8 as mentioned above. The given clustering algorithm is then used on the filtered between-subject agreement matrix to obtain the final assignment of nodes. This consensus strategy is repeated several times with 11 different values as their resolution parameter *γ* ranging from 0.5 to 1.5 with a step size of 0.1. In total, 22 versions of each modular algorithm are fitted. The best solution with respect to each resolution parameter value is selected as the one with the highest modularity score.

#### Benchmarking Methodology

For this simulation, we evaluate each approach by the ability to (a) estimate the optimal number of clusters (i.e. *Q* = 10) and (b) achieve the true clustering. This is assessed by the Adjusted Rand Index (ARI) (Handl et al., 2005; Hubert and Arabie, 1985, see **SI** G, for details) which indicates the overall agreement between selected best fits and true clustering.

### 3.2 Simulation I Results

Table 2 reports, for Het-SBM in the Core-Modular scenarios, the ranges over the 100 Monte Carlo samples of the best adjusted ICL scores, each defined as the difference between the maximum ICL score for a given 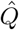 and the maximum ICL score over all 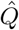. The range [0, 0] indicates that, for all 100 Monte Carlo samples, the best ICL score systematically points towards the true number of clusters (i.e. 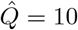). The ICL scores tend to decrease when 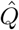 is taking values which are going further away from the ground truth 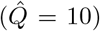 with a greater rate for values below 10. The same behaviour has been observed for all the other scenarios as well as for Bin-SBM (see **SI** H; Tables S1-S2 for Het-SBM and Tables S3-S5 for Bin-SBM), indicating that both models are reliable in retrieving the true number of clusters.

**Table 2:** Ranges of adjusted ICL scores for Het-SBM in the scenarios with Core-Modular cluster structure. For each simulated dataset and a given 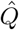, the adjusted ICL scores are given as the difference between the maximum score across the restarts (local maximum) and all values of 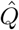 (global maximum). For each 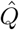, the range of ICL scores is shown across all 100 Monte Carlo samples. The range [0, 0] indicates that, for all 100 Monte Carlo samples, the best ICL score systematically points towards the correct number of clusters (i.e. 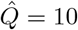). ICL scores tend to decrease when 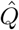 utilise values that increasingly diverge from the ground truth 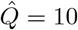 with a greater rate for values below 10.

| $\hat{Q}$ | Core-Modular | | |
| --- | --- | --- | --- |
|  | Balanced | M. Unbalanced | Unbalanced |
| 2 | [-31252, -30193] | [-44884, -43403] | [-51276, -49308] |
| 3 | [-23621, -22803] | [-24265, -23200] | [-22118, -21019] |
| 4 | [-18689, -17173] | [-16489, -15498] | [-10824, -10095] |
| 5 | [-14118, -12602] | [-10843, -10036] | [-6742, -5866] |
| 6 | [-9936, -8991] | [-6912, -6318] | [-3058, -2704] |
| 7 | [-6296, -5649] | [-4536, -3566] | [-1827, -1547] |
| 8 | [-3365, -2986] | [-2331, -1992] | [-843, -675] |
| 9 | [-974, -763] | [-520, -335] | [-628, -140] |
| 10 | [0, 0] | [0, 0] | [0, 0] |
| 11 | [-76, -63] | [-77, -63] | [-77, -62] |
| 12 | [-164, -137] | [-156, -134] | [-157, -139] |
| 13 | [-243, -215] | [-238, -215] | [-244, -221] |
| 14 | [-327, -299] | [-328, -300] | [-334, -315] |
| 15 | [-429, -389] | [-427, -395] | [-432, -407] |
| 16 | [-521, -479] | [-531, -496] | [-540, -513] |
| 17 | [-633, -594] | [-641, -602] | [-656, -607] |
| 18 | [-739, -703] | [-758, -704] | [-770, -728] |

Figure 3 shows the distribution of estimated 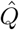 for all approaches. Each modular fit is given in terms of the three *γ* values (i.e. *γ* ∈ {0.5, 1, 1.5}). The full set of results can be found in **SI** H; see Figures S1, S3 and S5. Consistent with the above mentioned results, Bin-SBM and Het-SBM accurately estimate the optimal number of clusters in all simulation scenarios. In contrast, modular algorithms show a variation in their estimates, and tend to depend heavily on the block designs and cluster structures. In particular, both algorithms are accurate in the Hom-Modular case with the Balanced and M. Unbalanced designs, where there is minimal influence from the tuning parameter (*γ*) and, minimal difference between average and consensus clustering. However, in the case of Hom-Modular with Unbalanced design, there is a strong influence of *γ* in the average clustering but not in consensus clustering. This suggests that consensus clustering is robust in cases with small clusters, and perfectly modular networks. Unlike Hom-Modular, all Het-Modular and Core-Modular cases exhibit great dependence on *γ*. For the Fast Louvain algorithm, there is a tendency to underestimate *Q* for small values of *γ* and to overestimate *Q* for large values of *γ*. In contrast, the Newman Spectral algorithm exhibits a tendency to underestimate *Q* for all the range of *γ* values investigated. It is interesting to note that the values of *γ* that yield the most accurate results are strongly varied across block designs and cluster structures, indicating a potential difficulty in setting its value in real data applications. Another noteworthy point is that, although the Fast Louvain algorithm had the same initialisations as the MS-SBMs, it obtained very different estimates of the optimal number of clusters.

**Figure 3:**
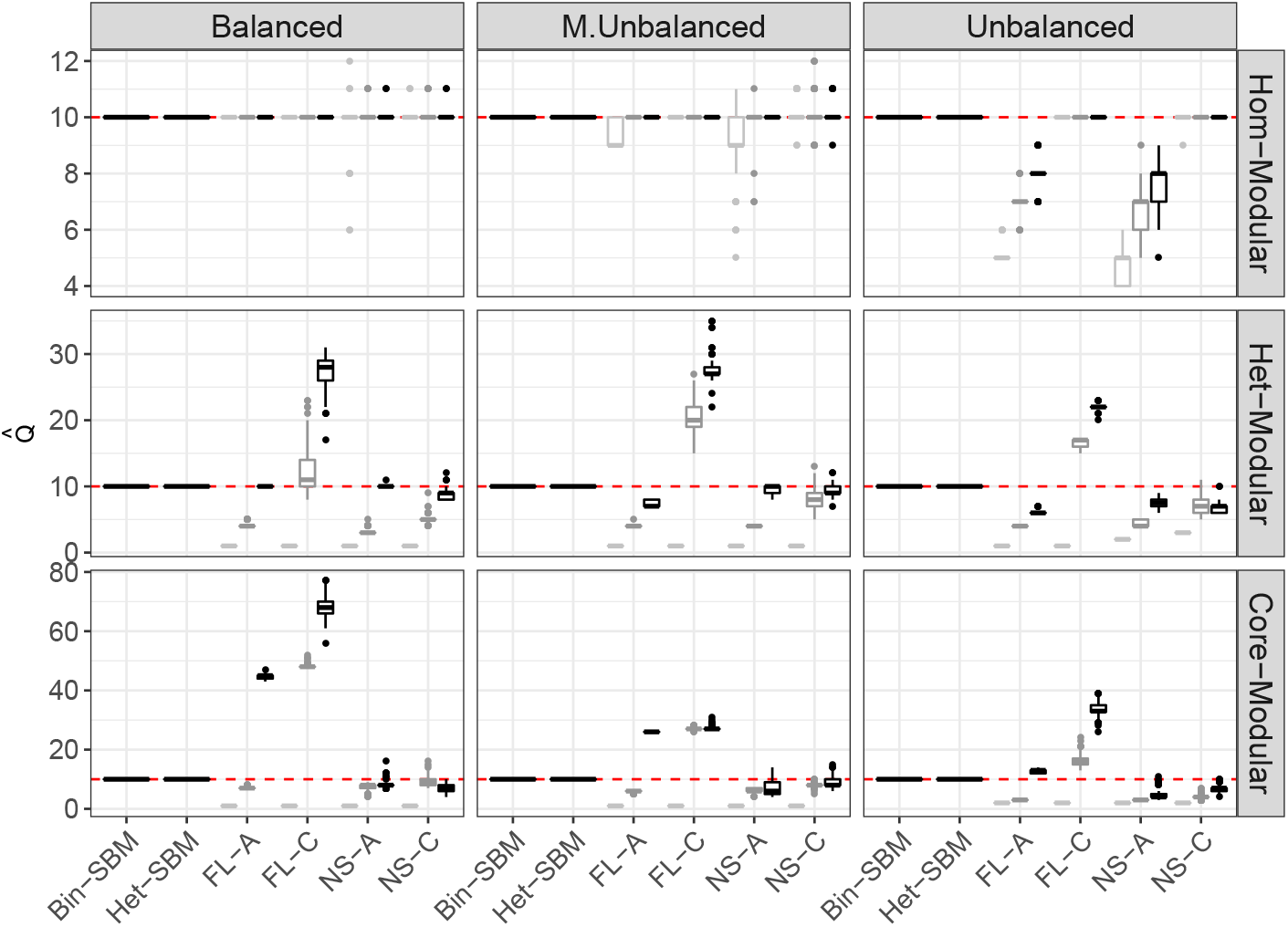
Box-plots of estimated 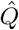 over 100 Monte Carlo samples in all simulation scenarios. The *x*-axis shows MS-SBMs and the modular algorithms such that: FL-A and FL-C stands for the Fast Louvain algorithm with averaged data and consensus clustering, and NS-A and NS-C stands for the Newman Spectral algorithm with averaged data and consensus clustering. For each modular algorithms, box-plot colours correspond to solutions with different *γ* values (i.e. *γ* ∈ 0.5, 1, 1.5, with a central score of 1 (default value). Ground truth is indicated by the horizontal red line plotted at 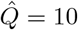. In brief, Bin-SBM and Het-SBM consistently estimate the ground truth *Q* while the modular algorithms tend to be unreliable in many scenarios.

In Figure 4, box-plots represent the Adjusted Rand Index (ARI) scores (ARI ∈ [0, 1]; see **SI** G) over 100 Monte Carlo samples, with 1 denoting a perfect agreement and 0 a complete disagreement. For space considerations, only three different values of the tuning parameter (i.e. *γ* ∈ {0.5, 1, 1.5}; see **SI** H Figures S2, S4 & S6 for the complete results) are shown for each of the modular fits. Bin-SBM and Het-SBM exhibit high accuracy in all simulation scenarios. In contrast, the modular algorithms show difficulties when retrieving true cluster labels, especially in the Het-Modular and Core-Modular cases. The ARI scores appear to be strongly influenced by the tuning parameter *γ*, are highest when *γ* is large or, in a few cases (e.g., FL-C in the Balanced Core-Modular scenario), around 1. The most surprising aspect of this analysis is that some fits with very high ARI scores (close to 1) strongly overestimate the total number of clusters. The most notable example of this can be seen for FL-A and FL-C (*γ* = 1.5) in the Core-Modular and M. Unbalanced design (see Figures 3 and 4). The reason for this is that the majority of the clusters are correctly retrieved while the few remaining clusters are entirely split into single-node clusters. This type of misclassification is not well captured by ARI scores as they stay relatively close to 1 even for non-parsimonious partitions, which is highly undesirable. A potential remedy is to consider the ICL criterion (see Eq. (25)) given the fitted partition 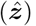. In this simulation, each fitted partition was assigned an ICL score, and the results across 100 Monte Carlo datasets have been summarised in Table 3. To make a comparison across all approaches, a range of adjusted ICL scores for the Core-Modular scenarios was computed as the difference between (1) ICL scores of each approach, and (2) highest ICL score across all approaches. The fits of Bin-SBM and Het-SBM, as shown in Figure 4, perfectly recover the true partitions in all designs, yielding a [0, 0] range. However, the modular algorithms were not found to yield optimal partitions. For example, despite ARI values close to 1 in the M. Unbalanced case (see Figure 4), both the fit of FL-A and the fit of FL-C (*γ* = 1.5) resulted in ranges of [−6454, −5275], and [−2822, −2040], respectively. Similar results were obtained in the Het-Modular cases while, for the Hom-Modular cases, the fits were fairly accurate (see **SI** H Figures S6 and S7).

**Table 3:**
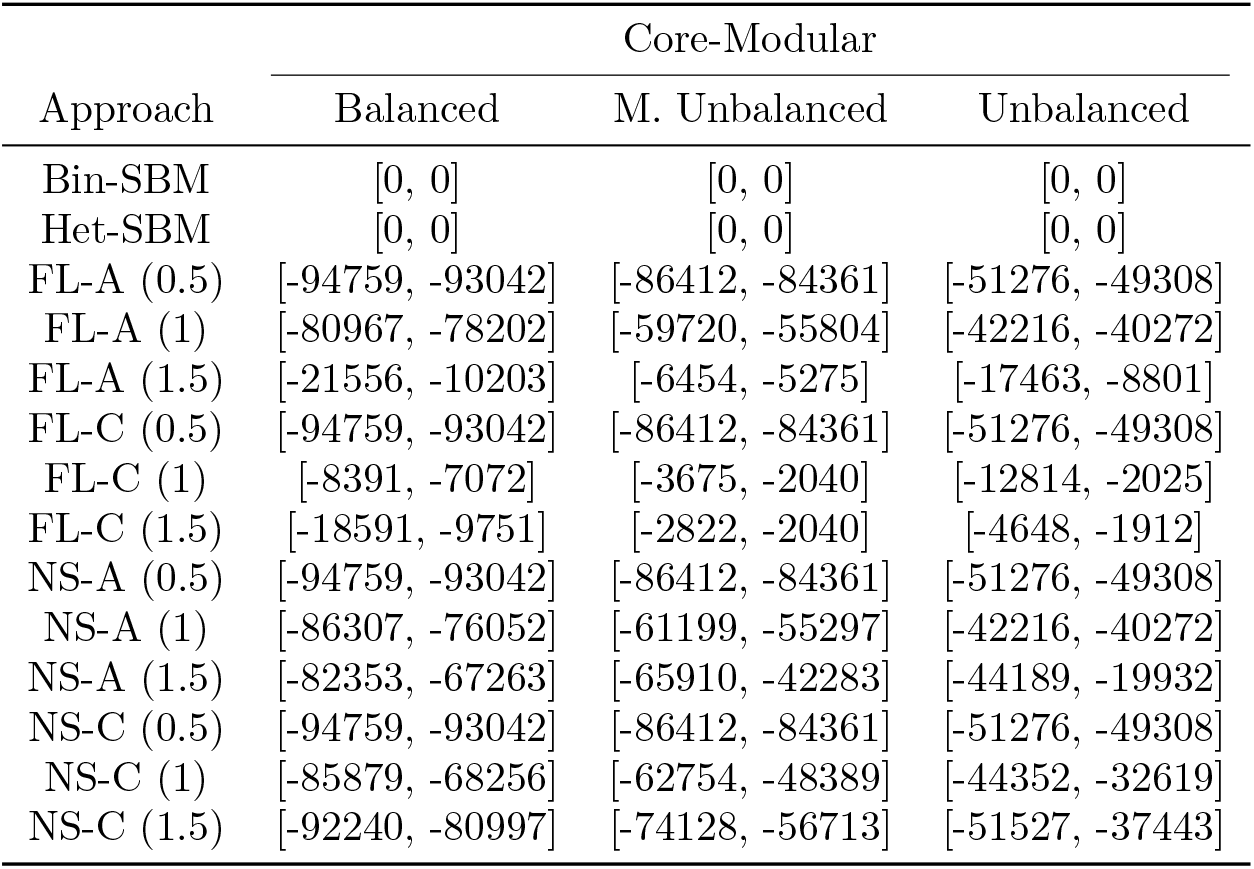
Ranges of adjusted ICL score of the best fits for several approaches in the scenarios with Core-Modular cluster structure. For each simulated dataset and approach, the adjusted ICL score of the best fit is the difference between its ICL score and the maximum ICL score across all approaches. For each approach, the range of ICL scores is shown across all 100 Monte Carlo samples. In brief, Bin-SBM and Het-SBM systematically yielded the highest ICL scores.

| Approach | Core-Modular |  |  |
| --- | --- | --- | --- |
|  | Balanced | M. Unbalanced | Unbalanced |
| Bin-SBM | [0, 0] | [0, 0] | [0, 0] |
| Het-SBM | [0, 0] | [0, 0] | [0, 0] |
| FL-A (0.5) | [-94759, -93042] | [-86412, -84361] | [-51276, -49308] |
| FL-A (1) | [-80967, -78202] | [-59720, -55804] | [-42216, -40272] |
| FL-A (1.5) | [-21556, -10203] | [-6454, -5275] | [-17463, -8801] |
| FL-C (0.5) | [-94759, -93042] | [-86412, -84361] | [-51276, -49308] |
| FL-C (1) | [-8391, -7072] | [-3675, -2040] | [-12814, -2025] |
| FL-C (1.5) | [-18591, -9751] | [-2822, -2040] | [-4648, -1912] |
| NS-A (0.5) | [-94759, -93042] | [-86412, -84361] | [-51276, -49308] |
| NS-A (1) | [-86307, -76052] | [-61199, -55297] | [-42216, -40272] |
| NS-A (1.5) | [-82353, -67263] | [-65910, -42283] | [-44189, -19932] |
| NS-C (0.5) | [-94759, -93042] | [-86412, -84361] | [-51276, -49308] |
| NS-C (1) | [-85879, -68256] | [-62754, -48389] | [-44352, -32619] |
| NS-C (1.5) | [-92240, -80997] | [-74128, -56713] | [-51527, -37443] |

**Figure 4:**
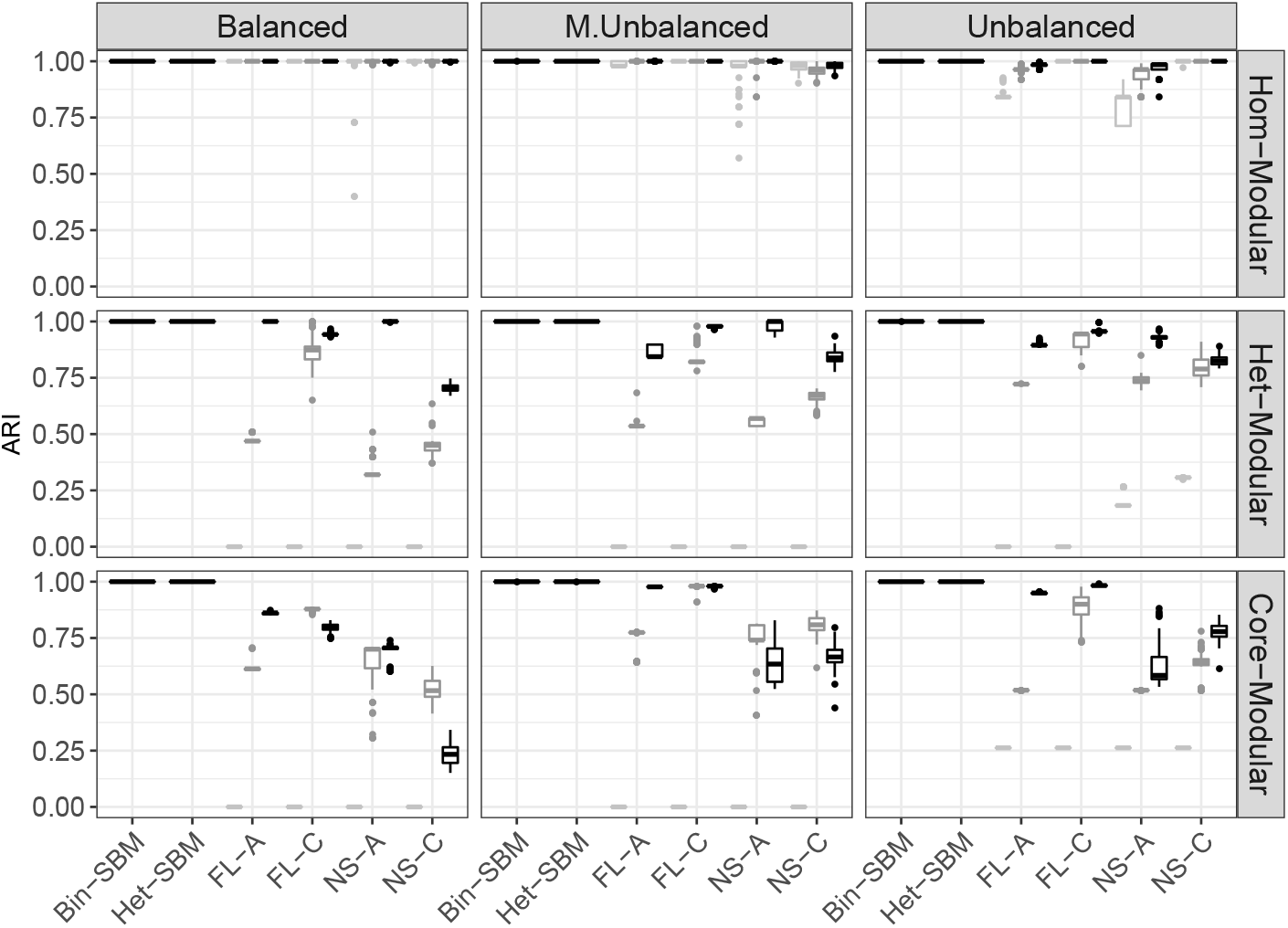
Boxplots showing the recovery of the true cluster assignments over 100 Monte Carlo samples in terms of ARI scores. FL and NS stand for Fast Louvain and Newman Spectral algorithms while -A and -C indicate the strategy to use the algorithm on the average adjacency matrix and the consensus strategy, respectively. For space considerations, only the results with *γ* = 0.5 (light grey), 1 (mild grey) and 1.5 (black) are shown for the modular algorithms (see **SI** H Figures S2, S4 and S6 for the full results).

### 3.3 Simulation II Goals and Setup

#### Goals of Simulation II

The goal of *Simulation II* is to explore situations in which covariate effects pose a strong influence on the estimation of cluster labels. Although our data has been simulated according to fixed cluster labels for all subjects, it is still possible to encounter situations in which the joint optimisation of cluster labels and logistic regression covariates strongly informs the final clustering fit. In this simulation, the fit based on Bin-SBM portrays an example in which covariates are discounted from the estimation of cluster labels. For example, this would be a case in a which a researcher initially fits cluster labels independent of the logistic regression model, and then subsequently uses the clusters to fit a logistic regression model. At first glance, this may appear to be a viable approximation that reduces the computational burden from nested logistic regression optimisation. However, there are situations in which a collection of nodes with fairly similar connectivities show evidence supporting further clustering when fitted with informative covariates.

#### Setup of Simulation II

In this simulation, the parameters related to the total number of blocks, subjects and nodes were set to the same values as in *Simulation I* (i.e. Q = 10, K = 30 and n = 200). Similarly, the block sizes varied according to the Balanced, M. Unbalanced and Unbalanced designs (see Table 1). The subjects are divided into two equal groups. The first 15 subjects have individual 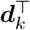 set to (+1, +1) (group 1) while the remaining 15 subjects have individual 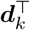 set to (+1, −1) (group 2). The first entry corresponds to a common intercept representing the average effect of both groups while the second entry corresponds to a differential effect between the first group and the second group. Figure 5 (a) and (b) show the block-specific common intercept and group effect coefficients, respectively. All 30 subjects share a common Core-Modular cluster structure with a densely integrated core comprising Blocks 1 & 2, and a heterogeneous set of modules comprising Blocks 3-10. Group effects only interact with this cluster structure of core blocks. To account for this, the regression coefficients for core blocks Block 1 & 2 were set to a range of values on a scale between +2.50 and −2.50. Differences between the two groups will only be apparent in the core blocks and not in the modular components of the cluster structure. This scenario can be expected in the clinical setting where it has been found that neurological disorders preferentially impact high degree hubs that constitute the core or rich club of the cluster structure (Crossley et al., 2014).

**Figure 5:**
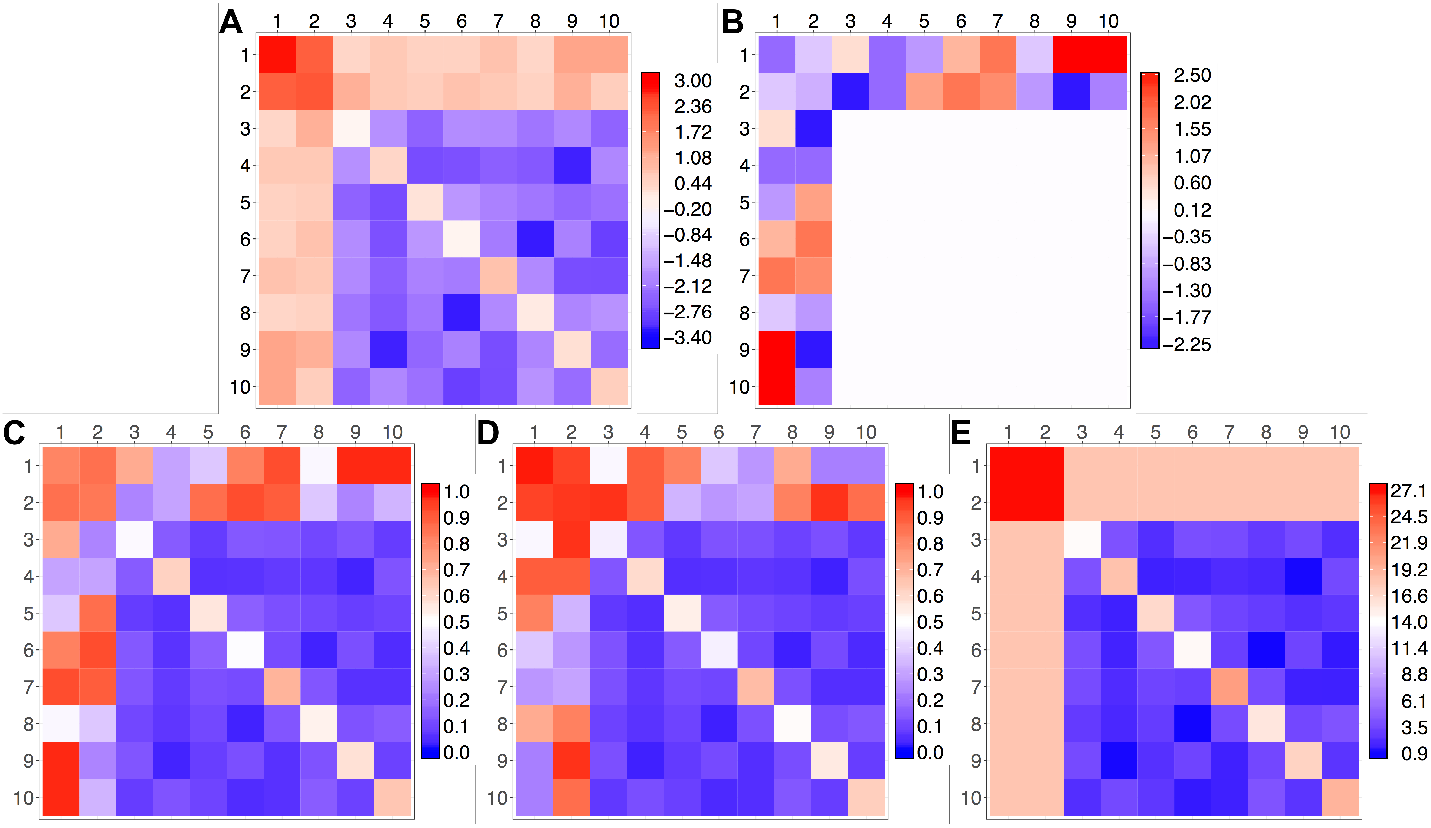
Simulation of between-group (control-case) differences in cluster structures. **A**: Block specific connectivity rates were common to all subjects and specified by intercept ((*β*_*ql*1_))_*q,l*∈{1,…,*Q*}_. **B**: Differential effect between first and second groups ((*β*_*ql*2_))_*q,l*∈{1,…,*Q*}_. Intercept scores clearly delineate a tightly integrated core comprised of Blocks 1 & 2, and a heterogeneous modular structure comprised from Blocks 3-10. Differential effect between the two groups is sampled on a scale between +2.50 and −2.50 for the two core blocks. For the remaining blocks, differential was set to 0.05. Thus, the control-case differences in cluster structure are represented specifically as deviations from intra-block and inter-block connectivity of a common Core-Modular cluster structure. **C**: Cluster structure of subject 11 in the first group. **D**: Cluster structure of subject 23 in the second group. **E**: Sum of connectivity matrices across two groups. While individual subjects have clearly delineated cluster structure with 10 blocks (**C** & **D**), the overall average of connectivity matrices shows only the evidence of 9 blocks, as Block 1 & Block 2 are merged together (**E**).

#### Fitting and Benchmarking

As noted earlier in Eq. (14), the intercepts and group coefficients in Figure 5 (a) and (b) are utilised to simulate (100 Monte Carlo samples) of individual connectivity matrices for each subject in each group. Subsequently, Bin-SBM, Het-SBM and modularity algorithms are fitted according to the procedures described in Simulation I. The accuracy of the estimated cluster structure is evaluated by ARI scores (see **SI G)**.

### 3.4 Simulation II Results

For Bin-SBM and Het-SBM, Table 4 and 5 report the ranges over the 100 Monte Carlo samples of the best adjusted ICL scores, each defined as the difference between the maximum ICL score for a given 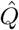 and the maximum ICL score over all 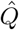. The range [0, 0] gives an estimation of the total number of clusters for all 100 Monte Carlo samples. In Table 4, Bin-SBM systematically under-estimates the true number of clusters across all three block designs, suggesting 9 clusters instead of 10. In contrast to this, the results in Table 5 show that Het-SBM accurately estimates the true number of clusters across the Balanced and M. Unbalanced designs in all the simulations. In the Unbalanced design, Het-SBM estimates the true number of clusters in 90% of the simulations, but overestimates it by one cluster in the remaining 10% of simulations. This behaviour might be explained by the fact that, in some simulations, the algorithm may not have converged to the global maximum.

**Table 4:** Ranges of adjusted ICL scores for Bin-SBM. For each simulated dataset and a given 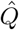, the adjusted ICL scores are given as the difference between the maximum score across the restarts (local maximum) and all values of 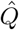 (global maximum). For each 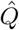, the range of ICL scores is shown across all 100 Monte Carlo samples. The range [0, 0] indicates that, for all 100 Monte Carlo samples, the best ICL score systematically underestimates optimal number of clusters (i.e. 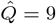).

| $\hat{Q}$ | Balanced | M. Unbalanced | Unbalanced |
| --- | --- | --- | --- |
| 2 | [-28480, -27138] | [-18727, -17732] | [-11519, -10847] |
| 3 | [-21206, -20078] | [-12998, -12049] | [-7678, -6280] |
| 4 | [-16318, -15356] | [-8938, -8318] | [-3482, -3152] |
| 5 | [-12402, -11101] | [-6500, -5443] | [-2080, -1800] |
| 6 | [-8487, -7671] | [-4303, -3379] | [-1269, -1058] |
| 7 | [-5618, -4785] | [-2039, -1778] | [-552, -390] |
| 8 | [-2371, -2121] | [-1074, -685] | [-330, -139] |
| 9 | [0, 0] | [0, 0] | [0, 0] |
| 10 | [-77, -44] | [-99, -46] | [-122, -47] |
| 11 | [-141, -92] | [-184, -107] | [-218, -105] |
| 12 | [-198, -148] | [-261, -158] | [-304, -183] |
| 13 | [-254, -212] | [-342, -212] | [-387, -269] |
| 14 | [-311, -278] | [-418, -284] | [-464, -334] |
| 15 | [-379, -342] | [-469, -354] | [-551, -400] |
| 16 | [-455, -423] | [-540, -429] | [-652, -464] |
| 17 | [-536, -496] | [-637, -510] | [-747, -536] |
| 18 | [-612, -578] | [-670, -592] | [-784, -605] |

**Table 5:** Ranges of adjusted ICL scores for Het-SBM. For each simulated dataset and a given 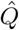, the adjusted ICL scores are given as the difference between the maximum score across the restarts (local maximum) and all values of 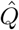 (global maximum). For each 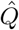, the range of ICL scores is shown across all 100 Monte Carlo samples. The range [0, 0] indicates that, for all 100 Monte Carlo samples in the Balanced and M. Unbalanced designs, the best ICL score systematically points towards the correct number of clusters (i.e. 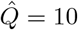). In the Unbalanced designs, the model accurately estimates optimal number of clusters in 90% of simulations but in 10% it overestimates the optimal number of clusters by one.

| $\hat{Q}$ | Balanced | M. Unbalanced | Unbalanced |
| --- | --- | --- | --- |
| 2 | [-66095, -64206] | [-68990, -67233] | [-63637, -62204] |
| 3 | [-49671, -48018] | [-49121, -46608] | [-45079, -43575] |
| 4 | [-37345, -35761] | [-36239, -34848] | [-32554, -31715] |
| 5 | [-25638, -23925] | [-22773, -21361] | [-17800, -17117] |
| 6 | [-18602, -17267] | [-13319, -12652] | [-8661, -8114] |
| 7 | [-13371, -12119] | [-9187, -7252] | [-3931, -3497] |
| 8 | [-8358, -7784] | [-3732, -3298] | [-1423, -1106] |
| 9 | [-4275, -3843] | [-1326, -1129] | [-1463, -228] |
| 10 | [0, 0] | [0, 0] | [-421, 0] |
| 11 | [-137, -126] | [-149, -127] | [-149, 0] |
| 12 | [-292, -266] | [-311, -274] | [-313, -147] |
| 13 | [-451, -424] | [-487, -426] | [-491, -313] |
| 14 | [-629, -598] | [-676, -603] | [-680, -512] |
| 15 | [-820, -779] | [-878, -783] | [-883, -681] |
| 16 | [-1021, -985] | [-1075, -974] | [-1093, -908] |
| 17 | [-1244, -1197] | [-1319, -1185] | [-1322, -1126] |
| 18 | [-1470, -1424] | [-1543, -1412] | [-1564, -1360] |

The differences in results between Bin-SBM and Het-SBM are not surprising since the simulated data contains a clear group effect (see Figure 5 (c) and (d), in which each group shows a strong evidence for 10 clusters). However, by assuming that the estimation of cluster labels is independent from the group effect, information about the 10 cluster structure is lost and smoothed out into a cluster structure with 9 connectivity profiles (see Figure 5 (d)). As a consequence of this, Bin-SBM fails to correctly estimate the optimal number of clusters by one cluster in all simulations and design cases. This example illustrates a potential issue that can arise when separating the clustering procedure from the logistic regression model and shows the benefit of using a joint optimisation of the cluster labels and the regression model. As we can see in the case of the Bin-SBM fit, this may lead to an inaccurate clustering, which then poses further problems for post-hoc analysis, like fitting a regression model or doing inferences.

The distribution of the total number of clusters across all approaches is shown in Figure 6. The estimates of 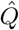 for Bin-SBM and Het-SBM are consistent with the results shown in Tables 4 and 5. The respective pairwise median cluster totals for Bin-SBM and Het-SBM are [9,10] across all three designs. Their corresponding ARI median values (see Figure 7) are [0.89,1], [0.76,1] and [0.66,1] across all three designs. The decrease of ARI scores observed for Bin-SBM across the three designs is linked to the increasing size of the first two clusters which tend be merged by Bin-SBM. In contrast to this, depending on the tuning parameter values, the modular algorithm estimates of 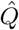 show a wide scope of fitted values ranging from 1 to less than 200 clusters. The default value 1 seems to yield less extreme estimates across all three designs with pairwise median cluster values for FL-A and NS-A [7,7], [5,4] and [4,3], and FL-C and NS-C [8,7], [17,6] and [14,5]. The ARI scores for such cases seems to show a decreasing accuracy across the three designs for FL-A and NS-A with their respective median values of [0.59,0.59], [0.32,0.27] and [0.19,0.15]. The median values for FL-C and NS-C suggest some improvements across the three designs [0.73,0.66], [0.85,0.72] and [0.91,0.83].

**Figure 6:**
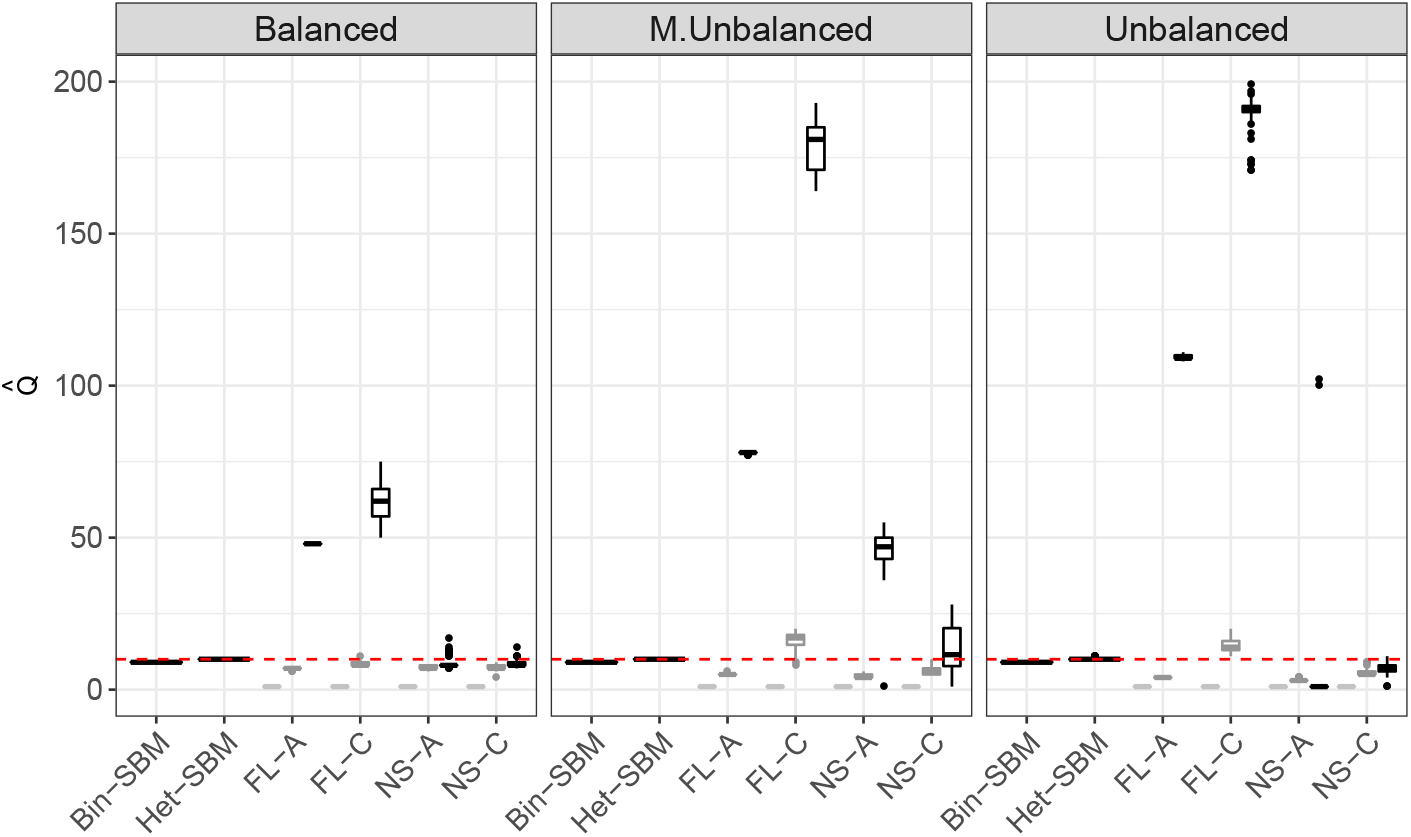
Boxplots showing the estimates of total number of clusters 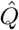 over 100 Monte Carlo samples in terms of ARI scores. FL and NS stand for Fast Louvain and Newman Spectral algorithms while -A and -C indicate the strategy to use the algorithm on the average adjacency matrix and the consensus strategy, respectively. For space considerations, only the results with *γ* = 0.5 (light grey), 1 (mild grey) and 1.5 (black) are shown for the modular algorithms (see **SI** K Figures S17, S19 and S21 for the full results).

**Figure 7:**
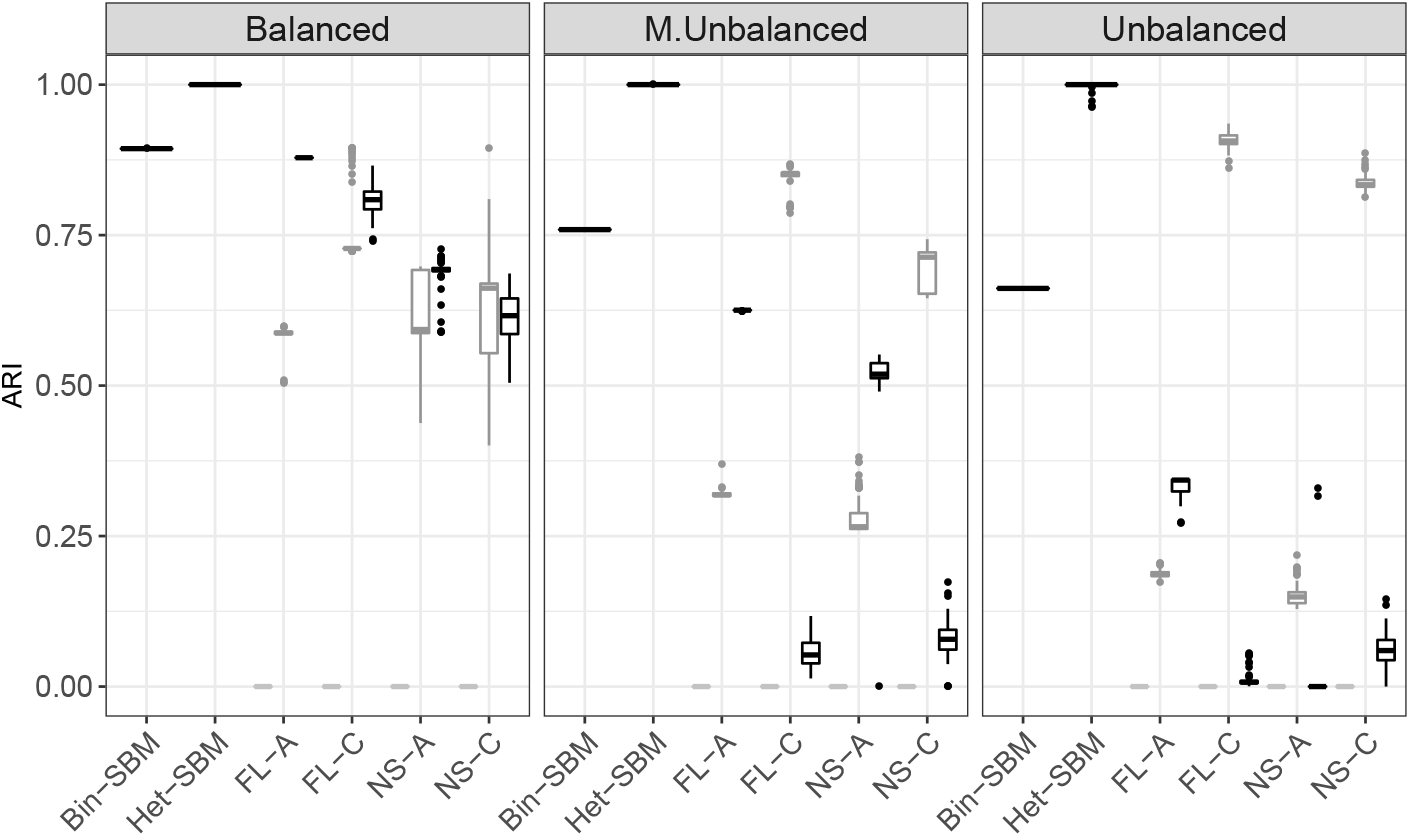
Boxplots showing the recovery of the true cluster assignments over 100 Monte Carlo samples in terms of ARI scores. FL and NS stand for Fast Louvain and Newman Spectral algorithms while -A and -C indicate the strategy to use the algorithm on the average adjacency matrix and the consensus strategy, respectively. For space considerations, only the results with *γ* = 0.5 (light grey), 1 (mild grey) and 1.5 (black) are shown for the modular algorithms (see **SI** K Figures S18, S20 and S22 for the full results).

### 3.5 Simulation III Goals and Setup

#### Goals of Simulation III

The goal of *Simulation III* is to assess the accuracy of the proposed parametric and non-parametric inference procedures in order to give recommendations for real data analyses. A special focus is given to the inferential validity of the parametric (Wald and likelihood ratio) and non-parametric (permutation) tests.

#### Setup of Simulation III

The total number of subjects has been set to 10, 20 and 40 (*K* ∈ {10, 20, 40}), which corresponds to the typical number of subjects in a neuroimaging study. In the interest of computational feasibility and simplicity of presentation, the total number of clusters was set to 3 (*Q* = 3) and the networks with 30, 60 and 120 nodes (*n* ∈ {30, 60, 120}) were utilised. It is worth noting that this number of clusters is sufficient as this simulation is not testing the ability of the multi-subject models to recover the true cluster labels, but rather the overall accuracy of their inference procedures. Similarly to the previous simulations (Simulation I & II), the individual block sizes range according to the Balanced, M. Unbalanced and Unbalanced designs (see **SI L** Table S8 for exact details). Network data was generated according to the following logistic regression model. A logit transformation of the four cluster structures (**π**_1_, …, **π**_4_) shown in Figure 8 (a) was utilised for (1) the intercept and a random sample of values between 20 and 60 with replacement (2) was utilised for an age covariate and its effect was set to 0. It is important to highlight that the age effects are set to be the same for all 6 blocks and, although it is possible to consider heterogeneous effects of varying sizes, such setups would not bring any new insights other than make our analysis more cumbersome. In addition to this, we also differentiate between two cases. In the first case, we preserve the independence of edges within a block (no random effect block) and, in the second case, we violate this assumption by introducing dependences between edges within each block (random effect block). In the first case, the data is generated in the same way as in Simulation II. In the second case, the data is generated according to subject-specific connectivity rates obtained on the logit scale from 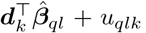, where *u*_*qlk*_ is a block and subject-specific random intercept obtained from the Standard Normal distribution and 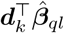 is the fixed effect data. The use of a random effect allows us to introduce dependence between the edges within each subject-specific block. This scenario is relevant as dependences between the edges of fitted blocks are quite plausible in real neuroimaging data and this allows us to assess the likely behaviour of the proposed parametric and non-parametric tests in a more realistic setting.

**Figure 8:**
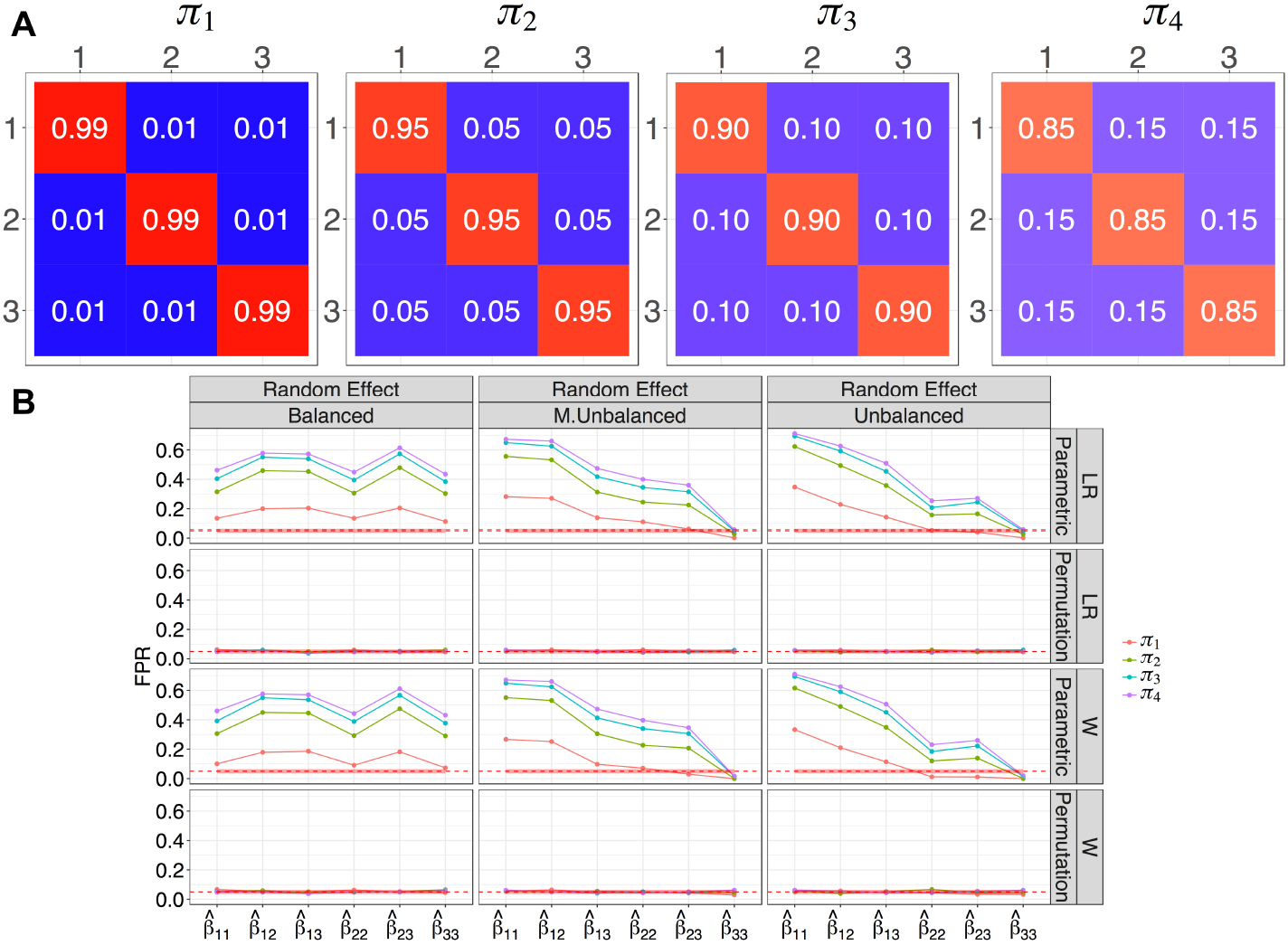
A: Four different connectivity rates (*π*_1_, …, *π*_4_) are used to simulate various levels of intra-block and inter-block connectivity in a homogeneous modular cluster structure. B: Observed FPR for the simulated networks with 30 nodes, 10 subjects and random effect. The *x*-axis represents the block specific fitted regression coefficients. The observed FPRs are colour coded according to 4 different connectivity matrices. The columns represent 3 different block designs and the rows denote the likelihood ratio (LR) and Wald (W) scores based on the parametric and permutation tests. The red shaded strip represents the 95% Monte Carlo binomial proportion confidence intervals. Overall the parametric tests show highly inflated FPRs while the permutation test is accurate.

In total, we simulated 1000 matrices for each condition of the simulations and we evaluated how well each of the parametric (Wald and LR) and non-parametric (permutation) inference procedures controlled false positive rates (FPR). The *P*-values of the Monte Carlo permutation test statistic were obtained by computing 1000 permutations of the age covariate in each of the simulations.

### 3.6 Simulation III Results

The average observed false positive rates were generally well controlled by all inferential procedures under all conditions of simulation when there was no random effect of block or dependence between edges in the same block (see Table 6). However, in a more realistic scenario of a per block random effect, parametric tests of both the Wald and likelihood ratio statistics had severely uncontrolled FPRs; whereas permutation tests of both statistics maintained FPR ~ 0.05 over all conditions; see Table 6 and Figure 8 (b). The failure of the parametric tests to control the FPRs is driven by the assumption of independence of edges within a block which is clearly violated. However, the assumption of exchangeability of the permutation tests is still valid as the presence of dependences between edges within a subject does not impact the exchangeability between subjects. The full set of results for all subjects and cases can be found in **SI** L.

**Table 6:** Average observed FPR at 5% level significance in several scenarios. The average is taken across the 6 blocks, the 4 connectivity matrices and the 3 block size designs.

|  |  | No random effect |  |  |  | Random effect |  |  |  |
| --- | --- | --- | --- | --- | --- | --- | --- | --- | --- |
|  |  | Parametric |  | Permutation |  | Parametric |  | Permutation |  |
|  |  | LR | W | LR | W | LR | W | LR | W |
| 30 nodes | 10 subjects | 0.044 | 0.035 | 0.053 | 0.050 | 0.338 | 0.324 | 0.052 | 0.051 |
|  | 20 subjects | 0.045 | 0.042 | 0.049 | 0.048 | 0.353 | 0.347 | 0.050 | 0.049 |
|  | 40 subjects | 0.050 | 0.046 | 0.051 | 0.050 | 0.367 | 0.363 | 0.052 | 0.052 |
| 60 nodes | 10 subjects | 0.049 | 0.045 | 0.051 | 0.050 | 0.554 | 0.546 | 0.053 | 0.052 |
|  | 20 subjects | 0.049 | 0.048 | 0.050 | 0.050 | 0.557 | 0.555 | 0.049 | 0.049 |
|  | 40 subjects | 0.050 | 0.049 | 0.051 | 0.051 | 0.581 | 0.579 | 0.053 | 0.053 |
| 120 nodes | 10 subjects | 0.051 | 0.049 | 0.053 | 0.053 | 0.735 | 0.733 | 0.052 | 0.051 |
|  | 20 subjects | 0.050 | 0.050 | 0.052 | 0.052 | 0.742 | 0.741 | 0.051 | 0.051 |
|  | 40 subjects | 0.051 | 0.050 | 0.052 | 0.052 | 0.755 | 0.755 | 0.053 | 0.054 |

## 4. Multi-Subject Resting State fMRI Data

Functional magnetic resonance imaging (fMRI) data were available on 13 healthy volunteers (Controls) and 12 patients with schizophrenia (Cases), who have been scanned at rest after being given placebo medication; (see Lynall et al., 2010, for details). Subjects were instructed to quietly lay in the scanner with their eyes closed for 17 mins and 12 s. In each session, a total of 512 scans were acquired with repetition time of 2 s. Each such data set was corrected for motion artefacts and then registered to the MNI standard space. A Gaussian kernel of 6 mm was used to spatially smooth registered images and the time series were high-pass filtered with a cutoff frequency of ≈ 0.008 Hz. Each image was parcellated into 325 anatomically defined regions (ROIs) using the AFNI TT N27 EZ ML atlas (in the Talairach coordinate system) and 28 regions were discarded due to missing data for some individuals. As the majority of these were from the cerebellum, the remaining 29 cerebellar nodes were also removed (see **SI** M Figure S40). Averaged voxel time series in each region were further decomposed into four frequency scales by the discrete wavelet transform (Percival and Walden, 2000). Our subsequent analysis considers correlations between regional fMRI time series in the frequency interval of 0.06 − 0.125 Hz, which was previously shown to be the low frequency range most strongly associated with Case-Control differences in pairwise wavelet correlations in these data (Lynall et al., 2010; Bullmore et al., 2001). These pairwise inter-regional estimates of band-passed correlations between low frequency fMRI oscillations were compiled to form a (268 × 268) functional connectivity matrix (((*r*_*ijk*_))_1≤*i,j*≤*n,k*_) for each subject *k* and then transformed by Fisher’s formula (Fisher, 1915)

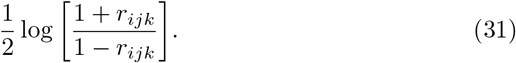

Under the null hypothesis that the population correlation is zero, the Fisher transformation statistic follows a Normal distribution with zero mean and variance 1/(*T* − 3), where *T* is the number of discrete wavelet coefficients within a given frequency band; in this study *T* = 128. On this basis, adjacency matrices can be constructed by testing if the Fisher transformation statistics are significantly greater than zero. Noting that this entails a large number of multiple, non-independent tests (35,778), the false discovery rate (FDR) was controlled at the 5% level (Benjamini and Yekutieli, 2001). If the results were significant, the value 1 was assigned and 0 otherwise. In this manner, each subject *k* is represented by an adjacency matrix ((*x*_*ijk*_))_1≤*i,j*≤*n*_ that captures a binary and undirected graph (or network).

To this set of adjacency matrices, we fit the Het-SBM, described in Section 2.4, including as covariates in the regression model a common intercept for both groups (Cases − Controls), differential effect (Cases − Controls), and mean centred: age, premorbid intelligence, and head movement parameters.

## 5. Multi-Subject Resting State fMRI Data Results and Inference

The Het-SBM estimated a common cluster structure across both groups (Cases & Controls) comprising 21 blocks, whose anatomical labels can be found in **SI** N Figures S42 and S43. Figure 9 (A) shows the average network data reorganised in terms of the 21-block fit found by the Het-SBM. As shown in Figure 9 (B), the network structure does not appear to be modular or core-modular but seems to exhibit a structure with a varying degree of connectivity pattern per cluster with some clusters acting strongly as a core (e.g., Block 1, 2, 4, 12, 17), some being mildly connected to other clusters (e.g., Block 3, 7, 10, 13 & 20) and finally some clusters acting more like modular cluster being almost not connected to any other clusters (e.g., Block 5 & 21). We can also observe that Het-SBM separate well some clusters that may have some high degree of connection between them, but a rather different profile of connection with the other clusters. For example, Blocks 5, 10 exhibits a high and between-block connectivity between their nodes, but they are separated by Het-SBM because their connectivity profile to the other clusters are clearly different.

**Figure 9:**
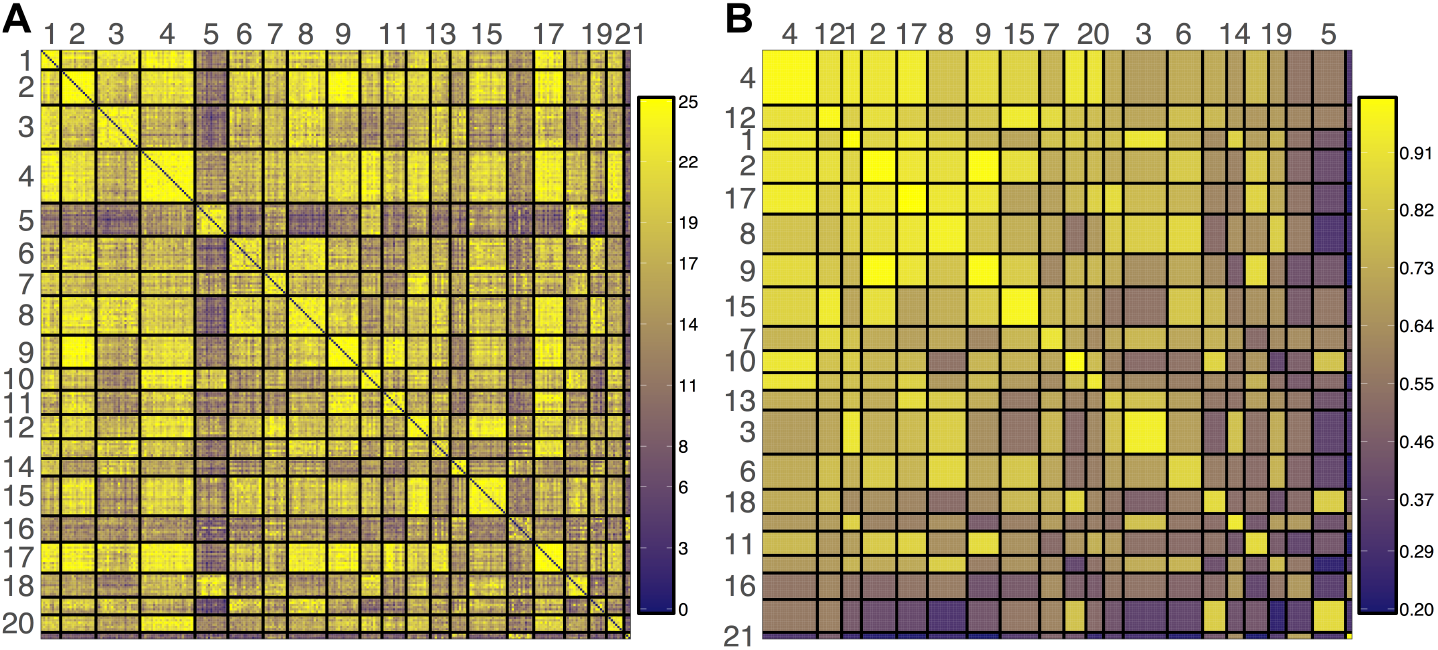
A: Grand average network reorganised in terms of the 21-block obtained from the Het-SBM fit. B: Fitted connectivity matrix from the Het-SBM fit, ordered by decreasing average block probabilities. The network structure seems to present a non-modular structure.

In Figure 10, we show the family-wise error rate corrected *P*-values for the test of difference between the average block connectivity levels of Patient and Control groups based on the LR test (shown in (A)) and permutation test (shown in (B)). The non-significant block *P*-values, thresholded at a 5% level significance, are given in white while the significant block *P*-values are plotted on a −log_10_(·) scale so that, for example, the value of 305.3 represents the *P*-value of 10^−305.3^.

**Figure 10:**
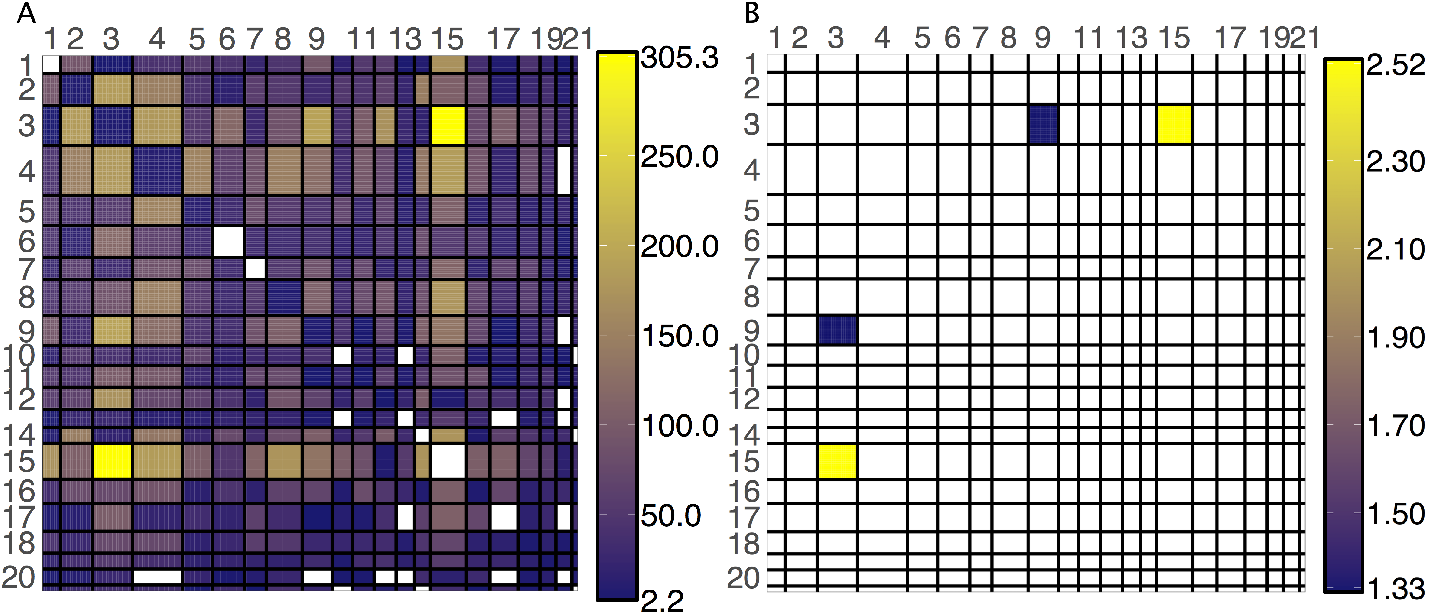
Family-wise error rate corrected significant *P*-values for the null hypothesis of no average difference in connectivity between the Patient and Control groups with A showing the results of the LR test and B showing the results of the permutation test based on the LR statistic. The *P*-values are stated in terms of −log_10_(*P*-values) (e.g., the legend score of 305.3 corresponds to the *P*-value of 10^−305.3^) and the non-significant *P*-values at 5% significance are given in white. Overall the LR test gives a very large number of significant results which are not detected with the permutation test. This corresponds to the example considered in Simulation III, where the additional correlations between the edges in a block (i.e. random effects) are shown to cause strongly inflated false positives in the asymptotic tests making them invalid. The permutation test seems to be robust to the block level dependences between the edges and since the exchangeability assumption is not violated, we see much less activations. The significant results relate to the connections between the Block 3 & 9 and are especially strong for the connections between Block 3 & 15.

There is a striking difference between the two tests in terms of the number of significantly declared blocks. In particular, the LR test seems to declare a total of 215 significant blocks which is almost 107 times more than the number of significant blocks declared by the permutation test. This particular scenario is largely consistent with the results of Simulation III and is suggestive of an invalid LR test. Indeed, in Simulation III, we have already showed that when there is some sort of dependence between the edges of a block (i.e. per block random effect) the asymptotic test tend to have profoundly inflated false positive rates. To investigate this on the real multi-subject data, we conducted a parametric bootstrap test based on LR statistics to find an evidence of a random effect in each of the blocks (the full details on the fitting and hypothesis testing procedures are given in SI M.1). With the exception of Block (1,1) and Block (17,17) all the remaining fitted blocks in Figure S41 show significant random effects. Given the nature of functional connectivity data and strongly coupled correlation patterns, it is not surprising that the dependences between the nodes continue to persist in the data, even after the clustering model is fitted. Interestingly, such dependences in Simulation III showed almost no particular impact on the clustering results but proved to be very disastrous for the control of false positive rates in the asymptotic tests. In contrast, the overall performance of the permutation test is unaffected by this condition and the false positive rates are reasonably well controlled. This specific observation is echoed in Figure 10 (B) where we only see two significant blocks, mainly the connectivity between Block 3 & 9 and Block 3 & 15.

In order to see the direction of differential effect (Cases vs. Controls) and the anatomical locations of nodes in Block 3, 9 and 15, we look at the observed differences in the connections between the two groups relative to their total number of subjects (as shown in Figure 11). Overall, the connectivity rates between the blocks tend to be much weaker in the Patients than Controls and this is consistent with the general literature which classifies schizophrenia as ‘dysconnectivity disorder’ (Friston and Frith, 1995). This profile of attenuated functional connectivity between areas of temporal, frontal and cingulate cortex is compatible with other graph theoretical and functional connectivity studies of schizophrenia that have emphasised fronto-temporal profiles of dysconnectivity (Fornito et al., 2012). The observation that functional dysconnectivity in schizophrenia is restricted to a subset of clusters is more novel but consistent with previous studies demonstrating that abnormal developmental trajectories of cortical shrinkage in schizophrenia were restricted to a specific module of the structural network community structure comprising areas of frontal, temporal and cingulate cortex (Alexander-Bloch et al., 2013, 2014). It seems that the cluster structure of brain networks may constrain the expression of dysconnectivity phenotypes in schizophrenia. This in turn may be related to the emerging idea that the modular organisation of brain networks reflects co-expression of genes and that genes associated with a risk for schizophrenia have a non-uniform expression across the cortex with high levels of schizophrenia risk gene expression associated with developmental changes in structure of frontal cortex (Whitaker et al., 2016).

**Figure 11:**
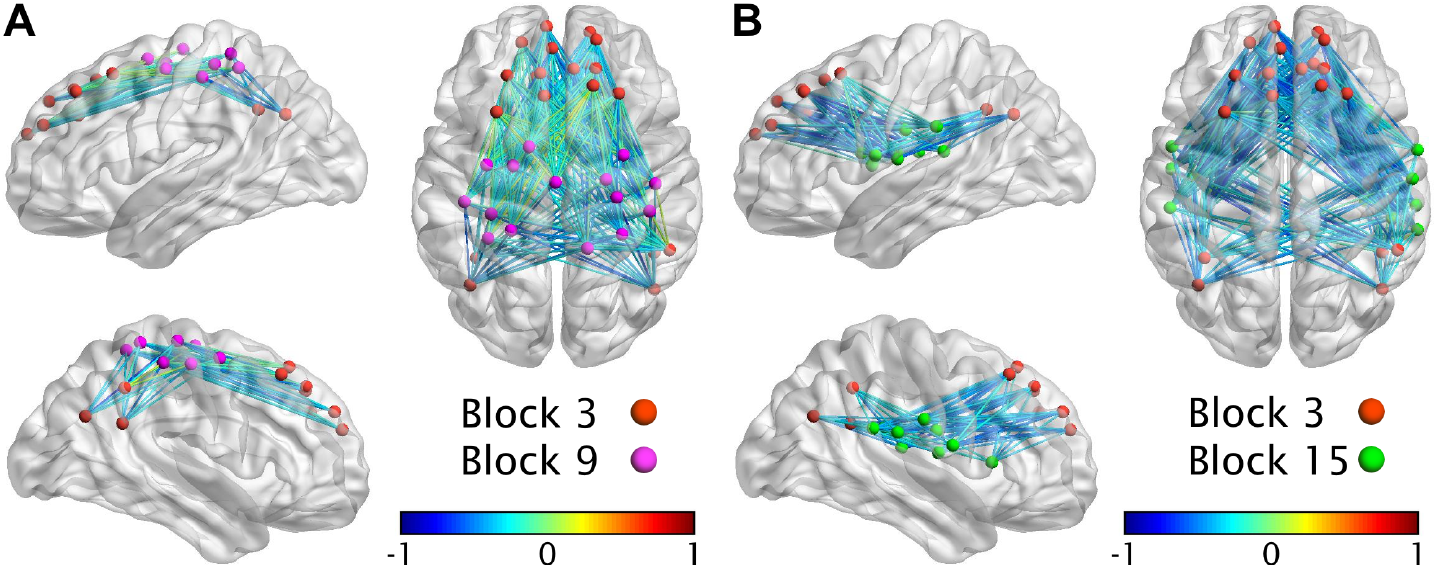
Between group average differences in the observed connectivity between Block 3 & Block 9 (A) and Block 9 & Block 15 (B) relative to the total number of subjects in each group. Overall, the connectivity strengths between blocks tend to be much weaker in the Patients than Controls.

## 6. Discussion

This work proposed a multi-subject framework based on three extension of the classical SBM, referred to as Bin-SBM, Hom-SBM and Het-SBM. The last two models are non-trivial and use subject specific covariates to explain variations in cluster structure between subjects. Focusing on Het-SBM, this work benchmarked it against classical modular algorithms, explored the validity of its inference procedures based on parametric and non-parametric tests and how these could be used in a real data analysis to test for a group effect between healthy controls and patients diagnosed with schizophrenia.

### 6.1 Benchmarking the Models

To investigate the modelling strengths and weaknesses of Het-SBM and Bin-SBM, we conducted three separate simulations (Simulation I-III).

#### Simulation I

In Simulation I, we compared our models to popular clustering methods such as the Newman Spectral (NS) and Fast Louvain (FL) algorithms, based on (1) average (-A) and (2) consensus clustering (-C), with different values for the resolution parameter *γ*. Overall, MS-SBMs outperformed the modular algorithms and were accurate in all scenarios. The modular algorithms showed accuracy in the Hom-Modular structure, and reasonably balanced cluster sizes with an almost negligible effect of *γ*. In the Hom-Modular structure with unbalanced cluster sizes, consensus clustering was found to perform better than average clustering. However, average clustering could potentially be improved with larger values of *γ*. For Het-Modular and Core-Modular structures, the modular solutions were found to be less accurate, with a higher dependence on *γ*. This suggests that, in practice, special care must be taken when choosing the value of *γ*, especially when the cluster structure deviates from a pure Hom-Modular structure. Another problematic aspect is that the resolution parameter can lead to a very strong over estimation of the total number of clusters and introduce partitions which are not parsimonious. In Het-SBM, this problem is solved with the ICL criterion which penalises the log likelihood for the complexity of the model and preserves parsimony. Unlike modularity scores which cannot be compared over different values of *γ* (controlling the total number of clusters), ICL scores are comparable across different values of *Q*. Furthermore, ICL scores can be used to evaluate the partitions obtained through any clustering method, which allows to assess their goodness of fit.

#### BN-SBM in Simulation I

In addition to the classical modular algorithms, we also benchmarked Block to Node SBM (or BN-SBM) (Newman and Leicht, 2007), which can be regarded as a special case (or special re-parametrisation) of the classical SBM of Snijders and Nowicki (1997) and Nowicki and Snijders (2001) (see **SI** I for details). There are, however, two main reasons as to why this model could not be benchmarked in the full simulation setting but only on a subset of cases. First, this model is intended for the analysis of a single, binary network, and our simulations are multi-subject. Second, the model and its C based implementation use estimated likelihood scores to compare different fits, which are only valid for partitions with the same *Q*. Unlike MS-SBMs, this model can only be benchmarked when the ground truth cluster number is supplied (i.e. *Q* was set to 10). As shown in Figure S8, BN-SBM performs very well in the case of Hom-Modular structures, but it is less accurate in Hom-Modular and Core-Modular cases. One possible explanation is that the combination of block-to-node parametrisation and the parametric constraints of Categorical density (i.e. that the parameters must add up to 1 across the nodal categories) pose limitations on the types of cluster structures that the model can estimate. It appears that the modelling of nodal connectivity profiles comes at a heavy cost to the model’s richness and ability to estimate different cluster structures, a trade-off that we have found to be unsatisfactory.

#### Simulation II

In Simulation II, we showcased an example in which the between-subject variation induced by a covariate strongly influenced the overall cluster structure, where two clusters can erroneously be merged by Bin-SBM but correctly estimated by Het-SBM. This demonstrates that it can be very important to account for between-subject variations in cluster structure to estimate the cluster labels. Therefore, it is not advisable to perform clustering separately from the regression analysis.

#### Simulation III

In Simulation III, we tested the accuracy of Het-SBM inference procedures based on parametric and non-parametric tests. Overall, non-parametric (permutation) tests were found to be more accurate than the Wald and LR tests. When the assumptions of independences between the edges in a block are satisfied, parametric and non-parametric test were both found to be valid. However, in instances of dependence between edges in a block, the Wald and LR were found to be invalid due to the more liberal control of FPRs, while the permutation testing remained accurate method. These results lend strong support for permutation testing as the most reasonable inference method for situations where independence between block edges cannot be assumed. Nevertheless, it is important to note that permutation testing relies on assumption that data is exchangeable under the null hypothesis. This assumption may not hold if the data exhibits some form of heteroskedasticity. For example, if we assume that data is generated through a mixed effects model with a different random intercept per group, data will not be exchangeable between groups, and the permutation test may not be valid.

#### Inference with Het-SBM in Real Data

In this paper, Het-SBM was illustrated in an application to a resting state fMRI study with healthy subjects and subjects diagnosed with schizophrenia, yielding a fit with 21 clusters. Interestingly, this data contained the same type of edge dependencies in the blocks as in Simulation III, which made LR test liberal. A much more reliable inference was achieved by the permutation test which found significant differences between the two groups in Block(3,9) and Block(3,15), containing nodes in temporal, frontal and cingulate cortex.

### 6.2 Generalised MS-SBMs

#### Non-binary edges

Using the general approach outlined in this paper, it is also relatively simple to derive other variants of Het-SBM or Hom-SBM for non-binary edges. As any density in exponential family of distributions can be utilised to describe the edges in a dataset, the fitting strategies described in this work can be utilised in the same way (as mentioned above in this paper), to derive a variant of Het-SBM that suits the edges of the dataset. For example, if researchers are interested in clustering observed correlations between ROI time series (without a thresholding step), one could utilise Fisher Z to transform the correlations and subsequently assume that edges are normally distribution. This greatly simplifies estimation procedure as a Normal linear model utilises closed form solutions for maximum likelihood estimates. Likewise, one can imagine the uses for this type of modelling in networks obtained from diffusion tensor imaging. For example, if the edges represent white matter fibre counts between the ROIs, a Poisson distribution might be a reasonable fit for such networks. Subsequently, the use of a Poisson regression model which would invoke the Fisher scoring step as in this work. In general, depending on the type of distribution used in the modelling of the edge data, researchers could choose an appropriate generalised linear model while still applying the methodology discussed in this work in order to develop a Het-SBM model tailored to their dataset.

#### Edge-wise covariates

While, in this work, we mainly focused on multi-subject models where the distribution of edges is dependent on subject-specific covariates like age or gender, it is also possible to adapt Het-SBM and Hom-SBM for a distribution of edges that is dependent on edge-level or node-level covariates. For example, (1) between regional Eculidean distances, (2) total numbers of tractography streamlines, and (3) correlations based on cortical thickness, can be seen as edge-level covariates. The Euclidean distance can be a strongly informative factor in explaining the cluster structure especially as the long-distance and short-distance connections are suggestive of integration and segregation in the brain. Thus, assuming heterogeneous effects (i.e. Het-SBM), we can write this model as *X*_*ijk*_|*Z*_*iq*_ = 1, *Z*_*jl*_ = 1 ~ Bernoulli(*π*_*ijk*_), where 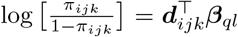 such that ***d***_*ijk*_ is 1 × *P* a vector of edge features associated with the subject *k* and edge *x*_*ij*_. In this particular example, *P* = 4, as the regression model consist of the intercept, Eculidean distance, total number of tractography streamlines and cortical thickness correlations. It is worth noting that in this particular context *π*_*ijk*_ is not treated as parameter. Instead, the only parameters in the model are associated with the regression coefficients ***β***_*ql*_. By setting ***d***_*ijk*_ to be equal to ***d***_*k*_, this model becomes exactly the Het-SBM discussed in this work. In this version of the model, edge-level covariates could have a much stronger influence on the estimation of cluster assignments than subject-level covariates, so it would be interesting to see how much of this influence is tied to the homogeneous and heterogenous versions of the model. In parallel to this, it would be also interesting to detect a collection of edge-covariates which can explain clustering of network data that give rise to known resting state networks. By doing this type of analysis the researchers could obtain a host of potential network based biomarkers which could potentially be more closely tied to specific brain disorders.

#### Nodal covariates

It is also possible to consider the use of node-based covariates for modelling distribution of edges. One way to account for them is to simply transform those node-based covariates into edge-based covariates using a set of meaningful symmetric transformation function *f*_*p*_(*a, b*) such that the element of the edge-based covariate *p* corresponding to the nodes *V*_*i*_ and *V*_*j*_ is given by *f*_*p*_(*d*_*ikp*_, *d*_*jkp*_). The choice these functions would depend on the nature of the node-based covariates and could be, for example, the mean function 1*/*2(*a* + *b*) which would assume an addictive effect of the node-based covariate values on the edge connectivities.

However, a potential limitation to these edge-covariate models is the possible violation of the exchangeability assumption that renders the permutation testing inappropriate. In such a situation, alternative resampling strategies will be needed, and the accuracy of selected inference procedures must be investigated.

### 6.3 Limitations and Future Work

It is important to note that the current setup of our proposed approach does not allow for per-subject varying cluster labels. This is a limitation of our model that might be important to overcome as some studies have identified that cluster assignments of nodes substantially change as a function of task (Hearne et al., 2017) and diagnostic status (Alexander-Bloch et al., 2012). Nevertheless, we are working on expanding the scope of our model to account for such variation in our next research project. The current model makes inferences on covariate effects which are dependent on cluster label fit. The accuracy of the fit is paramount in the interpretation of the results, and it is closely related to the functionally meaningful block structures. If blocks cannot be associated with a particular brain function, it may results in statistically significant blocks without any meaningful interpretation. This may be circumvented by conducting an edge-specific inference. However, this strategy goes against the principle of parsimony, currently enforced by making block-specific inference. Moreover, edge-specific inferences would drastically increase the number of tests, worsening the issue of multiple comparisons. Therefore, while it is indeed possible to conduct edge-wise inferences, block-specific inferences might be preferable, particularly if the blocks can be associated to meaningful functions. In addition, it is worth noting that the issue of selecting optimal clustering is not uniquely specific to our model, but to all clustering methods. In general, given the same dataset, all clustering methods experience similar limitations as they explore the same space of candidate models (i.e. potential cluster labels) and this space for a particular *Q* has a finite size of *Q*^*n*^. Given that this is the extremely high number of fits to consider, it is very difficult to ensure that the final model is indeed a global solution, and not merely a reasonable local maximum. In our analysis of real data, Het-SBM yielded a fit with 21 clusters and significant differences between the two groups in two blocks. Ideally, a repeated sample would be needed to confirm these findings, especially as the sample size of the dataset is rather small, which may have prevented to discover other significant associations. Thus, it would be interesting to repeat this analysis on a larger cohort and to see how robust and reproducible these results are.

Finally, we would like to invite the general graph network research community to consider validation of available clustering methodology through simulations. This would allow researchers to quickly check whether their choices of clustering methods conform to their expectations and to avoid to read too much into the results which could have been generated by the weakness of the clustering method and not the data. Given the flexibility of the SBMs and multi-subject SBM considered in this work, we remain hopeful that neuroimaging researchers will consider these seriously.

## Acknowledgements

We would like to thank Eugene Demidenko, David Firth, Iréne Gannaz, Georg Heinze and Stéphane Robin for valuable discussions at various stages of this project.

DMP is supported by the MRC Industrial CASE award with the GlaxoSmithKlines Clinical Unit Cambridge (UK) PhD studentship. BRLG is supported by the EU within the PEOPLE Programme (FP7): Initial Training Networks (FP7-PEOPLE-ITN-2008), Grant Agreement No. 238593 NEUROPHYS-ICSEKT. EKT is funded by Medical Research Council. PEV is supported by the Medical Research Council (grant number MR/K020706/1). YBTT is supported by NUS Tier 1, Singapore MOE Tier 2 (MOE2014-T2-2-016), NUS Strategic Research (DPRT/944/09/14), NUS SOM Aspiration Fund (R185000271720), Singapore NMRC (CBRG14nov007, NMRC/CG/013/2013) and NUS YIA. ETB is employed half-time by the University of Cambridge and half-time by GlaxoSmithKline; and holds stock in GSK. The Behavioural and Clinical Neuroscience Institute is supported by the Medical Research Council (UK) and Wellcome Trust. TEN is supported by NIH U54MH091657-03 and the Wellcome Trust. The funders had no role in study design, data collection and analysis, decision to publish, or preparation of the manuscript.

## Footnotes

1 Rare events refer to an unusual distribution of 0’s and 1’s in a sample where enormous disparity between their individual counts is expected.

2 Complete separation is a special case in the logistic regression, where it is possible to establish a perfect correspondence between all possible values of a covariate and {0, 1} outcomes in data. For example, complete separation has occurred when fitting a logistic regression curve to the block (*q, l*), if all smokers have an edge in this block while all the subjects who are not smokers do not have an edge in this block.

3 The clusters in a modular organisation are characterised by maximised within-cluster connectivity and minimised between-cluster connectivity. The probabilities of connections within each cluster are assumed to be equal and the probabilities of connections between clusters are also assumed to be equal.

4 17 different values of *Q* times 30 restarts

