## Supplementary Material for "Multi-Subject Stochastic Blockmodels for Adaptive Analysis of Individual Differences in Human Brain Network Cluster Structure"

Contents

**Appendix A Derivation of the Variational Bound** **3**

**Appendix B Derivation of the Fixed Point Estimates** **5**

**Appendix C Derivation of Integrated Classification Likelihood (ICL) Criterion** **8**

**Appendix D Optimisation Algorithm for Bin-SBM** **10**

D.1 Numerical Stability of Cluster Assignment Probabilities . . . . 10

D.2 Variational EM algorithm for Binomial SBM (Bin-SBM) . . . 12

**Appendix E Estimation in Homogeneous Stochastic Blockmodel (Hom-SBM)** **14**

**Appendix F Optimisation algorithm for Heterogeneous SBM (Het-SBM)** **20**

**Appendix G Evaluation methods** **23**

**Appendix H Results for Simulation I** **24**

**Appendix I Comparisons to Block to Node SBM for Simulation I** **37**

**Appendix J Initialisation Strategy Comparison** **44**

**Appendix K Additional Results for Simulation II** **52**

**Appendix L Additional Results for Simulations III** **56**

**Appendix M Real Data Analysis** **75**

M.1 Het-SBM Inference Results . . . . . . . . . . . . . . . . . . . . . . 77

**Appendix N Additional Results for Schizophrenia Study** **77**

### Web-based Supplementary Information for Stochastic Blockmodelling and Inference in Multi-Subject Networks with Mixture Models

Dragana M. Pavlović[a,b], Bryan R. L. Guillaume[c], Emma K. Towlson[d], Petra E. Vértes[e], Soroosh Afyouni[h], B. T. Thomas Yeo[b], Edward T. Bullmore[e,f,g], Thomas E. Nichols[a,h]

[a] *Department of Statistics, University of Warwick, Coventry, United Kingdom*
[b] *Department of Electrical and Computer Engineering, Clinical Imaging Research Centre, Singapore Institute of Neurotechnology and Memory Networks Programme, National University of Singapore*
[c] *Department of Biomedical Engineering, National University of Singapore, Singapore*
[d] *Theory of Condensed Matter Group, Department of Physics, Cavendish Laboratory, University of Cambridge, Cambridge CB3 0HE, United Kingdom*
[e] *Behavioural and Clinical Neuroscience Institute, Department of Psychiatry, University of Cambridge, Cambridge CB2 0SP, United Kingdom*
[f] *Cambridgeshire and Peterborough National Health Service Foundation Trust, Cambridge CB21 5EF, United Kingdom*
[g] *GlaxoSmithKline, Clinical Unit Cambridge, Addenbrooke's Hospital, Cambridge CB2 0AA, United Kingdom*
[h] *Warwick Manufacturing Group, University of Warwick, Coventry, United Kingdom*

#### A. Derivation of the Variational Bound

In order to describe the variational approach, we first explicitly derive a lower bound for the incomplete data, represented by the marginal density $f(\boldsymbol{x};\boldsymbol{\pi},\boldsymbol{\alpha})$. Denoting a sample space of $\boldsymbol{Z}$ as $\mathcal{S}$, we can derive the following inequality

$$\begin{aligned}
\log f(\boldsymbol{x};\boldsymbol{\pi},\boldsymbol{\alpha}) &= \log\left[\sum_{\boldsymbol{z}\in\mathcal{S}} f(\boldsymbol{x},\boldsymbol{z};\boldsymbol{\pi},\boldsymbol{\alpha})\right] \\
&= \log\left[\sum_{\boldsymbol{z}\in\mathcal{S}} f(\boldsymbol{x},\boldsymbol{z};\boldsymbol{\pi},\boldsymbol{\alpha})\frac{f^*(\boldsymbol{z};\boldsymbol{\tau})}{f^*(\boldsymbol{z};\boldsymbol{\tau})}\right] \\
&= \log\left[\mathbb{E}_{f^*}\left(\frac{f(\boldsymbol{x},\boldsymbol{Z};\boldsymbol{\pi},\boldsymbol{\alpha})}{f^*(\boldsymbol{Z};\boldsymbol{\tau})}\right)\right] \\
&\geq \mathbb{E}_{f^*}\left(\log\left[\frac{f(\boldsymbol{x},\boldsymbol{Z};\boldsymbol{\pi},\boldsymbol{\alpha})}{f^*(\boldsymbol{Z};\boldsymbol{\tau})}\right]\right) \quad \text{(by Jensen's inequality)} \\
&= \mathbb{E}_{f^*}\Big(\log[f(\boldsymbol{x},\boldsymbol{Z};\boldsymbol{\pi},\boldsymbol{\alpha})]\Big) - \mathbb{E}_{f^*}\Big(\log[f^*(\boldsymbol{Z};\boldsymbol{\tau})]\Big). \qquad \text{(SI1)}
\end{aligned}$$

In particular, the lower bound in Eq. (SI1) is closely related to the Kullback-Leibler (KL) divergence of $f^*(\boldsymbol{z};\boldsymbol{\tau})$ to $f(\boldsymbol{z}|\boldsymbol{x};\boldsymbol{\pi},\boldsymbol{\alpha})$, as

$$\begin{aligned}
\mathrm{KL}\left[f^*(\boldsymbol{z};\boldsymbol{\tau})\middle|\middle| f(\boldsymbol{z}|\boldsymbol{x};\boldsymbol{\pi},\boldsymbol{\alpha})\right] = &\log f(\boldsymbol{x};\boldsymbol{\pi},\boldsymbol{\alpha}) \\
&- \left[\mathbb{E}_{f^*}\Big(\log[f(\boldsymbol{x},\boldsymbol{Z};\boldsymbol{\pi},\boldsymbol{\alpha})]\Big)\right. \\
&\left. - \mathbb{E}_{f^*}\Big(\log[f^*(\boldsymbol{Z};\boldsymbol{\tau})]\Big)\right]. \qquad \text{(SI2)}
\end{aligned}$$

Rearranging the last equation, we write the lower bound Eq. (SI1) instead as

$$\log f(\boldsymbol{x};\boldsymbol{\pi},\boldsymbol{\alpha}) - \mathrm{KL}\left[f^*(\boldsymbol{z};\boldsymbol{\tau})\middle|\middle| f(\boldsymbol{z}|\boldsymbol{x};\boldsymbol{\pi},\boldsymbol{\alpha})\right]. \qquad \text{(SI3)}$$

This shows that the lower bound in Eq. (SI1) is precisely attained when the KL divergence is zero, that is, when $f^*(\boldsymbol{z};\boldsymbol{\tau})$ coincides with $f(\boldsymbol{z}|\boldsymbol{x};\boldsymbol{\pi},\boldsymbol{\alpha})$. As we cannot explicitly calculate $\log f(\boldsymbol{x};\boldsymbol{\pi},\boldsymbol{\alpha})$ due to the intractability of $f(\boldsymbol{z}|\boldsymbol{x};\boldsymbol{\pi},\boldsymbol{\alpha})$, we find a family of densities with variational parameter $\boldsymbol{\tau}$ $f^*(\boldsymbol{z};\boldsymbol{\tau})$ to approximate $\log f(\boldsymbol{x};\boldsymbol{\pi},\boldsymbol{\alpha})$ by its lower bound,

$$\begin{aligned}\log f(\boldsymbol{x};\boldsymbol{\pi},\boldsymbol{\alpha}) - \mathrm{KL}\left[f^*(\boldsymbol{z};\boldsymbol{\tau})\middle|\middle|f(\boldsymbol{z}|\boldsymbol{x};\boldsymbol{\pi},\boldsymbol{\alpha})\right] \geq\\ \mathbb{E}_{f^*}\Big(\log[f(\boldsymbol{x},\boldsymbol{Z};\boldsymbol{\pi},\boldsymbol{\alpha})]\Big)\\ - \mathbb{E}_{f^*}\Big(\log[f^*(\boldsymbol{Z};\boldsymbol{\tau})]\Big).\end{aligned} \tag{SI4}$$

Although the lower bound depends on expectations, it is still easier to work with it directly than with $\log f(\boldsymbol{x};\boldsymbol{\pi},\boldsymbol{\alpha})$. Indeed, since the random variables in $\boldsymbol{Z}$ are independent, the expectations of their products, as we will show later, can be factorised. This suggests that we can maximise $\log f(\boldsymbol{x};\boldsymbol{\pi},\boldsymbol{\alpha})$ (or more precisely the likelihood) without its calculation and, consequently, get the estimates of $\boldsymbol{\tau}$ and other parameters of interest. Note, that the accuracy of the variational approximation is of course dependant of our choice of $f^*(\boldsymbol{z};\boldsymbol{\tau})$ and the true log likelihood. Since the random variables $\boldsymbol{Z}$ are independent, a class of densities $f^*(\boldsymbol{z};\boldsymbol{\tau})$ is generally referred to as mean-field or fully-factored densities. To indicate the role of $\boldsymbol{\tau}$, the lower bound is formally denoted as

$$\mathcal{J}(f^*(\boldsymbol{z};\boldsymbol{\tau});\boldsymbol{\pi},\boldsymbol{\alpha}) = \mathbb{E}_{f^*}\left(\log[f(\boldsymbol{x},\boldsymbol{Z};\boldsymbol{\pi},\boldsymbol{\alpha})]\right) - \mathbb{E}_{f^*}\left(\log[f^*(\boldsymbol{Z};\boldsymbol{\tau})]\right). \quad \text{(SI5)}$$

In particular, the natural option for $f^*(\boldsymbol{z};\boldsymbol{\tau})$ is a categorical distribution with block-specific probabilities, independently for each node:

$$f^*(\boldsymbol{z};\boldsymbol{\tau}) = \prod_{i=1}^{n}\prod_{q=1}^{Q}\tau_{iq}^{z_{iq}}, \quad \text{(SI6)}$$

where $\sum_{q=1}^{Q}\tau_{iq} = 1$. Note, that this gives the probability of node $V_i$ belonging to block $q$ as $\mathbb{E}_{f^*}(Z_{iq}) = \tau_{iq}$ and that $\mathbb{E}_{f^*}(Z_{iq}Z_{jl}) = \tau_{iq}\tau_{jl}$.

#### B. Derivation of the Fixed Point Estimates

In this section, we derive the fixed point equations given by Eq. (8), (9), (10) in the main manuscript. The lower bound can be written as:

$$\mathcal{J}(f^*(\boldsymbol{z};\boldsymbol{\tau});\boldsymbol{\alpha},\boldsymbol{\pi}) = \frac{1}{2}\sum_{i=1}^{n}\sum_{j\neq i}^{n}\sum_{q,l}^{Q}\tau_{iq}\tau_{jl}\log f(x_{ij}|z_{iq},z_{jl};\pi_{ql}) + \sum_{i=1}^{n}\sum_{q=1}^{Q}\tau_{iq}\log\alpha_q - \sum_{i=1}^{n}\sum_{q=1}^{Q}\tau_{iq}\log\tau_{iq}, \quad \text{(SI7)}$$

where $f(x_{ij}|z_{iq},z_{jl};\pi_{ql}) = \binom{K}{x_{ij}}\pi_{ql}^{x_{ij}}(1-\pi_{ql})^{K-x_{ij}}$. The fixed point estimates are found by maximising the variational bound with respect to each parameter. Considering $\boldsymbol{\alpha}$, we first take derivatives with respect to $\alpha_{q'}$ and subject it to the constraint $\sum_{q=1}^{Q} \alpha_q = 1$ with the help of Lagrange multiplier,

$$\frac{\partial}{\partial \alpha_q} \mathcal{J}(f^*(\boldsymbol{z}; \boldsymbol{\tau}); \boldsymbol{\alpha}, \boldsymbol{\pi}) = \frac{\partial}{\partial \alpha_q} \left[ \sum_{i=1}^{n} \sum_{q'=1}^{Q} \tau_{iq} \log \alpha_{q'} - \lambda \left( \sum_{q'=1}^{Q} \alpha_{q'} - 1 \right) \right]$$

$$= \sum_{i=1}^{n} \tau_{iq} - \lambda \alpha_q, \tag{SI8}$$

$$\frac{\partial}{\partial \lambda} \mathcal{J}(f^*(\boldsymbol{z}; \boldsymbol{\tau}); \boldsymbol{\alpha}, \boldsymbol{\pi}) = -\sum_{q=1}^{Q} \alpha_q + 1. \tag{SI9}$$

Solving $\sum_{i=1}^{n} \tau_{iq} - \lambda \alpha_q = 0$ for $\alpha_q$ gives $\alpha_q = \frac{1}{\lambda} \sum_{i=1}^{n} \tau_{iq}$. Setting $-\sum_{q=1}^{Q} \alpha_q + 1 = 0$ gives $\sum_{q=1}^{Q} \alpha_q = 1$. Next,

$$\alpha_q = \frac{1}{\lambda} \sum_{i=1}^{n} \tau_{iq}$$

$$\sum_{q=1}^{Q} \alpha_q = \frac{1}{\lambda} \sum_{i=1}^{n} \sum_{q=1}^{Q} \tau_{iq}$$

$$1 = \frac{1}{\lambda} n$$

$$\lambda = n.$$

This gives that $\hat{\alpha}_q = \frac{1}{n} \sum_{i=1}^{n} \hat{\tau}_{iq}$.

Next, we find fixed point estimate for $\pi_{ql}$,

$$\frac{\partial}{\partial \pi_{ql}} \mathcal{J}(f^*(\boldsymbol{z};\boldsymbol{\tau});\boldsymbol{\alpha},\boldsymbol{\pi}) = \frac{\partial}{\partial \pi_{ql}} \left[ \frac{1}{2} \sum_{i=1}^{n} \sum_{j\neq i}^{n} \sum_{q',l'}^{Q} \tau_{iq'}\tau_{jl'} \log f(x_{ij}|z_{i'q'}, z_{j'l'}; \pi_{q'l'}) \right]$$

$$= \frac{\partial}{\partial \pi_{ql}} \left[ \frac{1}{2} \sum_{i=1}^{n} \sum_{j\neq i}^{n} \sum_{q',l'}^{Q} \tau_{iq'}\tau_{jl'} \log \left( \binom{K}{x_{ij}} \pi_{q'l'}^{x_{ij}} (1-\pi_{q'l'})^{K-x_{ij}} \right) \right]$$

$$= \sum_{i=1}^{n} \sum_{j\neq i}^{n} \tau_{iq}\tau_{jl}x_{ij} - \pi_{ql}K \sum_{i=1}^{n} \sum_{j\neq i}^{n} \tau_{iq}\tau_{jl}, \quad \text{(SI10)}$$

and setting to zero and solving for $\pi_{ql}$ gives $\hat{\pi}_{ql} = \frac{\sum_{i=1}^{n}\sum_{j\neq i}^{n} \hat{\tau}_{iq}\hat{\tau}_{jl}x_{ij}}{K\sum_{i=1}^{n}\sum_{j\neq i}^{n} \hat{\tau}_{iq}\hat{\tau}_{jl}}$.

Finally, we find fixed point estimate for $\tau_{iq}$,

$$\frac{\partial}{\partial \tau_{iq}} \mathcal{J}(f^*(\boldsymbol{z};\boldsymbol{\tau});\boldsymbol{\alpha},\boldsymbol{\pi}) = \frac{\partial}{\partial \tau_{iq}} \left[ \frac{1}{2} \sum_{i'=1}^{n} \sum_{j'\neq i'}^{n} \sum_{q',l'}^{Q} \tau_{i'q'}\tau_{j'l'} \log f(x_{i'j'}|z_{i'q'}, z_{j'l'}; \pi_{q'l'}) \right.$$

$$\left. + \sum_{i'=1}^{n} \sum_{q'=1}^{Q} \tau_{i'q'} \log \alpha_{q'} - \sum_{i'=1}^{n} \sum_{q'=1}^{Q} \tau_{i'q'} \log \tau_{i'q'} \right].$$

The first term has two contributions when $i' = i$ and $q' = q$, and $j' = i$ and $l' = q$. This gives

$$\frac{1}{2} \sum_{j'\neq i'}^{n} \sum_{l'=1}^{Q} \tau_{j'l'} \log f(x_{i'j'}|z_{i'q'}, z_{j'l'}; \pi_{q'l'}) + \frac{1}{2} \sum_{i'\neq i}^{n} \sum_{q'=1}^{Q} \tau_{i'q'} \log f(x_{i'i}|z_{i'q'}, z_{iq}; \pi_{q'q}), \quad \text{(SI11)}$$

next, renaming indices in the second term $i' = j'$ and $q' = l$ and because there is symmetry in $\boldsymbol{x}$ and $\boldsymbol{\pi}$ it allows us to add these two terms which yields

$$\frac{\partial}{\partial \tau_{iq}} \mathcal{J}(f^*(\boldsymbol{z};\boldsymbol{\tau});\boldsymbol{\alpha},\boldsymbol{\pi}) = \sum_{j'\neq i}^{n} \sum_{l'=1}^{Q} \tau_{j'l'} \log f(x_{ij'}|z_{iq}, z_{j'l'}; \pi_{ql'}) + \log \alpha_q - \log \tau_{iq} - 1. \tag{SI12}$$

Relabelling dummy indices, setting to zero and solving for $\tau_{iq}$ yields $\hat{\tau}_{iq} \propto \hat{\alpha}_q \prod_{j\neq i}^{n} \prod_{l=1}^{Q} \left[f(x_{ij}|z_{iq}, z_{jl}; \hat{\pi}_{ql})\right]^{\hat{\tau}_{jl}}$.

#### C. Derivation of Integrated Classification Likelihood (ICL) Criterion

The starting point in the ICL derivation is the integrated classification likelihood $f(\boldsymbol{x}, \boldsymbol{z}|\mathrm{m}_Q)$ which considers the evidence of the clustering in the data. Provided that the priors for model parameters can be factored as $p(\boldsymbol{\pi}, \boldsymbol{\alpha}|\mathrm{m}_Q) = p(\boldsymbol{\pi}|\mathrm{m}_Q)p(\boldsymbol{\alpha}|\mathrm{m}_Q)$, the integrated classification likelihood can be written as

$$\log f(\boldsymbol{x}, \boldsymbol{z}|\mathrm{m}_Q) = \log f(\boldsymbol{x}|\boldsymbol{z}, \mathrm{m}_Q) + \log f(\boldsymbol{z}|\mathrm{m}_Q), \tag{SI13}$$

Applying a Laplace approximation (Schwarz et al., 1978) for the the integral $\log f(\boldsymbol{x}|\boldsymbol{z}, \mathrm{m}_Q) = \int_{\boldsymbol{\Pi}} f(\boldsymbol{x}|\boldsymbol{z}, \mathrm{m}_Q, \boldsymbol{\pi}) p(\boldsymbol{\pi}|\mathrm{m}_Q) d\boldsymbol{\pi}$, we obtain

$$\begin{aligned}\log f(\boldsymbol{x}|\boldsymbol{z}, \mathrm{m}_Q) \approx & \log f(\boldsymbol{x}|\boldsymbol{z}, \mathrm{m}_Q, \hat{\boldsymbol{\pi}}) \\ & - \frac{Q(Q+1)}{4} \log \left[\frac{n(n-1)}{2}\right],\end{aligned} \tag{SI14}$$

where $Q(Q+1)/2$ denotes the number of parameters in $\boldsymbol{\pi}$ and $n(n-1)/2$ the number of data points in $\boldsymbol{x}$.

Similarly, for the remaining term, $\log f(\boldsymbol{z}|\mathrm{m}_Q)$, we obtain

$$\log f(\boldsymbol{z}|\mathrm{m}_Q) \approx \log f(\boldsymbol{z}|\mathrm{m}_Q, \hat{\boldsymbol{\alpha}}) - \frac{Q-1}{2} \log[n], \tag{SI15}$$

where $Q-1$ denotes total number of parameters in $\boldsymbol{\alpha}$ and $n$ is the total number of missing data $\boldsymbol{z}$. Combining Eq. (SI14) and (SI15), the ICL criterion, defined as an approximation of $\log f(\boldsymbol{x}, \boldsymbol{z}|\mathrm{m}_Q)$, is given as

$$\begin{aligned}\mathrm{ICL}(\mathrm{m}_Q) = & \log f(\boldsymbol{x}, \boldsymbol{z}|\mathrm{m}_Q, \hat{\boldsymbol{\pi}}, \hat{\boldsymbol{\alpha}}) - \frac{1}{2}\frac{Q(Q+1)}{2} \log[\frac{n(n-1)}{2}] \\ & - \frac{Q-1}{2} \log[n].\end{aligned} \tag{SI16}$$

Hence, the estimate of the number the blocks ($Q$) is found by selecting the model which maximises the ICL score.

#### D. Optimisation Algorithm for Bin-SBM

In this section, we discuss numerically stable way to evaluate $\boldsymbol{\tau}$ and then we spell out all the gory details regarding the optimisation algorithm for Bin-SBM.

##### *D.1. Numerical Stability of Cluster Assignment Probabilities*

Evaluating $\tau_{iq}$ directly in the probability domain may lead to some numerical issues. To avoid this, in practice, $\tau_{iq}$ is evaluated by using a logarithmic transformation on both sides of Eq. (10) and Eq. (18)), given in the main manuscript. For the simplicity of our discussion, let us concentrate only on Eq. (10) (the same ideas hold for Eq. (18). Therefore, we have

$$\begin{aligned}\log(\hat{\tau}_{iq}) &\propto \log(\hat{\alpha}_q) + \sum_{j\neq i}^{n}\sum_{l=1}^{Q} \hat{\tau}_{jl} \log(f(x_{ij}|z_{iq}, z_{jl}; \hat{\pi}_{ql})) \\ &\propto \log(\hat{\alpha}_q) + \sum_{j\neq i}^{n} \binom{K}{x_{ij}} + \sum_{j\neq i}^{n} x_{ij} \sum_{l=1}^{Q} \hat{\tau}_{jl} \big( \log[\hat{\pi}_{ql}] - \log[1-\hat{\pi}_{ql}]\big) \\ &\qquad + K \sum_{j\neq i}^{n}\sum_{l=1}^{Q} \hat{\tau}_{jl} \log[1-\hat{\pi}_{ql}]. \end{aligned} \tag{SI17}$$

Posterior cluster assignment probabilities are updated in the log domain using the numerically stable equation. Since the binomial coefficients are not depending on any of the model parameters they can be evaluated only once. When the updates for each node are completed, their cluster probabilities are normalised so that they sum to 1. Letting $t_{iq}$ to be the right hand side of the equation above

$$t_{iq} = \log(\hat{\alpha}_q) + \sum_{j\neq i}^{n} \binom{K}{x_{ij}} + \sum_{j\neq i}^{n} x_{ij} \sum_{l=1}^{Q} \hat{\tau}_{jl}\big(\log[\hat{\pi}_{ql}] - \log[1-\hat{\pi}_{ql}]\big) + \sum_{j\neq i}^{n} \binom{K}{x_{ij}} + K \sum_{j\neq i}^{n} \sum_{l=1}^{Q} \hat{\tau}_{jl} \log[1-\hat{\pi}_{ql}], \qquad \text{(SI18)}$$

a normalisation step is used to evaluate $\tau_{iq}$, wherein

$$\hat{\tau}_{iq} = \frac{\exp(t_{iq})}{\sum_{l=1}^{Q} \exp(t_{il})} = \frac{1}{\sum_{l=1}^{Q} \exp(t_{il} - t_{iq})}, \qquad \text{(SI19)}$$

where the first equality is numerically unstable while the last equality is numerically stable and consequently used in our implementation. We also bound the smallest probabilities to be of the order of $10^{-10}$ to avoid any further numerical instability caused when $\tau_{iq}$ is exactly equal to 0. To demonstrate that the implementation detailed above is numerically stable, let us consider the example with $Q = 10$, $n = 200$, in which for all $i$ and $q$, $F = 0.01$ (this value is assumed to be the maximum value of $f(x_{ij}|z_{iq}, z_{jl}; \pi_{ql})$ across all $i$ and $j$). In this instance, the evaluation of Eq. (10) in the probability domain would be numerically unstable as we would have, for the upper bound,

$$\hat{\tau}_{iq} \propto \frac{1}{Q} F^{n-1} = \frac{1}{Q}(0.01)^{199}, \qquad \text{(SI20)}$$

would be evaluated as 0. This indicates that Eq. (10) is numerically unstable. Utilising this approach all values of cluster probabilities would be zero, making it impossible to obtain a valid normalisation. Evaluating this in the log domain, we would have

$$\log(\hat{\tau}_{iq}) \propto 199 \times \log(0.01) = -918.7, \tag{SI21}$$

which is numerically stable. Subsequently, for the normalisation step and the conversion to the probability space, we can see that the first equality given above would be unstable as the numerator would be evaluated as 0 for each $q$. However, the last equality is numerically stable as the numerator is always 1 and the denominator is always a sum with at least an element equal to 1 (when $l = q$) and the other elements are numerically $\geq 0$.

*D.2. Variational EM algorithm for Binomial SBM (Bin-SBM)*

We show the algorithm for Bin-SBM as a pseudocode. We define operator $\Delta$ and $\delta$ which measure absolute and relative change in a parameter between two consecutive steps (e.g., $m$ and $m-1$) in the following way

$$\Delta\boldsymbol{\alpha} = \boldsymbol{\alpha}^{(m)} - \boldsymbol{\alpha}^{(m-1)} \tag{SI22}$$

$$\delta\boldsymbol{\alpha} = \frac{\Delta\boldsymbol{\alpha}}{\boldsymbol{\alpha}^{(m-1)}}. \tag{SI23}$$

**Algorithm 1:** Variational EM algorithm for Bin-SBM

**Input:** $\boldsymbol{x}$: is the adjacency data given as $n \times n$ matrix (see Eq. (3)), $Q$: is the total number of clusters, $M$: maximum number of M-steps, and $E$: maximum number of E-steps, $\epsilon_1$: is the convergence criterion in parameter updates for $\boldsymbol{\alpha}$ and $\boldsymbol{\pi}$, $\epsilon_2$: is convergence criterion in parameter updates for $\boldsymbol{\tau}$.

**Output:** Point estimates of $\boldsymbol{\tau}$, $\boldsymbol{\alpha}$, $\boldsymbol{\pi}$. Model selection criterion ICL score, VB which is a vector of length $M$ containing all the variational bound estimates.

`// Initialisation`

1. Use an algorithm (e.g. hierarchical clustering, k-means etc.) or random initialisation to estimate cluster labels $\boldsymbol{z}^{(0)}$.
2. Define $\boldsymbol{\tau}^{(0)}$ to be matrix of size $n \times Q$ populated entirely with $10^{-10}$.
3. Use these cluster labels to compute $\boldsymbol{\tau}^{(0)}$ by setting $\tau_{iq}^{(0)} = 1 - (Q-1) \times 10^{-10}$ when $z_{iq} = 1$ for a particular $q \in \{1, \ldots, Q\}$.
4. Set $m = 1$
5. Define $\boldsymbol{\alpha}$ to be a vector of length $Q$ and $\boldsymbol{\pi}$ to be a matrix $Q \times Q$.

`// Alternating between E-step and M-step`

**while** ($m \leq M$ *and* $\max(|\delta\boldsymbol{\alpha}|, |\delta\boldsymbol{\pi}|) > \epsilon_1$) **do**

   1. Update $\boldsymbol{\alpha}$ by Eq. (8)
   2. Update $\boldsymbol{\pi}$ by Eq. (9)
   3. Set $e = 1$
   4. **while** ($e \leq E$ *and* $\max(|\delta\boldsymbol{\tau}|) > \epsilon_2$) **do**

      4.1. Update $\boldsymbol{\tau}$ (see SI Section D.1 for details)

      4.2. e = e + 1

      **end**
   5. Compute VB for current $m$
   6. $m = m + 1$

**end**

1. Compute ICL.

#### E. Estimation in Homogeneous Stochastic Blockmodel (Hom-SBM)

In this model, we use the variational optimisation strategy to estimate the parameters $(\boldsymbol{\tau}, \boldsymbol{\alpha}, \boldsymbol{\theta}, \boldsymbol{\beta})$ with a Newton-Raphson algorithm for the parameters $(\boldsymbol{\theta}, \boldsymbol{\beta})$. In the continuation of this section, we assume that all block wise intercept parameters are collected into a $Q \times Q$ matrix $\boldsymbol{\theta}$. In addition to this, we also consider a Firth type estimation (Firth, 1993) for the intercept parameters in $\boldsymbol{\theta}$ but not for the regression coefficients. The reason for this is that for $\boldsymbol{\beta}$ estimates we are using all data points and thus it is less likely that we will run into bias issues. However, this is not the case for the intercept parameter $\boldsymbol{\theta}$ which depends on the number of data points in a block. Now, given that the multi-subject SBMs pose no restriction on the size of individual blocks it can easily happen that some smaller blocks may have connection rate estimates that are close to 1 or 0 which makes inferences invalid. Nevertheless, this type of bias can be removed with the Firth type estimates.

As for the other two multi-subject SBMs, the starting point for the estimation of the parameters is the variational bound,

$$\begin{aligned} \mathcal{J}(f^*(\boldsymbol{z}; \boldsymbol{\tau}); \boldsymbol{\alpha}, \boldsymbol{\theta}, \boldsymbol{\beta}) &= \sum_{i=1}^{n} \sum_{q=1}^{Q} \tau_{iq} \log[\alpha_q] - \sum_{i=1}^{n} \sum_{q=1}^{Q} \tau_{iq} \log[\tau_{iq}] \\ &+ \frac{1}{2} \sum_{k=1}^{K} \sum_{i=1}^{n} \sum_{j \neq i}^{n} \sum_{q,l}^{Q} \tau_{iq} \tau_{jl} \log[f(x_{ijk} | z_{iq}, z_{jl}; \theta_{ql}, \boldsymbol{\beta})]. \end{aligned} \tag{SI24}$$

Similarly to the previous multi-subject models, we obtain the point estimating equations for $\hat{\boldsymbol{\tau}}$ and $\hat{\boldsymbol{\alpha}}$ by maximising the variational bound in Eq. (SI24), which yields

$$\hat{\tau}_{iq} \propto \hat{\alpha}_q \prod_{k=1}^{K} \prod_{j \neq i}^{n} \prod_{l=1}^{Q} \left[ f(x_{ijk} | z_{iq}, z_{jl}; \theta_{ql}, \boldsymbol{\beta}) \right]^{\hat{\tau}_{jl}}, \tag{SI25}$$

$$\hat{\alpha}_q = \frac{1}{n} \sum_{i=1}^{n} \hat{\tau}_{iq}. \tag{SI26}$$

In order to obtain the estimates of the parameters in the logistic regression model $(\boldsymbol{\theta}, \boldsymbol{\beta})$, we use the Newton-Raphson algorithm. In what follows next, we first fix a general notation and we state the implicit forms of various preliminary quantities that are needed for the subsequent discussion. Thus, we note that the first order partial derivatives of the variational bound $\mathcal{J}(f^*(\boldsymbol{z}; \boldsymbol{\tau}); \boldsymbol{\alpha}, \boldsymbol{\theta}, \boldsymbol{\beta})$ with respect to $\theta_{ql}$ and $\boldsymbol{\beta}$ as $\mathrm{U}(\theta_{ql})$ and $\mathbf{U}(\boldsymbol{\beta})$ and note that they are explicitly given as

$$\mathrm{U}(\theta_{ql}) = \frac{1}{2} \sum_{k=1}^{K} \sum_{i \neq j}^{n} \sum_{j \neq i}^{n} \tau_{iq} \tau_{jl} (x_{ijk} - \pi_{qlk}), \tag{SI27}$$

$$\mathbf{U}(\boldsymbol{\beta}) = \frac{1}{2} \sum_{k=1}^{K} \sum_{i \neq j}^{n} \sum_{j \neq i}^{n} \sum_{q,l}^{Q} \tau_{iq} \tau_{jl} (x_{ijk} - \pi_{qlk}) \boldsymbol{d}_k. \tag{SI28}$$

which is the $(Q(Q+1)/2+P) \times (Q(Q+1)/2+P)$ matrix, and it is related to the $Q(Q+1)/2$ intercept parameters in $\boldsymbol{\theta}$ and the $P$ regression coefficients in $\boldsymbol{\beta}$. Similarly, we note the negative second order partial derivatives of the variational bound yield the observed Fisher Information matrix $\boldsymbol{\mathcal{I}}(\boldsymbol{\theta}, \boldsymbol{\beta})$. In particular, $\boldsymbol{\mathcal{I}}(\boldsymbol{\theta}, \boldsymbol{\beta})$ consists of three sub-matrices

$$\boldsymbol{\mathcal{I}}(\boldsymbol{\theta}, \boldsymbol{\beta}) = \left[\begin{array}{c|c} \boldsymbol{\mathcal{I}}_1(\boldsymbol{\theta}) & \boldsymbol{\mathcal{I}}_2^\top(\boldsymbol{\theta}, \boldsymbol{\beta}) \\ \hline \boldsymbol{\mathcal{I}}_2(\boldsymbol{\theta}, \boldsymbol{\beta}) & \boldsymbol{\mathcal{I}}_3(\boldsymbol{\beta}) \end{array}\right], \tag{SI29}$$

such that

- $\boldsymbol{\mathcal{I}}_1(\boldsymbol{\theta})$ is a diagonal $Q(Q+1)/2 \times Q(Q+1)/2$ matrix,
- $\boldsymbol{\mathcal{I}}_2(\boldsymbol{\theta}, \boldsymbol{\beta})$ is a $P \times Q(Q+1)/2$ matrix, and
- $\boldsymbol{\mathcal{I}}_3(\boldsymbol{\beta})$ is a $P \times P$ matrix.

Below, we give their specific elements

$$\mathcal{I}_1(\theta_{ql}, \theta_{q'l'}) = \begin{cases} \frac{1}{2}\sum_{k=1}^{K}\sum_{i=1}^{n}\sum_{j\neq i}^{n} \tau_{iq}\tau_{jl}\pi_{qlk}(1-\pi_{qlk}) & \text{if } q, l = q', l' \\ 0 & \text{otherwise,} \end{cases} \tag{SI30}$$

$$\boldsymbol{\mathcal{I}}_2(\theta_{ql}, \boldsymbol{\beta}) = \frac{1}{2}\sum_{k=1}^{K}\sum_{i=1}^{n}\sum_{j\neq i}^{n} \tau_{iq}\tau_{jl}\pi_{qlk}(1-\pi_{qlk})\boldsymbol{d}_k \tag{SI31}$$

$$\boldsymbol{\mathcal{I}}_3(\boldsymbol{\beta}) = \frac{1}{2}\sum_{k=1}^{K}\sum_{i=1}^{n}\sum_{j\neq i}^{n}\sum_{q,l}^{Q} \tau_{iq}\tau_{jl}\pi_{qlk}(1-\pi_{qlk})\ \boldsymbol{d}_k\boldsymbol{d}_k^\top. \tag{SI32}$$

However, in order to obtain the Firth type estimates of $\boldsymbol{\theta}$, we want to maximise a modified variational bound $\mathcal{J}^*(f^*(\boldsymbol{z};\boldsymbol{\tau});\boldsymbol{\alpha},\boldsymbol{\theta},\boldsymbol{\beta})$ given as

$$\mathcal{J}^*(f^*(\boldsymbol{z};\boldsymbol{\tau});\boldsymbol{\alpha},\boldsymbol{\theta},\boldsymbol{\beta}) = \mathcal{J}(f^*(\boldsymbol{z};\boldsymbol{\tau});\boldsymbol{\alpha},\boldsymbol{\theta},\boldsymbol{\beta}) + \frac{1}{2}\log\left[\text{Det}(\boldsymbol{\mathcal{I}}(\boldsymbol{\theta},\boldsymbol{\beta}))\right], \tag{SI33}$$

whose partial derivatives $U^*(\theta_{ql})$ are given as

$$U^*(\theta_{ql}) = U(\theta_{ql}) + \frac{1}{2}\mathrm{Tr}\left[\boldsymbol{\mathcal{I}}^{-1}(\boldsymbol{\theta},\boldsymbol{\beta})\frac{\partial}{\partial\theta_{ql}}\,\boldsymbol{\mathcal{I}}(\boldsymbol{\theta},\boldsymbol{\beta})\right], \tag{SI34}$$

and the derivatives of the observed Fisher Information matrix are given as

$$\frac{\partial \mathcal{I}_1(\theta_{q'l'},\theta_{q''l''})}{\partial\theta_{ql}} = \begin{cases} \frac{1}{2}\sum_{k=1}^{K}\sum_{i=1}^{n}\sum_{j\neq i}^{n}\tau_{iq}\tau_{jl}\pi_{qlk}(1-\pi_{qlk})(1-2\pi_{qlk}) & q,l=q',l'=q'',l'' \\ 0 & \text{otherwise} \end{cases} \tag{SI35}$$

$$\frac{\partial \boldsymbol{\mathcal{I}}_2(\boldsymbol{\theta},\boldsymbol{\beta})}{\partial\theta_{ql}} = \frac{1}{2}\sum_{k=1}^{K}\sum_{i=1}^{n}\sum_{j\neq i}^{n}\tau_{iq}\tau_{jl}\pi_{qlk}(1-\pi_{qlk})(1-2\pi_{qlk})\boldsymbol{d}_k, \tag{SI36}$$

$$\frac{\partial \boldsymbol{\mathcal{I}}_3(\boldsymbol{\beta})}{\partial\theta_{ql}} = \frac{1}{2}\sum_{k=1}^{K}\sum_{i=1}^{n}\sum_{j\neq i}^{n}\tau_{iq}\tau_{jl}\pi_{qlk}(1-\pi_{qlk})(1-2\pi_{qlk})\boldsymbol{d}_k\boldsymbol{d}_k^{\top}. \tag{SI37}$$

Finally, the estimating equations for $\boldsymbol{\theta}$ and $\boldsymbol{\beta}$ are found using the Fisher Scoring formula which, for the $(r)$-th step, updates the parameters according to the following expression

$$\begin{pmatrix}\boldsymbol{\theta}\\ \boldsymbol{\beta}\end{pmatrix}^{(r)} = \begin{pmatrix}\boldsymbol{\theta}\\ \boldsymbol{\beta}\end{pmatrix}^{(r-1)} + \boldsymbol{\mathcal{I}}^{-1}(\boldsymbol{\theta}^{(r-1)},\boldsymbol{\beta}^{(r-1)})\begin{pmatrix}\mathbf{U}^*(\boldsymbol{\theta})\\ \mathbf{U}(\boldsymbol{\beta})\end{pmatrix}^{(r-1)}, \tag{SI38}$$

where $\boldsymbol{\theta}$ is vectorised and have length $Q(Q+1)/2$ and $\mathbf{U}^*(\boldsymbol{\theta})$ is the column vector of length $Q(Q+1)/2$ whose elements are given by Eq. (SI34). It is important to highlight that the $\boldsymbol{\theta}$ parameters are based on the Firth type

MLEs and thus they are updated with $\mathbf{U}^*(\boldsymbol{\theta})$, while the $\boldsymbol{\beta}$ parameters are based on the Ordinary MLEs and thus are updated with $\mathbf{U}(\boldsymbol{\beta})$. Also, it is possible to obtain the Ordinary MLEs for $\boldsymbol{\theta}$ by simply using $\mathbf{U}(\boldsymbol{\theta})$ in Eq. (SI38).

Finally, the ICL criterion is given as

$$\begin{aligned}\mathrm{ICL}(\mathrm{m}_Q) = &\log f(\boldsymbol{x}, \hat{\boldsymbol{z}}|\mathrm{m}_Q, \hat{\boldsymbol{\alpha}}, \hat{\boldsymbol{\theta}}, \hat{\boldsymbol{\beta}}) \\ &- \frac{1}{2}\left(\frac{Q(Q+1)}{2} + P\right)\log\left[\frac{n(n-1)}{2}K\right] \\ &- \frac{Q-1}{2}\log[n]. \end{aligned} \qquad \text{(SI39)}$$

*Inference for Hom-SBM*

In the general context of logistic regression analysis, the Wald test has been commonly used to make inferences on the estimates of the regression coefficients. For the Hom-SBM, we write the set of parameters $\{\boldsymbol{\theta}, \boldsymbol{\beta}\} = \boldsymbol{\psi}$ as a column vector. To perform inference on a combination of the parameters, $\mathcal{H}_0 : \boldsymbol{L}\boldsymbol{\psi} = \boldsymbol{b}_0$, where $\boldsymbol{L}$ is a matrix (or a vector) which defines the combination of the parameters (i.e. linear contrast) tested and $\boldsymbol{b}_0$ are some constants under the null hypothesis, the Wald statistic is

$$W = (\boldsymbol{L}\hat{\boldsymbol{\psi}} - \boldsymbol{b}_0)^\top(\boldsymbol{L}\boldsymbol{\mathcal{I}}^{-1}(\hat{\boldsymbol{\psi}})\boldsymbol{L}^\top)^{-1}(\boldsymbol{L}\hat{\boldsymbol{\psi}} - \boldsymbol{b}_0)/c, \qquad \text{(SI40)}$$

where $c$ is the rank of $\boldsymbol{L}$. Asymptotically, $W$ follows a $\chi^2_c$ distribution. Alternatively, if $\boldsymbol{L}$ is a vector, then the Wald statistic can be noted as

$$W^* = \frac{\boldsymbol{L}\hat{\boldsymbol{\psi}} - \boldsymbol{b}_0}{\sqrt{\boldsymbol{L}\boldsymbol{\mathcal{I}}^{-1}(\hat{\boldsymbol{\psi}})\boldsymbol{L}^\top}}, \tag{SI41}$$

which asymptotically follows a Standard Normal distribution. The standard errors of the model parameters depend on the observed Fisher Information matrix given by Eq. (SI29). Note that Eq. (SI29) applies for the Ordinary and Firth MLEs (Firth, 1993; Heinze and Schemper, 2002).

*Likelihood ratio test*

In the case of the Hom-SBM, to perform inference on a combination of parameters $\mathcal{H}_0 : \boldsymbol{L}\boldsymbol{\psi} = \boldsymbol{b}_0$, we can use the likelihood ratio (LR) test. For the Firth approach, we base the likelihood ratio test statistic on the modified variational bound given by Eq. (SI33), and thus the test statistic can be stated as

$$\Lambda = 2\Big[\mathcal{J}^*(f^*(\boldsymbol{z};\hat{\boldsymbol{\tau}});\hat{\boldsymbol{\alpha}},\hat{\boldsymbol{\psi}}) - \mathcal{J}^*(f^*(\boldsymbol{z};\hat{\boldsymbol{\tau}});\hat{\boldsymbol{\alpha}},\tilde{\boldsymbol{\psi}})\Big], \tag{SI42}$$

where $\tilde{\boldsymbol{\psi}}$ is a column vector containing the parameter estimates under the null hypothesis. In particular, $\Lambda$ is assumed to follow a $\chi^2_c$ distribution where $c$ is the rank of $\boldsymbol{L}$. Note that the $\tilde{\boldsymbol{\psi}}$ is derived from a restricted model whose penalisation term is different from the penalisation term associated with the full model. Hence, to derive $\Lambda$, we do not use the bound associated with the restricted model, but we use instead the bound of the full model evaluated with $\tilde{\boldsymbol{\psi}}$.

#### F. Optimisation algorithm for Heterogeneous SBM (Het-SBM)

In this section, we present the pseudocode for the Het-SBM which is given in terms of two algorithms. The first algorithm is the main variational EM algorithm and the second algorithm shows the optimisation procedure for $\boldsymbol{\beta}$. To avoid using extra indices, we set $P$ to denote the total number of covariates ($P^*$) plus 1 for the intercept term (i.e. $P = P^* + 1$). Also, we set $\Delta$ to be an operator defining the absolute change in parameters between two consecutive steps (e.g., fitted values at $r$ and $r-1$), so that

$$\Delta\boldsymbol{\beta}_{ql} = \boldsymbol{\beta}_{ql}^{(r)} - \boldsymbol{\beta}_{ql}^{(r-1)}, \tag{SI43}$$

and $\delta$ to be operator defining the relative rate of change

$$\delta\boldsymbol{\beta}_{ql} = \frac{\Delta\boldsymbol{\beta}_{ql}}{\boldsymbol{\beta}_{ql}^{(r-1)}}. \tag{SI44}$$

We also define block-wise bound ($\mathcal{J}^*_{ql}$) with respect to $\boldsymbol{\beta}_{ql}$ to be

$$\mathcal{J}^*_{ql} = \frac{1}{2}\sum_{k=1}^{K}\sum_{i=1}^{n}\sum_{j\neq i}^{n}\gamma_{ijql}\log f(x_{ij}|z_{iq}, z_{jl};\boldsymbol{\beta}_{ql}) + \frac{1}{2}\log[\mathrm{Det}(\boldsymbol{\mathcal{I}}_{\boldsymbol{ql}}(\boldsymbol{\beta}_{ql})]. \tag{SI45}$$

**Algorithm 2:** Variational EM algorithm for Het-SBM

**Input:** $\boldsymbol{x}$: adjacency data given as an $n \times n \times K$ tensor, $\boldsymbol{d}$: an $K \times P$ design matrix, $Q$: number of clusters, $M$: maximum number of M-steps, $E$: maximum number of E-steps, $\epsilon_1$: convergence criterion in parameter updates for $\boldsymbol{\alpha}$, $\boldsymbol{\beta}$, $\epsilon_2$: convergence criterion in parameter updates for $\boldsymbol{\tau}$.

**Output:** Point estimates of $\boldsymbol{\tau}$, $\boldsymbol{\alpha}$, $\boldsymbol{\beta}$, Fisher Information $\boldsymbol{\mathcal{I}}$, ICL score and VB which is a vector of length $M$ containing all the variational bound estimates.

`// Initialisation`

1. Use an algorithm (e.g. hierarchical clustering, k-means etc.) or a random initialisation to estimate the cluster labels $\boldsymbol{z}^{(0)}$.
2. Set $\boldsymbol{\tau}$ to be a matrix of size $n \times Q$ such that $\tau_{iq}$ is equal to $1-(Q-1)\times 10^{-10}$ when $z_{iq}=1$, or $10^{-10}$ otherwise.
3. Set $\boldsymbol{\alpha}$ to be a zero vector of length $Q$
4. Set $\boldsymbol{\beta}$ to be a zero tensor of size $Q \times Q \times P$ .
5. Set $m = 1$

`// Alternating between E-step and M-step`

**while** ($m \leq M$ *and* $\max(|\delta\boldsymbol{\beta}|, |\delta\boldsymbol{\alpha}|) > \epsilon_1$) **do**

  1. Update $\boldsymbol{\alpha}$ using Eq. (17)
  2. Update $\boldsymbol{\beta}$ using Algorithm 3
  3. Set $e = 1$
  4. **while** ($e \leq E$ *and* $\max(|\delta\boldsymbol{\tau}| > \epsilon_2)$ **do**
     - 4.1. Update $\boldsymbol{\tau}$ (see SI Section D.1 for details)
     - 4.2. e = e + 1

     **end**
  5. Compute VB for current $m$
  6. $m = m + 1$

**end**

1. Compute ICL.

**Algorithm 3:** Het-SBM implementation of the Firth regression step

**Input:** $\boldsymbol{\beta}$, $\boldsymbol{\tau}$, $\boldsymbol{d}$, $\boldsymbol{x}$, $\epsilon = 10^{-10}$, $R = 10$, $\Delta^{\max} = 5$, $H = 5$

**Output:** $\boldsymbol{\beta}$, $\boldsymbol{\mathcal{I}}(\boldsymbol{\beta})$, $\mathcal{J}^*_{ql}$

**for** $(q \in \{1, \dots, Q\})$ **do**

  **for** $(l \in \{q, \dots, Q\})$ **do**

    Compute $\mathcal{J}^*_{ql}(\boldsymbol{\beta}_{ql})$

    $r = 1$, $\Delta\mathcal{J}^*_{ql} = 1$

    **while** $(r \leq R$ *and* $\Delta\mathcal{J}^*_{ql} > \epsilon)$ **do**

      Compute $\boldsymbol{\mathcal{I}_{ql}(\beta_{ql})}$, $\frac{\partial \boldsymbol{\mathcal{I}_{ql}}(\boldsymbol{\beta}_{ql})}{\partial \boldsymbol{\beta}_{ql}}$, $\mathbf{U}^*_{ql}(\boldsymbol{\beta}_{ql})$ and $\Delta\boldsymbol{\beta}_{ql}$ (Eq. (SI43))

      Clamp the elements of $\Delta\boldsymbol{\beta}_{ql}$ to be in $[-\Delta^{\max}, \Delta^{\max}]$

      Compute $\Delta\mathcal{J}^*_{ql} = \mathcal{J}^*_{ql}(\boldsymbol{\beta}_{ql} + \Delta\boldsymbol{\beta}_{ql}) - \mathcal{J}^*_{ql}(\boldsymbol{\beta}_{ql})$

      `// Step halving procedure`

      $h = 1$

      **while** $(h \leq H$ *and* $\Delta\mathcal{J}^*_{ql} < 0)$ **do**

        $\Delta\boldsymbol{\beta}_{ql} = \Delta\boldsymbol{\beta}_{ql}/2$

        Compute $\Delta\mathcal{J}^*_{ql} = \mathcal{J}^*_{ql}(\boldsymbol{\beta}_{ql} + \Delta\boldsymbol{\beta}_{ql}) - \mathcal{J}^*_{ql}(\boldsymbol{\beta}_{ql})$

        $h = h + 1$

      **end**

      **if** $(\Delta\mathcal{J}^*_{ql} < 0)$ **then**

        $\Delta\boldsymbol{\beta}_{ql} = 0$

      **end**

      $\boldsymbol{\beta}_{ql} = \boldsymbol{\beta}_{ql} + \Delta\boldsymbol{\beta}_{ql}$

    **end**

    Compute $\mathcal{J}^*_{ql}(\boldsymbol{\beta}_{ql})$ and $\boldsymbol{\mathcal{I}_{ql}(\beta_{ql})}$

  **end**

**end**

#### G. Evaluation methods

To measure the similarity between the true partitions and the partitions estimated by the multi-subject models, we use the Adjusted Rand Index (ARI) (Handl et al., 2005; Hubert and Arabie, 1985). This quantity is a modification of the Rand Index (RI) (Rand, 1971), that is expressed as the fraction of node pairs that are consistent: a node pair is consistent between two partitions if either (a) the node pair is within the same block in both partitions, or (b) the node pair is split between two blocks in both partitions. The interpretation of the RI depends on the number of blocks (Morey and Agresti, 1984), whereas the ARI is adjusted for the chance agreement and the number of blocks (Hubert and Arabie, 1985). It is defined as

$$\mathrm{ARI} = \frac{\mathrm{RI} - \mathbb{E}(\mathrm{RI})}{\max(\mathrm{RI}) - \mathbb{E}(\mathrm{RI})}, \tag{SI46}$$

where the expectation is computed assuming a hypergeometric distribution of the counts of consistent node pairs. The ARI scores range from 0 to 1, and indicate the proportion of overlap; for example, if two partitions have an ARI score of 0.6, this means that 60% of the nodes are classified in the same blocks.

#### H. Results for Simulation I

| $\hat{Q}$ | Hom-Modular | | |
|---|---|---|---|
| | Balanced | M. Unbalanced | Unbalanced |
| 2 | [-108071, -106896] | [-109322, -108028] | [-98724, -97407] |
| 3 | [-88108, -87011] | [-74162, -72985] | [-49380, -48089] |
| 4 | [-72541, -71567] | [-49423, -48534] | [-24706, -23950] |
| 5 | [-59529, -57472] | [-31911, -31104] | [-12977, -12361] |
| 6 | [-46548, -45770] | [-20322, -19501] | [-6575, -6200] |
| 7 | [-34857, -34235] | [-12315, -11656] | [-3250, -2983] |
| 8 | [-23171, -22760] | [-6546, -6160] | [-1594, -1369] |
| 9 | [-11531, -11268] | [-2594, -2413] | [-655, -543] |
| 10 | [0, 0] | [0, 0] | [0, 0] |
| 11 | [-70, -59] | [-76, -59] | [-101, -61] |
| 12 | [-147, -131] | [-155, -134] | [-180, -143] |
| 13 | [-229, -210] | [-238, -213] | [-271, -221] |
| 14 | [-318, -295] | [-329, -298] | [-363, -298] |
| 15 | [-412, -384] | [-422, -388] | [-446, -401] |
| 16 | [-511, -484] | [-518, -489] | [-537, -501] |
| 17 | [-620, -577] | [-626, -584] | [-657, -604] |
| 18 | [-730, -698] | [-739, -696] | [-768, -708] |

Table S1: **Ranges of adjusted ICL scores for Het-SBM in the scenarios with Hom-Modular cluster structure.** For each simulated dataset and a given $\hat{Q}$, the adjusted ICL scores are given as the difference between the maximum score across the restarts (local maximum) and the maximum score across all values of $\hat{Q}$ (global maximum). For each $\hat{Q}$, the range of ICL scores is shown across all 100 Monte Carlo samples. The range $[0, 0]$ indicates that for all 100 Monte Carlo samples, the best ICL score systematically pointed towards the correct number of clusters (i.e. $\hat{Q} = 10$). The ICL scores tends to decrease when $\hat{Q}$ is taking values that are going further away from the ground truth $\hat{Q} = 10$ with a greater rate for values below 10.

| $\hat{Q}$ | Het-Modular | | |
|---|---|---|---|
| | Balanced | M. Unbalanced | Unbalanced |
| 2 | [-41412, -40287] | [-44590, -43250] | [-42133, -40883] |
| 3 | [-34439, -32992] | [-31513, -30611] | [-23043, -22150] |
| 4 | [-28210, -27127] | [-23158, -21791] | [-14296, -13575] |
| 5 | [-22866, -21790] | [-16662, -15906] | [-10163, -9337] |
| 6 | [-17648, -16209] | [-12583, -11715] | [-6311, -5858] |
| 7 | [-12334, -11512] | [-8271, -7471] | [-3567, -3206] |
| 8 | [-8184, -7575] | [-4901, -4387] | [-2007, -1742] |
| 9 | [-4115, -3629] | [-1734, -1447] | [-833, -504] |
| 10 | [0, 0] | [0, 0] | [0, 0] |
| 11 | [-76, -62] | [-75, -58] | [-82, -61] |
| 12 | [-152, -131] | [-155, -134] | [-158, -136] |
| 13 | [-235, -207] | [-237, -211] | [-242, -217] |
| 14 | [-322, -297] | [-326, -300] | [-329, -300] |
| 15 | [-419, -386] | [-424, -387] | [-424, -394] |
| 16 | [-520, -483] | [-523, -485] | [-534, -495] |
| 17 | [-623, -594] | [-628, -590] | [-634, -601] |
| 18 | [-732, -690] | [-741, -696] | [-753, -700] |

Table S2: **Ranges of adjusted ICL scores for Het-SBM in the scenarios with Het-Modular cluster structure.** For each simulated dataset and a given $\hat{Q}$, the adjusted ICL scores are given as the difference between the maximum score across the restarts (local maximum) and the maximum score across all values of $\hat{Q}$ (global maximum). For each $\hat{Q}$, the range of ICL scores is shown across all 100 Monte Carlo samples. The range $[0, 0]$ indicates that for all 100 Monte Carlo samples, the best ICL score systematically pointed towards the correct number of clusters (i.e. $\hat{Q} = 10$). The ICL scores tends to decrease when $\hat{Q}$ is taking values that are going further away from the ground truth $\hat{Q} = 10$ with a greater rate for values below 10.

| $\hat{Q}$ | Hom-Modular | | |
|---|---|---|---|
| | Balanced | M. Unbalanced | Unbalanced |
| 2 | [-108159, -106985] | [-109410, -108116] | [-98812, -97496] |
| 3 | [-88191, -87094] | [-74246, -73069] | [-49464, -48172] |
| 4 | [-72618, -71644] | [-49500, -48611] | [-24783, -24027] |
| 5 | [-59597, -57540] | [-31980, -31172] | [-13045, -12429] |
| 6 | [-46606, -45828] | [-20380, -19559] | [-6633, -6258] |
| 7 | [-34903, -34281] | [-12361, -11702] | [-3297, -3030] |
| 8 | [-23203, -22792] | [-6579, -6192] | [-1627, -1402] |
| 9 | [-11548, -11285] | [-2611, -2430] | [-672, -561] |
| 10 | [0, 0] | [0, 0] | [0, 0] |
| 11 | [-52, -40] | [-57, -40] | [-82, -42] |
| 12 | [-108, -85] | [-116, -95] | [-141, -101] |
| 13 | [-168, -147] | [-176, -148] | [-206, -166] |
| 14 | [-234, -205] | [-237, -212] | [-269, -213] |
| 15 | [-302, -275] | [-310, -272] | [-335, -292] |
| 16 | [-373, -343] | [-377, -345] | [-398, -363] |
| 17 | [-452, -409] | [-457, -417] | [-485, -438] |
| 18 | [-529, -490] | [-539, -504] | [-564, -511] |

Table S3: **Ranges of adjusted ICL scores for Bin-SBM in the scenarios with Hom-Modular cluster structure.** For each simulated dataset and a given $\hat{Q}$, the adjusted ICL scores are given as the difference between the maximum score across the restarts (local maximum) and the maximum score across all values of $\hat{Q}$ (global maximum). For each $\hat{Q}$, the range of ICL scores is shown across all 100 Monte Carlo samples. The range $[0, 0]$ indicates that for all 100 Monte Carlo samples, the best ICL score systematically pointed towards the correct number of clusters (i.e. $\hat{Q} = 10$). The ICL scores tends to decrease when $\hat{Q}$ is taking values that are going further away from the ground truth $\hat{Q} = 10$ with a greater rate for values below 10.

| | Het-Modular | | |
|---|---|---|---|
| $\hat{Q}$ | Balanced | M. Unbalanced | Unbalanced |
| 2 | [-41500, -40376] | [-44679, -43339] | [-42221, -40971] |
| 3 | [-34523, -33075] | [-31596, -30694] | [-23126, -22234] |
| 4 | [-28286, -27204] | [-23234, -21867] | [-14373, -13651] |
| 5 | [-22934, -21858] | [-16730, -15974] | [-10231, -9405] |
| 6 | [-17706, -16267] | [-12640, -11773] | [-6369, -5916] |
| 7 | [-12380, -11558] | [-8317, -7517] | [-3613, -3252] |
| 8 | [-8216, -7607] | [-4933, -4419] | [-2039, -1774] |
| 9 | [-4132, -3646] | [-1751, -1464] | [-850, -521] |
| 10 | [0, 0] | [0, 0] | [0, 0] |
| 11 | [-54, -42] | [-56, -40] | [-63, -42] |
| 12 | [-113, -92] | [-115, -91] | [-119, -99] |
| 13 | [-171, -146] | [-174, -150] | [-179, -156] |
| 14 | [-237, -212] | [-241, -215] | [-244, -215] |
| 15 | [-308, -273] | [-314, -274] | [-314, -283] |
| 16 | [-379, -344] | [-383, -347] | [-390, -357] |
| 17 | [-457, -425] | [-458, -424] | [-465, -430] |
| 18 | [-534, -492] | [-543, -499] | [-550, -510] |

Table S4: **Ranges of adjusted ICL scores for Bin-SBM in the scenarios with Het-Modular cluster structure.** For each simulated dataset and a given $\hat{Q}$, the adjusted ICL scores are given as the difference between the maximum score across the restarts (local maximum) and the maximum score across all values of $\hat{Q}$ (global maximum). For each $\hat{Q}$, the range of ICL scores is shown across all 100 Monte Carlo samples. The range $[0, 0]$ indicates that for all 100 Monte Carlo samples, the best ICL score systematically pointed towards the correct number of clusters (i.e. $\hat{Q} = 10$). The ICL scores tends to decrease when $\hat{Q}$ is taking values that are going further away from the ground truth $\hat{Q} = 10$ with a greater rate for values below 10.

| $\hat{Q}$ | Core-Modular | | |
|---|---|---|---|
| | Balanced | M. Unbalanced | Unbalanced |
| 2 | [-31340, -30282] | [-44972, -43491] | [-51364, -49396] |
| 3 | [-23705, -22886] | [-24348, -23283] | [-22201, -21102] |
| 4 | [-18765, -17250] | [-16565, -15575] | [-10900, -10171] |
| 5 | [-14186, -12670] | [-10911, -10104] | [-6812, -5934] |
| 6 | [-9994, -9049] | [-6970, -6376] | [-3115, -2761] |
| 7 | [-6342, -5695] | [-4582, -3612] | [-1873, -1593] |
| 8 | [-3398, -3018] | [-2364, -2024] | [-876, -707] |
| 9 | [-991, -780] | [-537, -352] | [-645, -157] |
| 10 | [0, 0] | [0, 0] | [0, 0] |
| 11 | [-58, -44] | [-58, -44] | [-56, -44] |
| 12 | [-124, -98] | [-116, -95] | [-117, -100] |
| 13 | [-181, -154] | [-177, -153] | [-183, -160] |
| 14 | [-241, -215] | [-243, -215] | [-249, -230] |
| 15 | [-314, -278] | [-316, -284] | [-322, -297] |
| 16 | [-383, -344] | [-393, -357] | [-402, -375] |
| 17 | [-461, -427] | [-474, -435] | [-490, -440] |
| 18 | [-542, -506] | [-560, -506] | [-572, -529] |

Table S5: **Ranges of adjusted ICL scores for Bin-SBM in the scenarios with Core-Modular cluster structure.** For each simulated dataset and a given $\hat{Q}$, the adjusted ICL scores are given as the difference between the maximum score across the restarts (local maximum) and the maximum score across all values of $\hat{Q}$ (global maximum). For each $\hat{Q}$, the range of ICL scores is shown across all 100 Monte Carlo samples. The range $[0, 0]$ indicates that for all 100 Monte Carlo samples, the best ICL score systematically pointed towards the correct number of clusters (i.e. $\hat{Q} = 10$). The ICL scores tends to decrease when $\hat{Q}$ is taking values that are going further away from the ground truth $\hat{Q} = 10$ with a greater rate for values below 10.

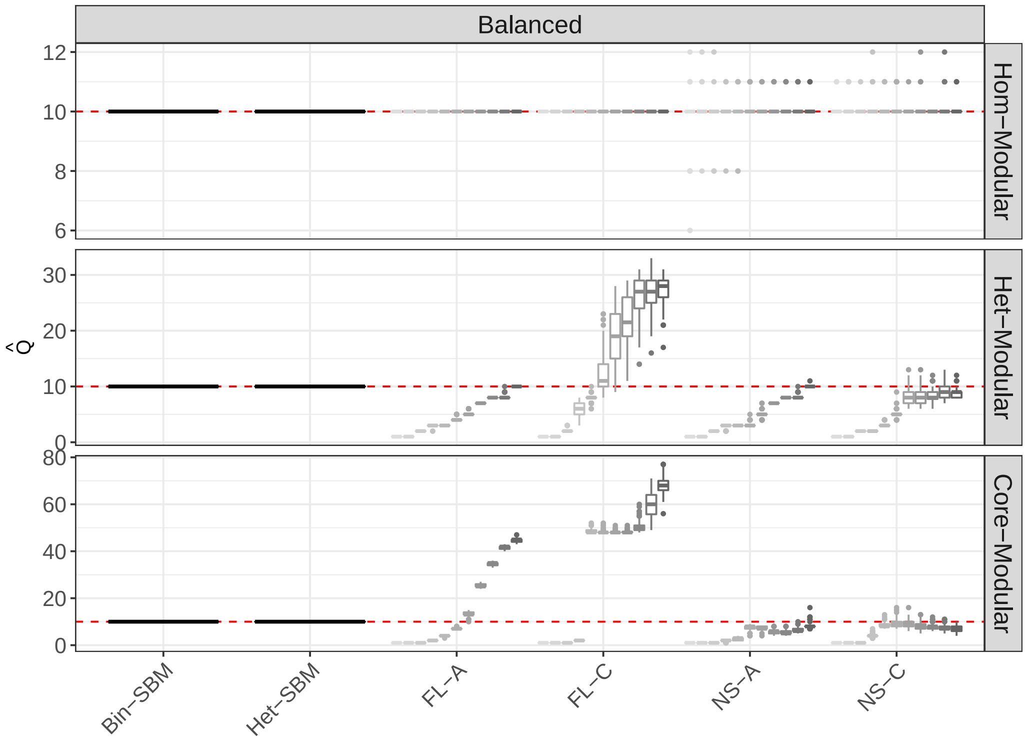


Figure S1: **Box-plots of estimated $\hat{Q}$ over 100 Monte Carlo samples in the Balanced scenarios**. The $x$-axis shows MS-SBMs and the modular algorithms, whose abbreviations FL-A and FL-C stand for the Fast Louvain algorithm with average & consensus clustering while the abbreviations NS-A and NS-C stand for the Newman Spectral algorithm with average & consensus clustering. For each modular algorithm, there are 11 box-plots whose colours correspond to the solutions with different $\gamma$ values starting from 0.5 and finishing at 1.5 with a step-size of 0.1 and the central score of 1 (default value).

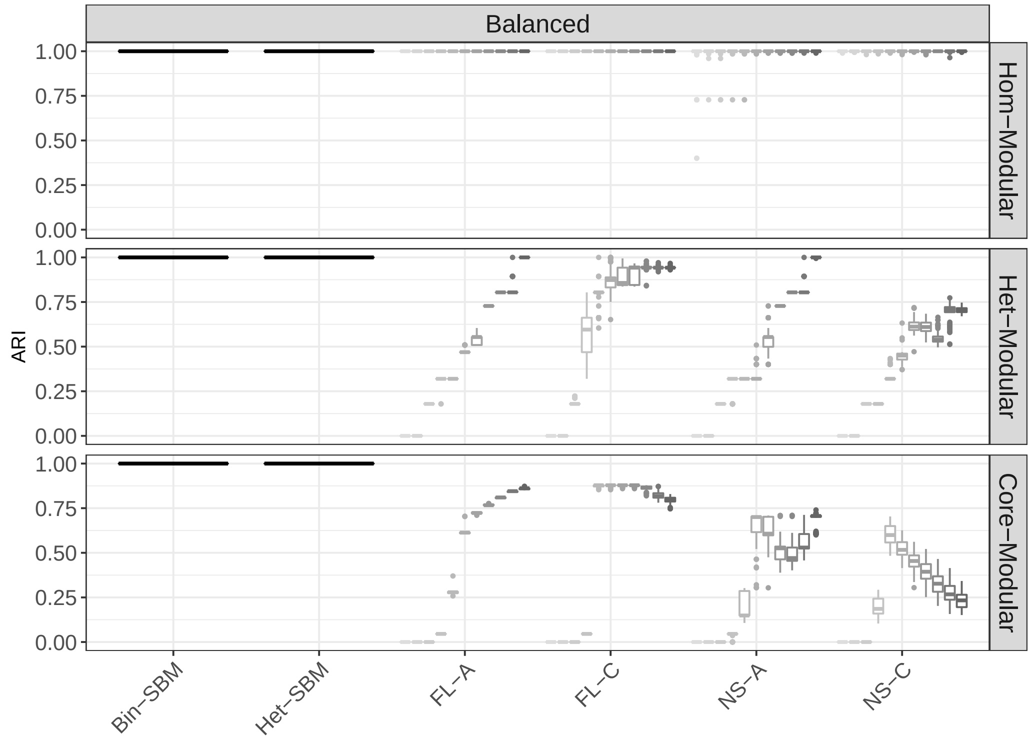


Figure S2: **Box-plots of ARI scores over 100 Monte Carlo samples in the Balanced scenarios**. The $x$-axis shows MS-SBMs and the modular algorithms, whose abbreviations FL-A and FL-C stand for the Fast Louvain algorithm with average & consensus clustering while the abbreviations NS-A and NS-C stand for the Newman Spectral algorithm with average & consensus clustering. For each modular algorithm, there are 11 box-plots whose colours correspond to the solutions with different $\gamma$ values starting from 0.5 and finishing at 1.5 with a step-size of 0.1 and the central score of 1 (default value).

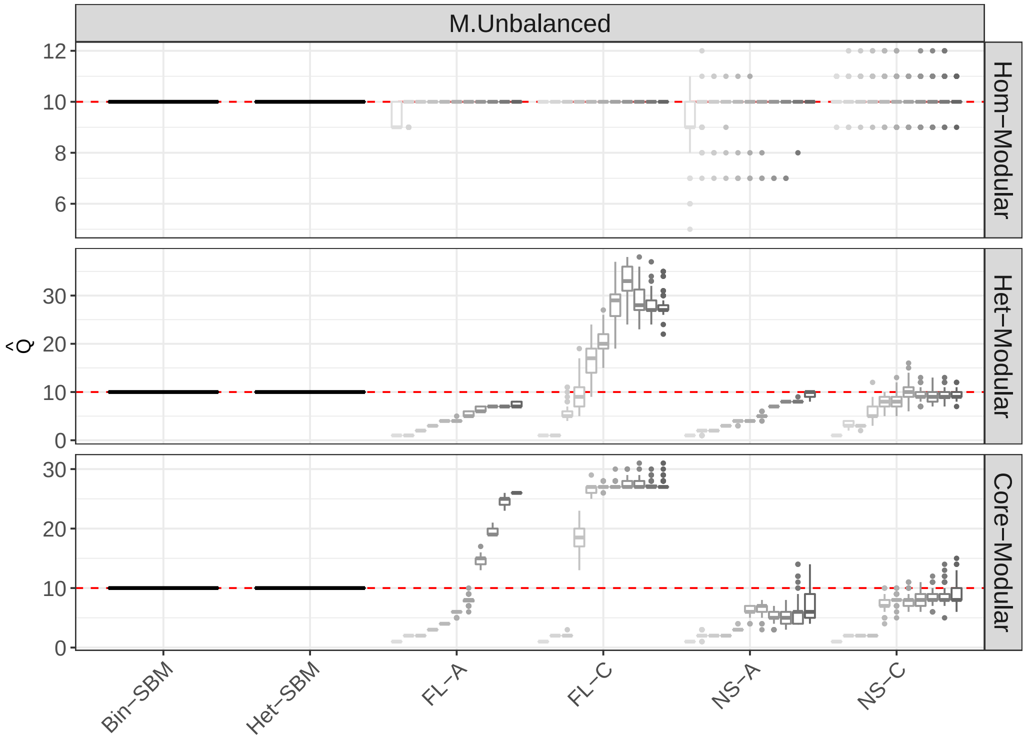


Figure S3: **Box-plots of estimated $\hat{Q}$ over 100 Monte Carlo samples in the M. Unbalanced scenarios**. The $x$-axis shows MS-SBMs and the modular algorithms, whose abbreviations FL-A and FL-C stand for the Fast Louvain algorithm with average & consensus clustering while the abbreviations NS-A and NS-C stand for the Newman Spectral algorithm with average & consensus clustering. For each modular algorithm, there are 11 box-plots whose colours correspond to the solutions with different $\gamma$ values starting from 0.5 and finishing at 1.5 with a step-size of 0.1 and the central score of 1 (default value).

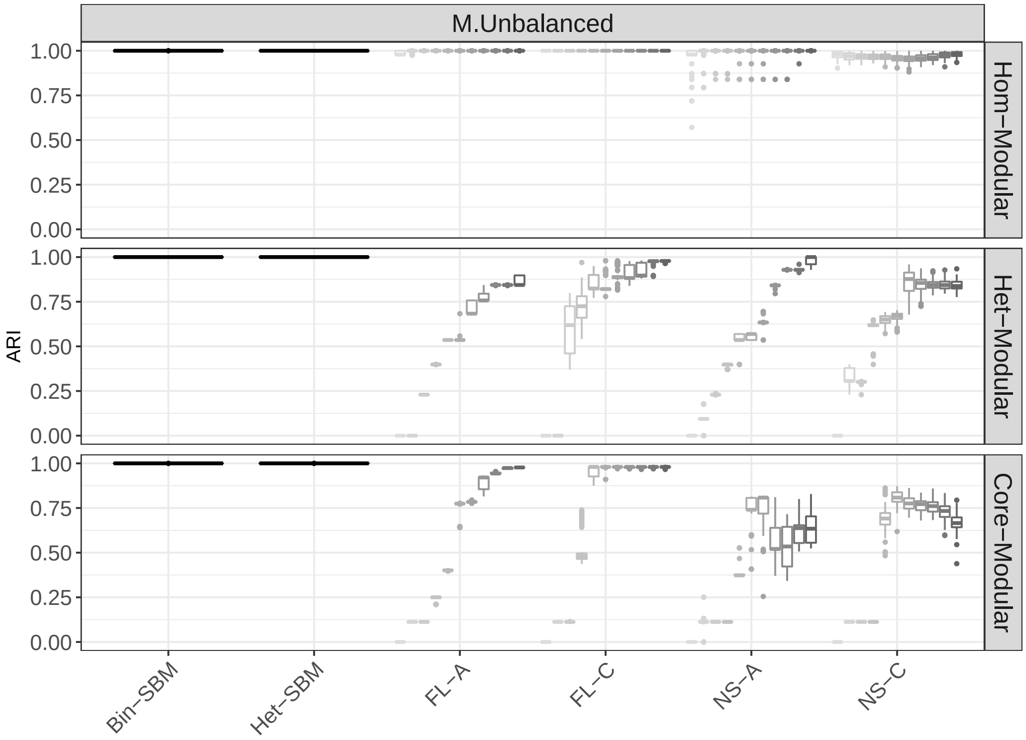


Figure S4: **Box-plots of ARI scores over 100 Monte Carlo samples in the M. Unbalanced scenarios**. The $x$-axis shows MS-SBMs and the modular algorithms, whose abbreviations FL-A and FL-C stand for the Fast Louvain algorithm with average & consensus clustering while the abbreviations NS-A and NS-C stand for the Newman Spectral algorithm with average & consensus clustering. For each modular algorithm, there are 11 box-plots whose colours correspond to the solutions with different $\gamma$ values starting from 0.5 and finishing at 1.5 with a step-size of 0.1 and the central score of 1 (default value).

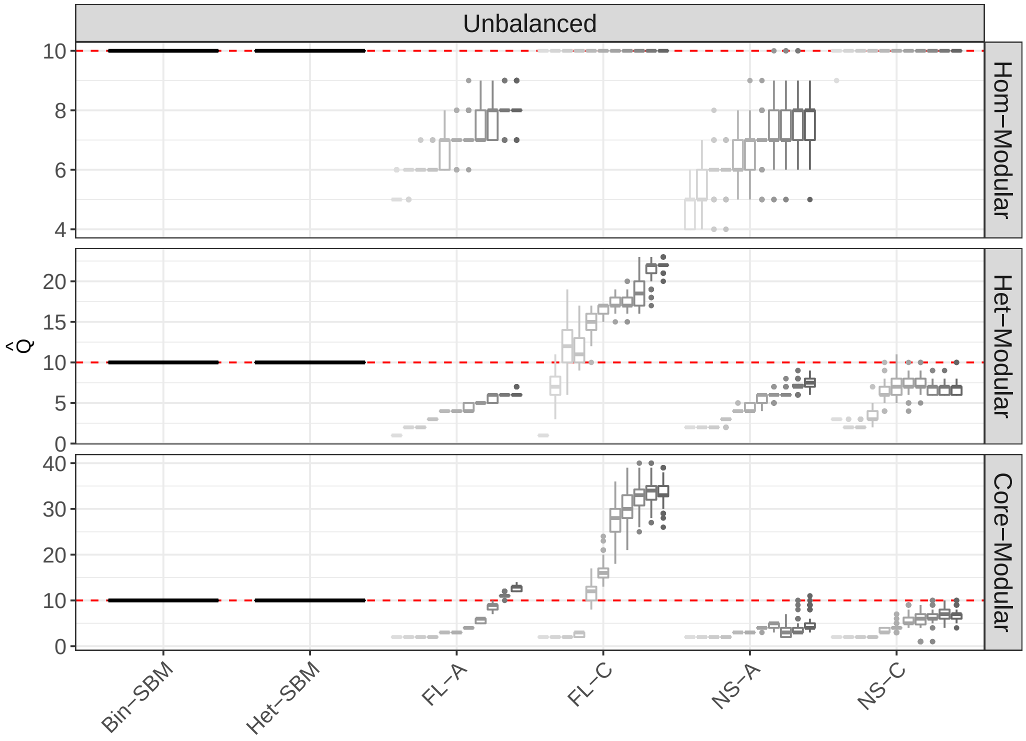


Figure S5: **Box-plots of estimated $\hat{Q}$ over 100 Monte Carlo samples in the Unbalanced scenarios**. The $x$-axis shows MS-SBMs and the modular algorithms, whose abbreviations FL-A and FL-C stand for the Fast Louvain algorithm with average & consensus clustering while the abbreviations NS-A and NS-C stand for the Newman Spectral algorithm with average & consensus clustering. For each modular algorithm, there are 11 box-plots whose colours correspond to the solutions with different $\gamma$ values starting from 0.5 and finishing at 1.5 with a step-size of 0.1 and the central score of 1 (default value).

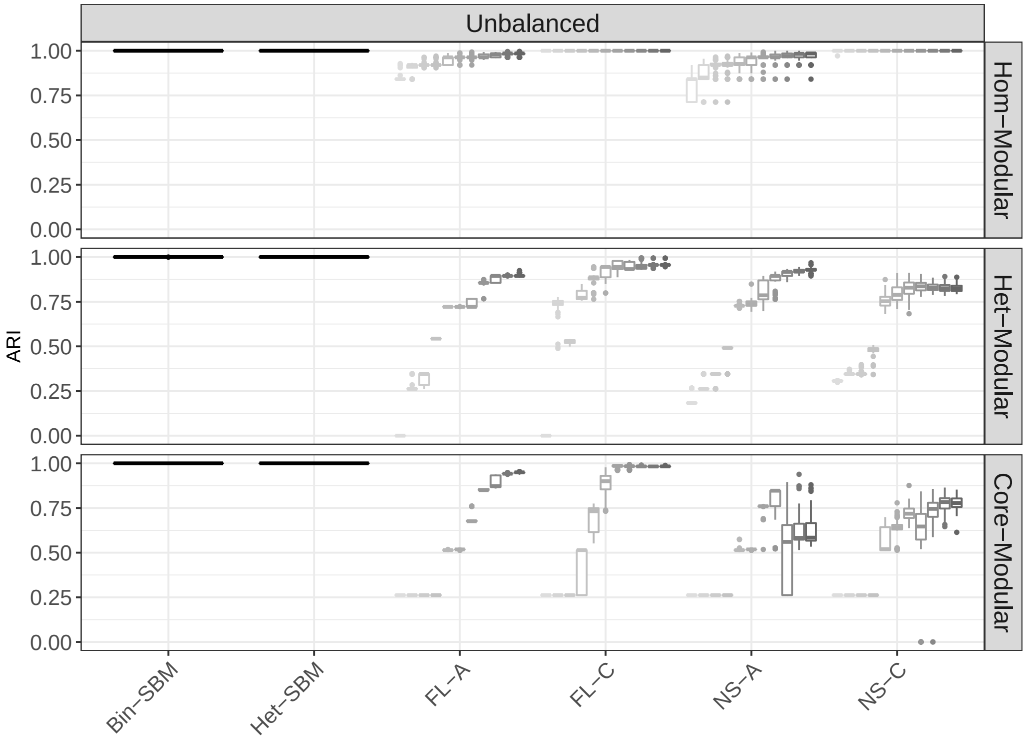


Figure S6: **Box-plots of ARI scores over 100 Monte Carlo samples in the Unbalanced scenarios**. The $x$-axis shows MS-SBMs and the modular algorithms, whose abbreviations FL-A and FL-C stand for the Fast Louvain algorithm with average & consensus clustering while the abbreviations NS-A and NS-C stand for the Newman Spectral algorithm with average & consensus clustering. For each modular algorithm, there are 11 box-plots whose colours correspond to the solutions with different $\gamma$ values starting from 0.5 and finishing at 1.5 with a step-size of 0.1 and the central score of 1 (default value).

| | Hom-Modular | | |
|---|---|---|---|
| Approach | Balanced | M. Unbalanced | Unbalanced |
| Bin-SBM | [0, 0] | [0, 0] | [0, 0] |
| Het-SBM | [0, 0] | [0, 0] | [0, 0] |
| FL-A (0.5) | [0, 0] | [-3321, 0] | [-16353, -6276] |
| FL-A (1) | [0, 0] | [0, 0] | [-6332, -1485] |
| FL-A (1.5) | [0, 0] | [0, 0] | [-3143, -585] |
| FL-C (0.5) | [0, 0] | [0, 0] | [0, 0] |
| FL-C (1) | [0, 0] | [0, 0] | [0, 0] |
| FL-C (1.5) | [0, 0] | [0, 0] | [0, 0] |
| NS-A (0.5) | [-53818, 0] | [-31440, 0] | [-28810, -6276] |
| NS-A (1) | [-77, 0] | [-11940, 0] | [-12764, -1655] |
| NS-A (1.5) | [-76, 0] | [0, 0] | [-12644, -574] |
| NS-C (0.5) | [-77, 0] | [-23366, 0] | [-4583, 0] |
| NS-C (1) | [-81, 0] | [-23549, 0] | [0, 0] |
| NS-C (1.5) | [-78, 0] | [-16837, 0] | [0, 0] |

Table S6: **Ranges of adjusted ICL score of the best fits for several approaches in the scenarios with Hom-Modular cluster structure.** For each simulated dataset and approach, the adjusted ICL score of the best fit is the difference between its ICL score and the maximum ICL score across all approaches. For each approach, the range of ICL scores is shown across all 100 Monte Carlo samples.

| Approach | Het-Modular | | |
|---|---|---|---|
| | Balanced | M. Unbalanced | Unbalanced |
| Bin-SBM | [0, 0] | [0, 0] | [0, 0] |
| Het-SBM | [0, 0] | [0, 0] | [0, 0] |
| FL-A (0.5) | [-55579, -54139] | [-68898, -67261] | [-100236, -98323] |
| FL-A (1) | [-29489, -22459] | [-28683, -21949] | [-24745, -18702] |
| FL-A (1.5) | [0, 0] | [-9987, -6090] | [-9065, -6293] |
| FL-C (0.5) | [-55579, -54139] | [-68898, -67261] | [-100236, -98323] |
| FL-C (1) | [-14485, 0] | [-10777, -1444] | [-14459, -3524] |
| FL-C (1.5) | [-2824, -647] | [-3708, -1283] | [-5737, -1421] |
| NS-A (0.5) | [-55579, -54139] | [-68898, -67261] | [-73637, -69173] |
| NS-A (1) | [-34239, -22980] | [-30941, -27768] | [-31349, -11805] |
| NS-A (1.5) | [-70, 0] | [-5464, 0] | [-9539, -3089] |
| NS-C (0.5) | [-55579, -54139] | [-68898, -67261] | [-64330, -62375] |
| NS-C (1) | [-30866, -20746] | [-32155, -19720] | [-36666, -15281] |
| NS-C (1.5) | [-29692, -22767] | [-26353, -8320] | [-36289, -21718] |

Table S7: **Ranges of adjusted ICL score of the best fits for several approaches in the scenarios with Het-Modular cluster structure.** For each simulated dataset and approach, the adjusted ICL score of the best fit is the difference between its ICL score and the maximum ICL score across all approaches. For each approach, the range of ICL scores is shown across all 100 Monte Carlo samples.

#### I. Comparisons to Block to Node SBM for Simulation I

The model described in Newman and Leicht (2007) is intended for a single and binary network $\boldsymbol{x}$ $(n \times n)$ and it can be viewed as a special case of the classical SBM of Snijders and Nowicki (1997) and Nowicki and Snijders (2001). This version imposes a Categorical density on the edges, with block-to-node parametrisation ($\boldsymbol{\pi}$ is $Q \times n$ matrix) rather than the standard block-to-block parametrisation ($\boldsymbol{\pi}$ is $Q \times Q$ matrix) and for this reason, hereafter, we refer to it as the Block to Node SBM (or BN-SBM). In BN-SBM, categories represent all the nodes in the network, thereby allowing for the modelling of heterogenous nodal profiles within a cluster.

To grasp the core philosophy of this model, in Figure S7, we provide an example of a network with 7 nodes and 3 clusters, for which we graphically illustrate its conditional likelihood $f(\boldsymbol{x}|\boldsymbol{z}; \boldsymbol{\pi})$. Although the network is undirected, the model treats it as directed, wherein each observed edge is a two-way edge (or dyad). Thus, the probability that a node $V_4$ receives an edge depends only on the cluster label of the sender nodes, which in this case is $V_5$ located in Block 2. Hence, we can write this probability as $\Pr(X_{45} = 1|Z_{52} = 1) = \pi_{25}$. This can be interpreted as the probability that a randomly selected node in Block 2 receives an edge from $V_5$; and in the opposite direction, we have $\Pr(X_{54} = 1|Z_{42} = 1) = \pi_{24}$. Furthermore, two-way-edges are assumed to be conditionally independent, such that $\Pr(X_{45} = 1, X_{54} = 1|Z_{52} = 1, Z_{42} = 1) = \pi_{25}\pi_{24}$. Due to the symmetry in the network edges, this probability can be further simplified as

$\Pr(X_{45} = 1 | Z_{52} = 1, Z_{42} = 1) = \pi_{25}\pi_{24}$. Using the notation of this work, we can write model's likelihood as

$$\begin{aligned}
\log f(\boldsymbol{x}, \boldsymbol{z}; \boldsymbol{\pi}, \boldsymbol{\alpha}) = \log f(\boldsymbol{x}|\boldsymbol{z}; \boldsymbol{\pi}) + \log f(\boldsymbol{z}; \boldsymbol{\alpha}) \\
= \sum_{i=1}^{n} \sum_{j>i}^{n} \sum_{q=1}^{Q} \sum_{l=1}^{Q} z_{iq} z_{jl} x_{ij} \log(\pi_{qi}\pi_{lj}) \\
+ \sum_{i=1}^{n} \sum_{q=1}^{Q} z_{iq} \log \alpha_q.
\end{aligned}$$

The estimation is performed with the EM algorithm as already discussed

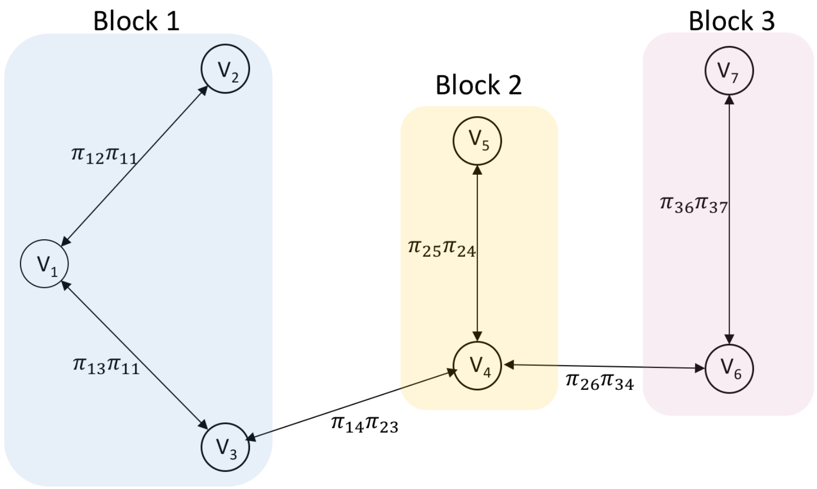


Figure S7: **BN-SBM, block-to-node parametrisation and a graphical illustration of the conditional likelihood** $\log f(\boldsymbol{x}|\boldsymbol{z}; \boldsymbol{\pi})$. The network with 7 nodes and 3 clusters is binary and undirected. Although the network is undirected, the edges are treated as two-way links. Each link is annotated with two parameters accounting for each direction.

in Snijders and Nowicki (1997), and fixed-point estimating equations can be found in Newman and Leicht (2007). Therein, the SI material provides a `C` based implementation from which we use to benchmark this model.

Similar to the work of Snijders and Nowicki (1997), the work of Newman and Leicht (2007) does not propose any new strategies to estimate the optimal number of clusters $Q$, and authors simply suggest a solution which maximises the likelihood. Although this is a reasonable procedure for selecting an optimal fit from a collection of different initialisations with fixed $Q$, it is not appropriate for comparisons between the fits with different $Q$. Indeed, the likelihood function is monotonically increasing with increasing $Q$ and, therefore, solutions with higher number of clusters will always be preferred over those with smaller number of clusters. For this reason, we could not test this model on its ability to estimate optimal $Q$ like we did for the other MS SBMs. Hence, we only considered the case in which the ground truth $Q$ is supplied to the model.

BN-SBM is adapted for a single and binary network and as such it cannot be fitted for all 30 subjects jointly. To compensate for this, we consider two implementations (labelled hereafter as M1 and M2). In M1, we fit the model only on the first subject and, thus, we assume that there is only one realisation per each Monte Carlo dataset. This seems reasonable as the cluster labels remain fixed across the subjects in Simulation I. In M2, we average data across 30 subjects and, threshold this average to find one representative, binary network for all subjects ($x_{ij} \geq 0.5 = 1$ and 0 otherwise). The latter version circumvents the small sample effects of the former approach. In both versions, we use 50 initialisations for each Monte Carlo realisation, and as suggested by Newman and Leicht (2007) we use the likelihood scores to select the winning model among 50 realisations.

Figure S8 shows that the model is accurate for the Hom-Modular case across both implementations (M1 and M2). In the Het-Modular case, however, there is a better performance in M2 over M1, which can be explained by the averaging and thresholding of datasets which tended to convert Heterogeneous-Modular structure to Homogeneous-Modular structure. This effect can be seen in Figure S9 in which we show one such representative matrix for the Het-Modular case and Balanced design. Thus, the results in M1 are a more accurate reflection of the model's ability to estimate Het-Modular structures. It is worth noting that BN-SBM struggles with Core-Modular cluster structure, and the `C` code found in SI Newman and Leicht (2007) failed to produce numerical outputs (with a total of 2,090 instances in M1 and 15,000 instances, the entire scope of Simulation I, in M2). There is some improvement in ARI scores across three block designs, but this upward trend can be easily explained by the smaller core block sizes and larger modular block sizes.

These results can be explained by the constraint of Categorical density on its parameters, i.e. $\sum_{i=1}^{n}\sum_{j=1}^{n}\pi_{qi}\pi_{lj} = 1$. This enforces a restriction on the types of cluster structures that BN-SBM can estimate. For example, consider a cluster structure with two blocks in which an edge probability is 0.98 in Block 1 (i.e. $\pi_{11} = 0.98$) and 0.7 in Block (1, 2) (i.e. $\pi_{12} = 0.7$). Due to the constraint that each block-to-node probability must add up to 1, it is evident that the model is unable to capture the standard catalog of cluster structures. This limitation is particularly pronounced in the Core-

Modular case. Indeed, Newman and Leicht (2007) advertise this model only for the modular and disassortative structures. Of the modular types, only the Homogeneous-Modular cases were well captured by our simulations. Based on this, it appears that the modelling of nodal-profiles comes at the heavy cost of the model's richness and ability to estimate different cluster structures, a trade-off that we have found to be unsatisfactory.

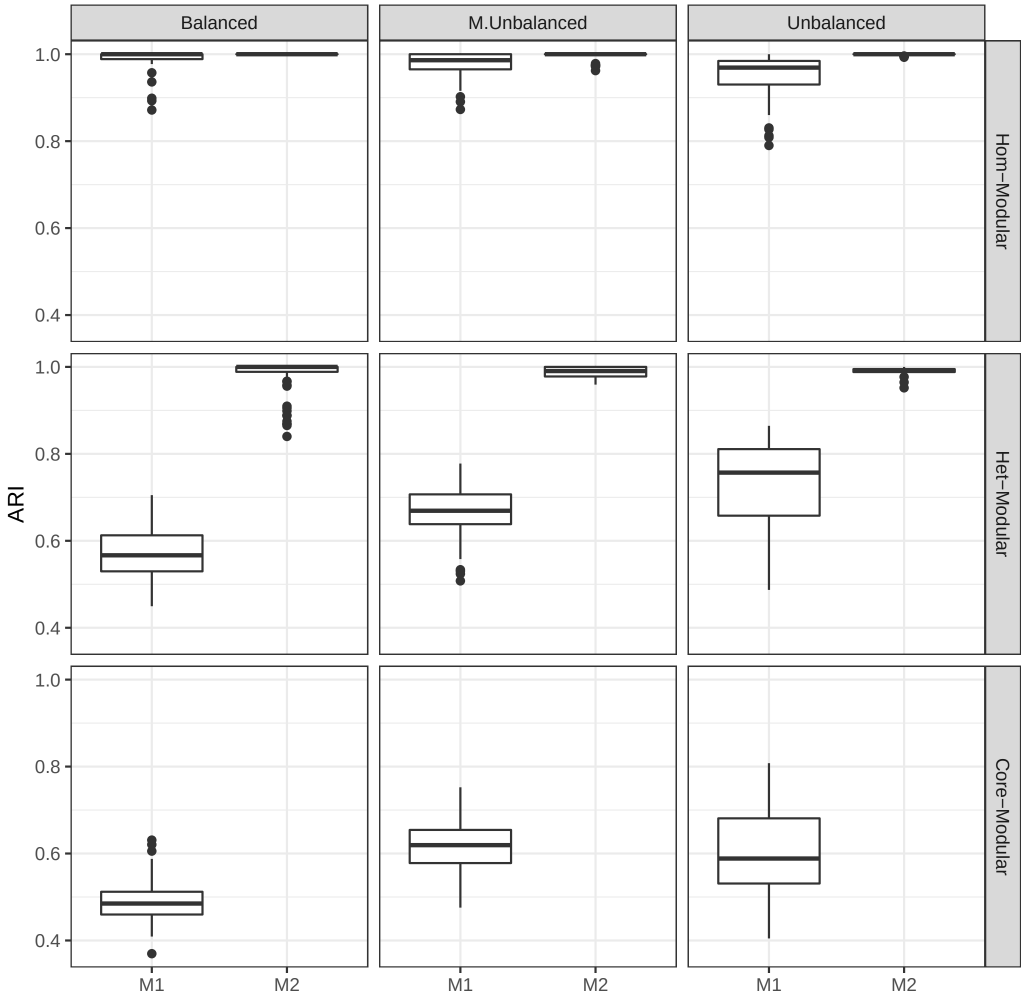


Figure S8: **Simulation I results for BN-SBM results**. The ground truth total number of clusters was supplied to the model. The x-axis shows two versions of the BN-SBM implementation (Newman and Leicht, 2007), labelled as M1 and M2. In M1, the first subject's dataset is used to estimate the cluster labels, while, in M2, each Monte Carlo dataset was averaged across 30 subjects and then thresholded ($x_{ij} \geq 0.5 = 1$ and 0 otherwise) to obtain a single, binary network. The y-axis represents ARI scores between estimated and true cluster labels. Each cell in the grid represents results in terms of three block designs (Balanced, M. Unbalanced and Unbalanced) and three cluster structures (Hom-Modular, Het-Modular and Core-Modular). The `C` code found in SI of Newman and Leicht (2007) failed to produce numerical outputs in the cases of Core-Modular cluster structure, with a total of 2,090 such instances in M1, and 15,000 in M2 (the entire scope of Simulation I).

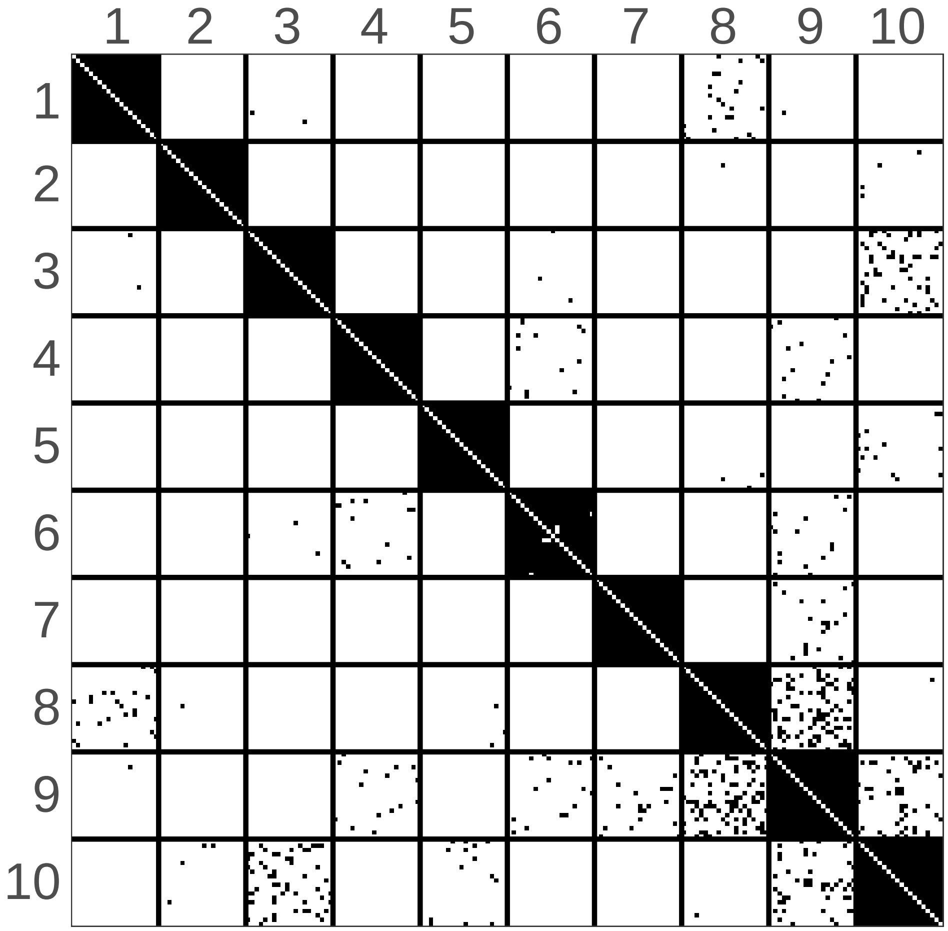


Figure S9: **Example of binarised network representation of one Monte Carlo multi-subject realisation**. This dataset was sampled from the Het-Modular structure, but as noted the averaging and binarisation tend to make this structure more homogeneous.

#### J. Initialisation Strategy Comparison

Since the goal of this exercise is to gauge the improvements of MS-SBMs over initialisations, we have only considered the slice of Simulation I where $Q$ was set to 10 (i.e. $\hat{Q} = 10$). Apart from this, the entire setup of Simulation I has been kept the same as in Section 3.1. In Figures S10 - S12, we compare the ARI scores of Bin-SBM fits, Het-SBM fits and the initialisations to the ground truth. We can observe a trend in which the degree of improvement from the starting point is moderated by proximity of initialisation to the local maximum. Specifically, in scenarios where initialisation is close to the local maximum (or already the local maximum), the algorithm will quickly converge to it and there will be not much improvement over the starting point. This finding is supported by a well known property of the variational EM algorithm (see Proposition 7 in Daudin et al. (2008)), which guarantees a monotonic increase of the variational bound and convergence to a local maximum. This property is also consistent with the classical EM algorithm which does not guarantee convergence to a global maximum, but only to a local maximum. To overcome the issue of convergence to a local maximum, the models rely on the ICL scores to select the global maximum. As shown in Figures S13 - S16, the ICL scores are the highest for the ARI scores of 1, suggesting that the ICL criterion can be taken as a reliable measure when fitting the models without knowledge of the grand truth.

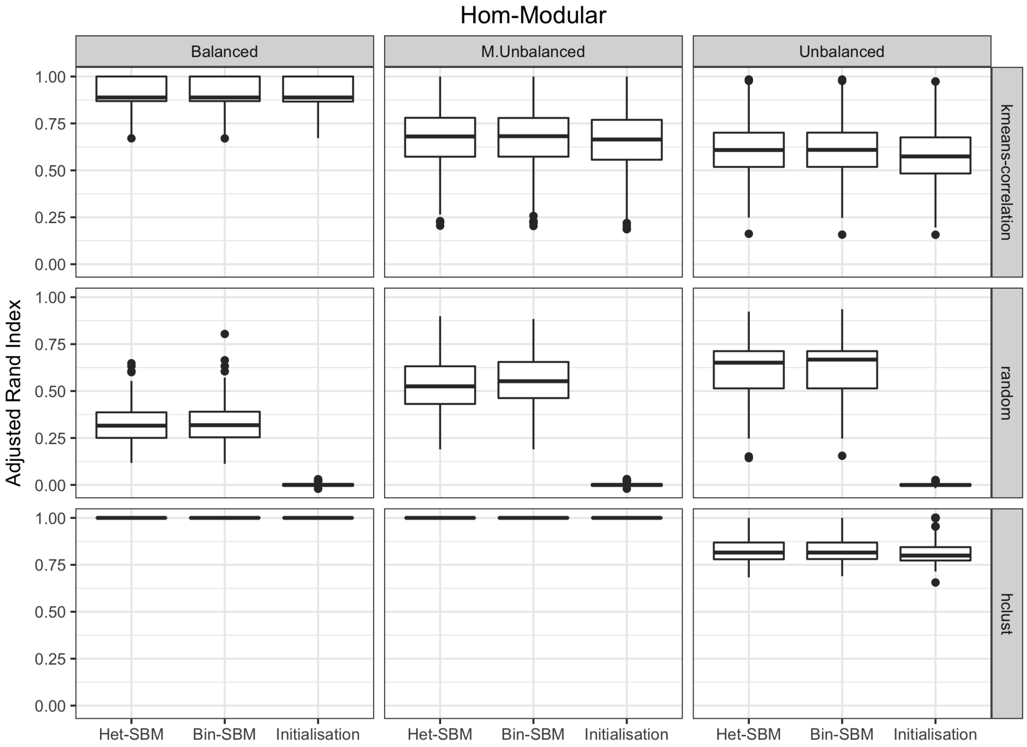


Figure S10: **Comparing ARI scores of the Bin-SBM fits, the Het-SBM fits and the initialisations to the ground truth for the Hom-Modular case**. Vertical panels show three different block designs, and horizontal panels show three different initialisation strategies. Overall, modest improvements have been achieved for the initialisations utilising k-means and hclust but strong improvements have been detected for the fits utilising random sampling.

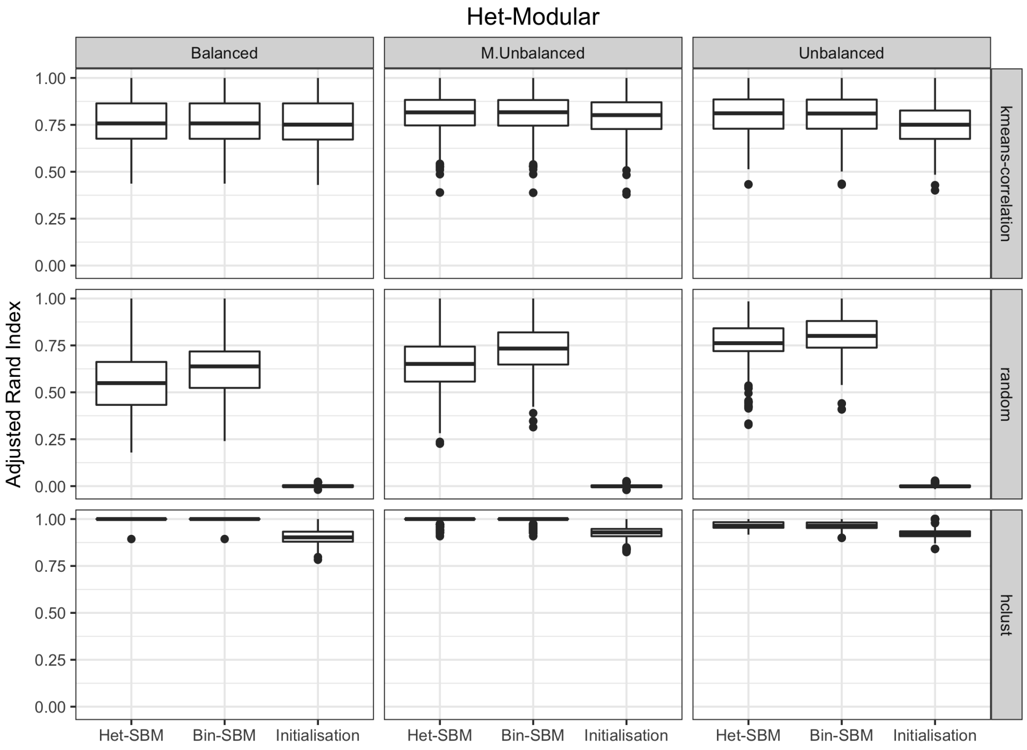


Figure S11: **Comparing ARI scores of the Bin-SBM fits, the Het-SBM fits and the initialisations to the ground truth for the Het-Modular case**. Vertical panels show three different block designs, and horizontal panels show three different initialisation strategies. Overall, modest improvements have been achieved for the initialisations with k-means and hclust but strong improvements are detected for the random sampling.

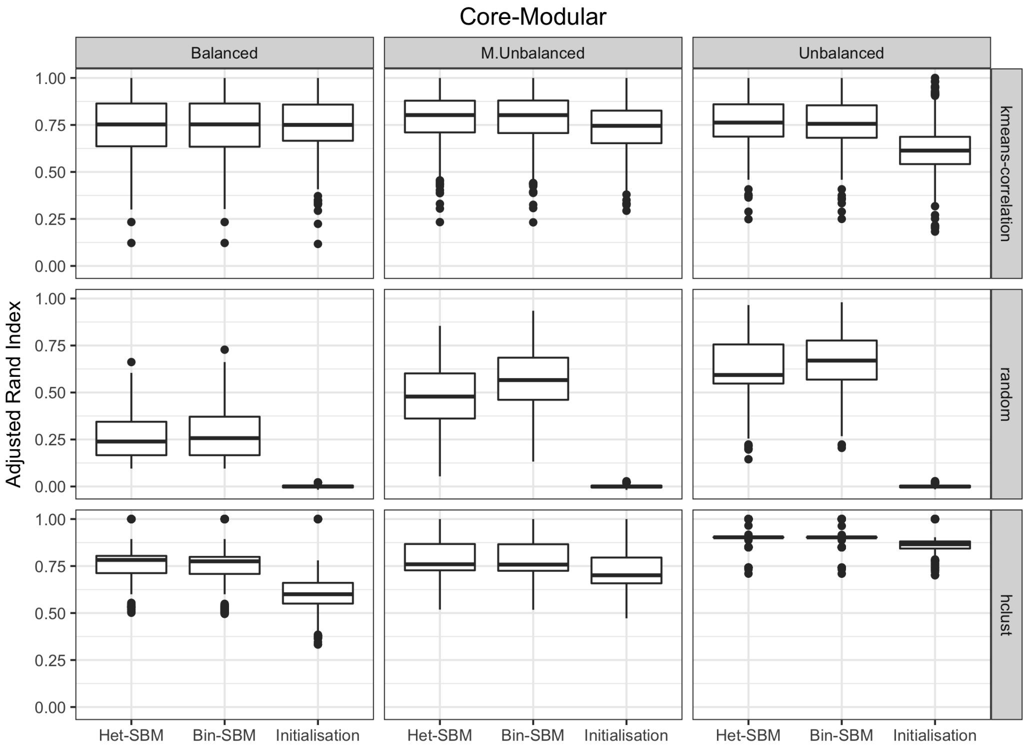


Figure S12: **Comparing ARI scores of the Bin-SBM fits, the Het-SBM fits and the initialisations to the ground truth for the Core-Modular case**. Vertical panels show three different block design, and horizontal panels show three different initialisation strategies. Overall, modest improvements have been achieved for the initialisations with k-means and hclust but strong improvements are detected for the random sampling.

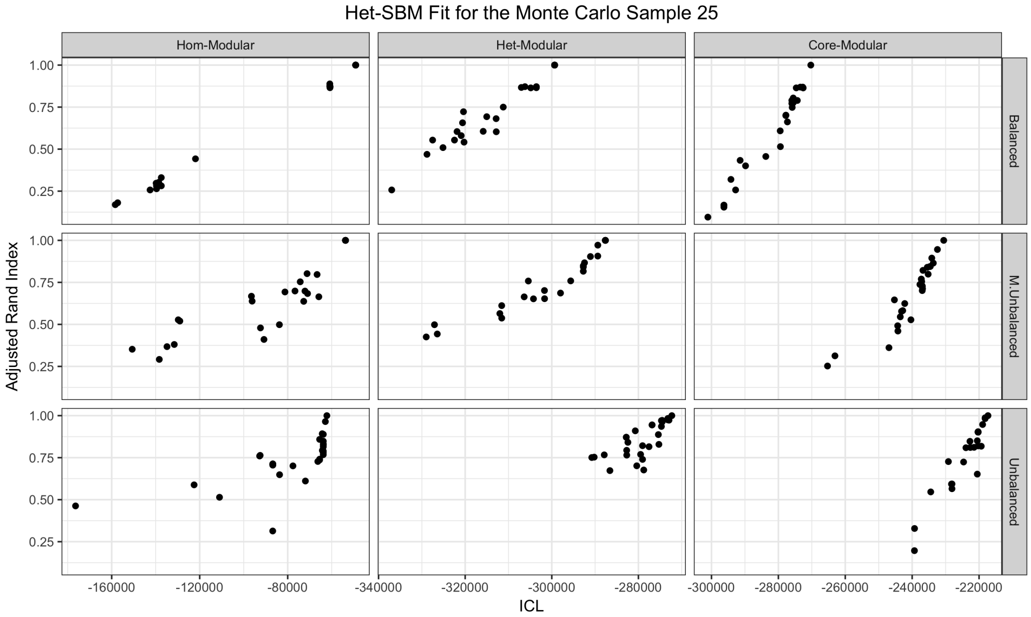


Figure S13: **ARI scores as a function of the ICL scores of the Het-SBM fits for the Monte Carlo Sample 25 across all initialisation strategies**. Vertical panels show three different cluster structures, and horizontal panels show three different block designs. Overall, there is a strong linear association between the ARI and ICL scores, so that ARI scores increase with increasing ICL scores, demonstrating the ability of the ICL criterion to select the best fits. In particular, the best ICL score always correspond to an ARI score of 1 (i.e. the ground truth).

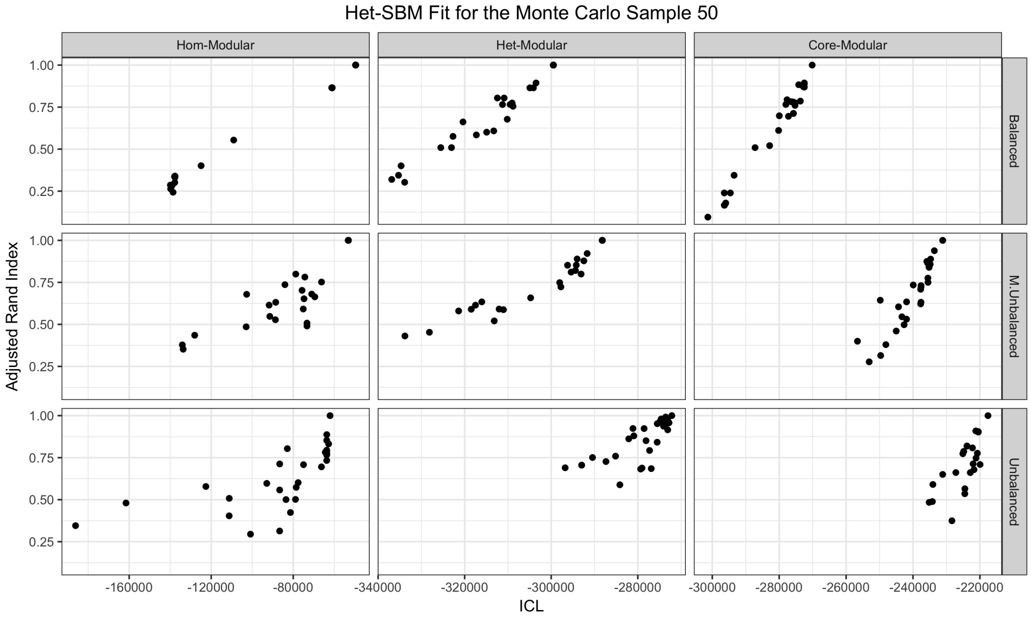


Figure S14: **ARI scores as a function of the ICL scores of the Het-SBM fits for the Monte Carlo Sample 50 across all initialisation strategies**. Vertical panels show three different cluster structures, and horizontal panels show three different block designs. Overall, there is a strong linear association between the ARI and ICL scores, so that ARI scores increase with increasing ICL scores, demonstrating the ability of the ICL criterion to select the best fits. In particular, the best ICL score always correspond to an ARI score of 1 (i.e. the ground truth).

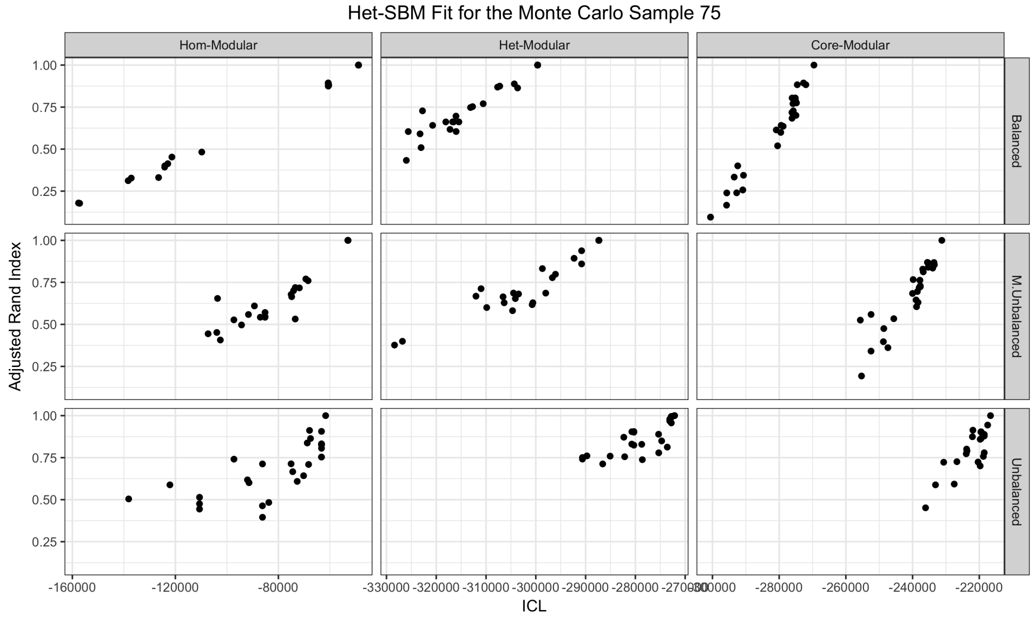


Figure S15: **ARI scores as a function of the ICL scores of the Het-SBM fits for the Monte Carlo Sample 75 across all initialisation strategies**. Vertical panels show three different cluster structures, and horizontal panels show three different block designs. Overall, there is a strong linear association between the ARI and ICL scores, so that ARI scores increase with increasing ICL scores, demonstrating the ability of the ICL criterion to select the best fits. In particular, the best ICL score always correspond to an ARI score of 1 (i.e. the ground truth).

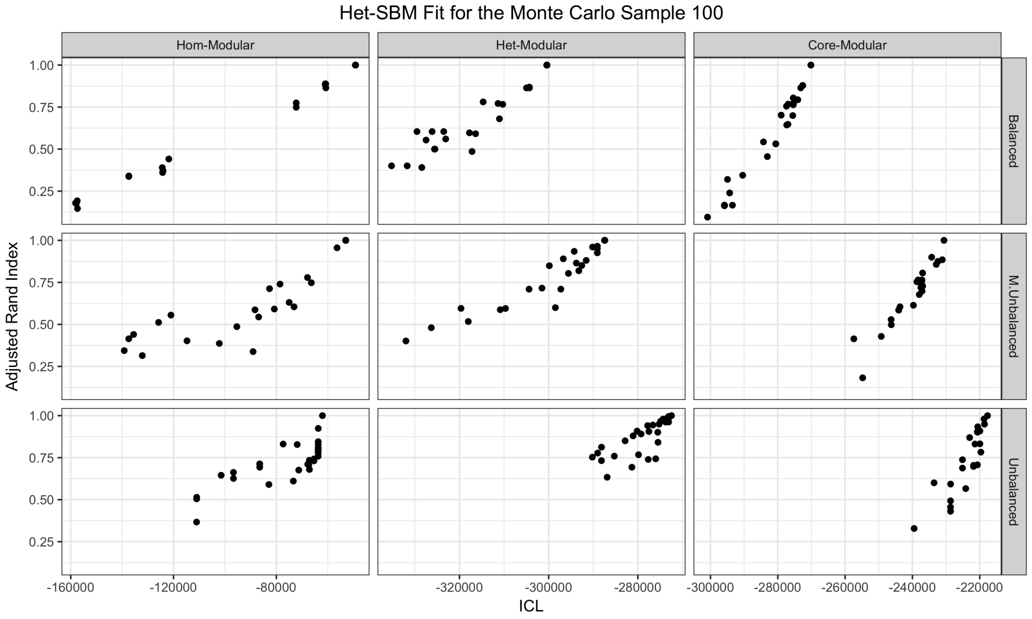


Figure S16: **ARI scores as a function of the ICL scores of the Het-SBM fits for the Monte Carlo Sample 100 across all initialisation strategies**. Vertical panels show three different cluster structures, and horizontal panels show three different block designs. Overall, there is a strong linear association between the ARI and ICL scores, so that ARI scores increase with increasing ICL scores, demonstrating the ability of the ICL criterion to select the best fits. In particular, the best ICL score always correspond to an ARI score of 1 (i.e. the ground truth).

#### K. Additional Results for Simulation II

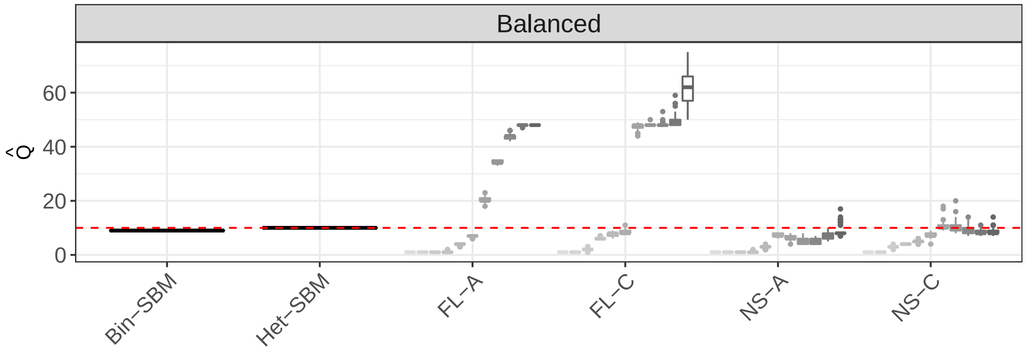


Figure S17: **Box-plots of total number of cluster $\hat{Q}$ estimates over 100 Monte Carlo samples in the Balanced design**. The $x$-axis shows MS-SBMs and the modular algorithms, whose abbreviations FL-A and FL-C stand for the Fast Louvain algorithm with average & consensus clustering while the abbreviations NS-A and NS-C stand for the Newman Spectral algorithm with average & consensus clustering. For each modular algorithm, there are 11 box-plots whose colours correspond to the solutions with different $\gamma$ values starting from 0.5 and finishing at 1.5 with a step-size of 0.1 and the central score of 1 (default value).

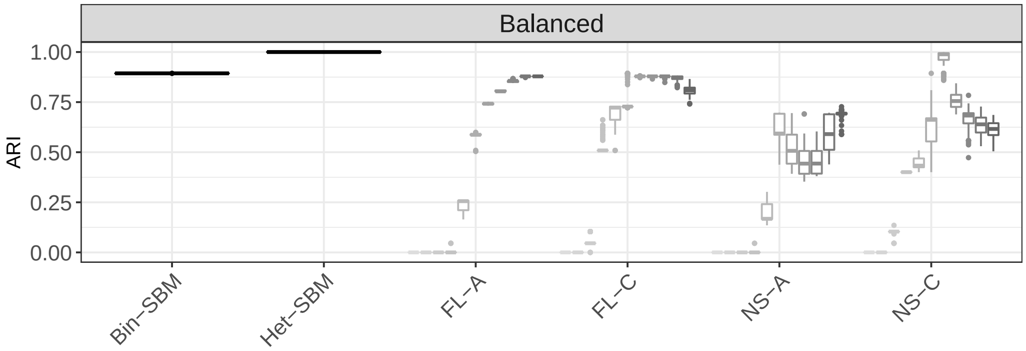


Figure S18: **Box-plots of ARI scores over 100 Monte Carlo samples in the Balanced design**. The $x$-axis shows MS-SBMs and the modular algorithms, whose abbreviations FL-A and FL-C stand for the Fast Louvain algorithm with average & consensus clustering while the abbreviations NS-A and NS-C stand for the Newman Spectral algorithm with average & consensus clustering. For each modular algorithm, there are 11 box-plots whose colours correspond to the solutions with different $\gamma$ values starting from 0.5 and finishing at 1.5 with a step-size of 0.1 and the central score of 1 (default value).

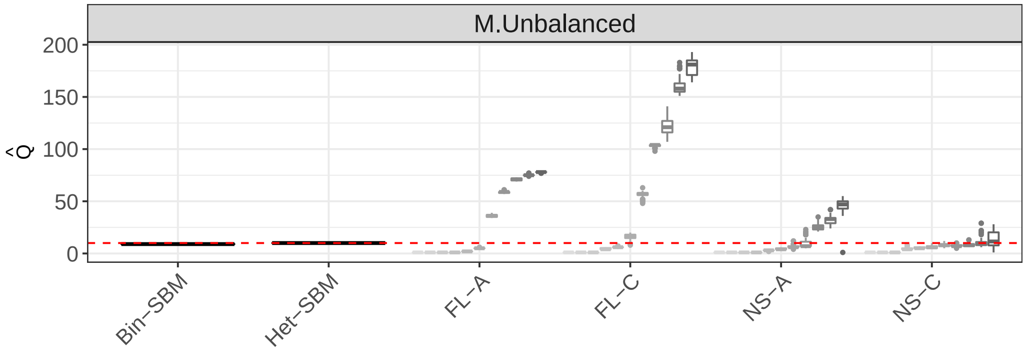


Figure S19: **Box-plots of total number of cluster $\hat{Q}$ estimates over 100 Monte Carlo samples in the M. Unbalanced design**. The $x$-axis shows MS-SBMs and the modular algorithms, whose abbreviations FL-A and FL-C stand for the Fast Louvain algorithm with average & consensus clustering while the abbreviations NS-A and NS-C stand for the Newman Spectral algorithm with average & consensus clustering. For each modular algorithm, there are 11 box-plots whose colours correspond to the solutions with different $\gamma$ values starting from 0.5 and finishing at 1.5 with a step-size of 0.1 and the central score of 1 (default value).

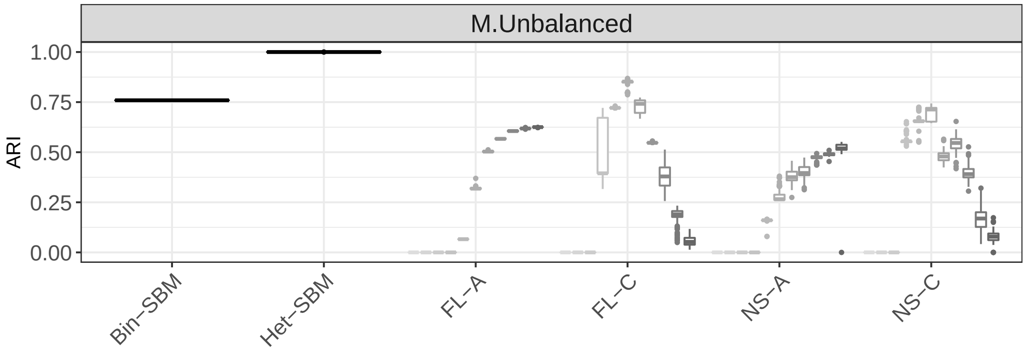


Figure S20: **Box-plots of ARI scores over 100 Monte Carlo samples in the M. Unbalanced design**. The $x$-axis shows MS-SBMs and the modular algorithms, whose abbreviations FL-A and FL-C stand for the Fast Louvain algorithm with average & consensus clustering while the abbreviations NS-A and NS-C stand for the Newman Spectral algorithm with average & consensus clustering. For each modular algorithm, there are 11 box-plots whose colours correspond to the solutions with different $\gamma$ values starting from 0.5 and finishing at 1.5 with a step-size of 0.1 and the central score of 1 (default value).

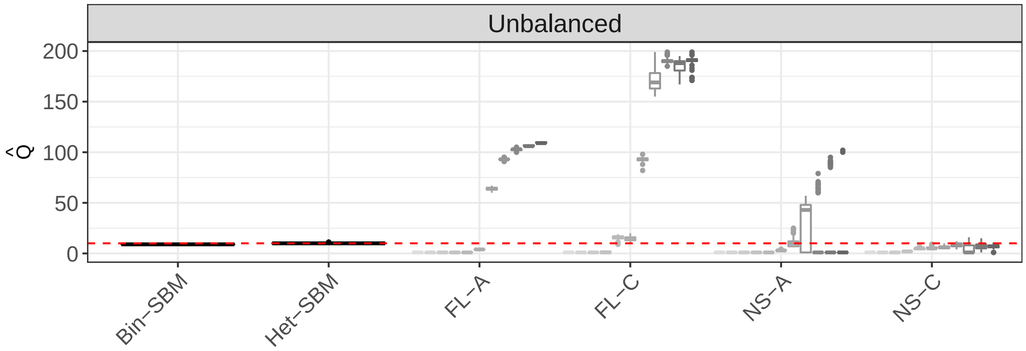


Figure S21: **Box-plots of total number of cluster $\hat{Q}$ estimates over 100 Monte Carlo samples in the Unbalanced design**. The $x$-axis shows MS-SBMs and the modular algorithms, whose abbreviations FL-A and FL-C stand for the Fast Louvain algorithm with average & consensus clustering while the abbreviations NS-A and NS-C stand for the Newman Spectral algorithm with average & consensus clustering. For each modular algorithm, there are 11 box-plots whose colours correspond to the solutions with different $\gamma$ values starting from 0.5 and finishing at 1.5 with a step-size of 0.1 and the central score of 1 (default value).

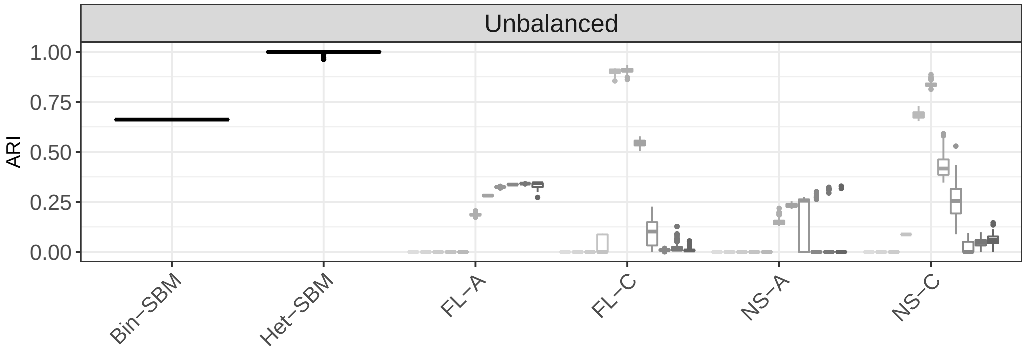


Figure S22: **Box-plots of ARI scores over 100 Monte Carlo samples in the Unbalanced design**. The $x$-axis shows MS-SBMs and the modular algorithms, whose abbreviations FL-A and FL-C stand for the Fast Louvain algorithm with average & consensus clustering while the abbreviations NS-A and NS-C stand for the Newman Spectral algorithm with average & consensus clustering. For each modular algorithm, there are 11 box-plots whose colours correspond to the solutions with different $\gamma$ values starting from 0.5 and finishing at 1.5 with a step-size of 0.1 and the central score of 1 (default value).

#### L. Additional Results for Simulations III

In this section, we provide some additional information related to the Simulation III. In Table S8, we summarize block sizes in terms of three different block designs and network with 30,60 and 120 nodes.

| Network Sizes | Block Design | Block Sizes | | |
|---|---|---|---|---|
| $n = 30$ | Balanced | 10 | 10 | 10 |
| | M. Unbalanced | 18 | 19 | 3 |
| | Unbalanced | 21 | 6 | 3 |
| $n = 60$ | Balanced | 20 | 20 | 20 |
| | M. Unbalanced | 36 | 18 | 6 |
| | Unbalanced | 42 | 12 | 6 |
| $n = 120$ | Balanced | 40 | 40 | 40 |
| | M. Unbalanced | 72 | 36 | 12 |
| | Unbalanced | 84 | 24 | 12 |

Table S8: **Proportion Designs**. Number of nodes for each scenario. The Balanced, Mildly Unbalanced (M. Unbalanced) and Unbalanced proportion designs are defined as the ratio of individual block sizes and the total number of nodes in a network.

In the remainder of this section, we provide several plots related to the control of the FPR that were not shown in the main paper. In the case of no random effect, the overall conclusion is that the parametric tests tend to be more accurate as the sample sizes increase (i.e. more subjects and nodes). Nevertheless, it is interesting to point out that some conservative behaviour is still present in the data for the Wald test especially for the data simulated according to $\boldsymbol{\pi}_1$. Thus, in order to get accurate inferences for this case, the blocks need to contain at least 12 nodes and the total number of subjects should be at least 20 (see Figure S28). However, the permutation tests seem to be more accurate even in the small sample settings and, for this reason, it is more recommended than the parametric procedures. The permutation tests are especially valuable in real data applications because there we really do not have any control over the sizes of blocks since the Het-SBM is free to fit any block size. Thus, even if the assumptions of the Het-SBM model are met, it is still possible to encounter some inaccuracies with the parametric tests. In addition to this, we also note that in the case of random effects (or imposed dependencies between edges within subject specific blocks), the permutation tests are still accurate, while the parametric tests show wildly inflated FPRs and are therefore invalid.

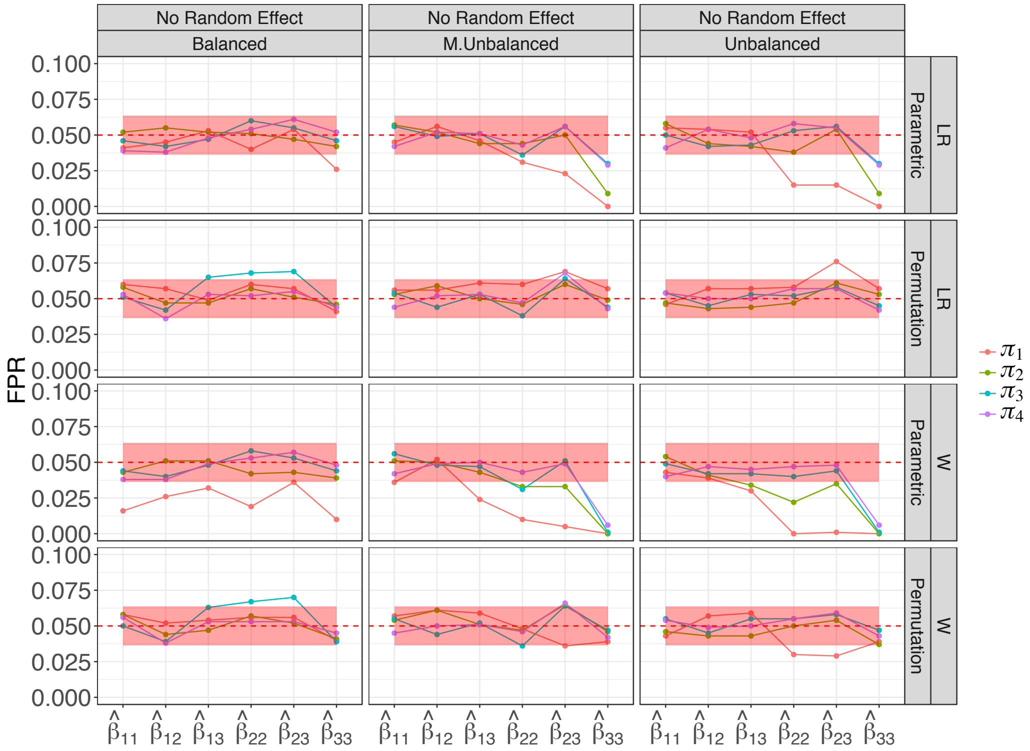


Figure S23: **Observed FPR for the simulated networks with 30 nodes, 10 subjects and no random effect.** The $x$-axis represents the block specific fitted regression coefficients. The observed FPR are colour coded according to 4 different connectivity matrices. The columns represent 3 different block designs and the rows denote the likelihood ratio (LR) and Wald (W) scores based on the parametric and permutation procedures. The red shaded strip represents the 95% Monte Carlo binomial proportion confidence intervals. While all tests appear to be valid, there is some conservative behaviour in the Unbalanced design for the Wald ($\boldsymbol{\pi}_1$-$\boldsymbol{\pi}_4$) and LR test ($\boldsymbol{\pi}_1$-$\boldsymbol{\pi}_2$). Overall permutation tests tend to be more accurate than parametric tests.

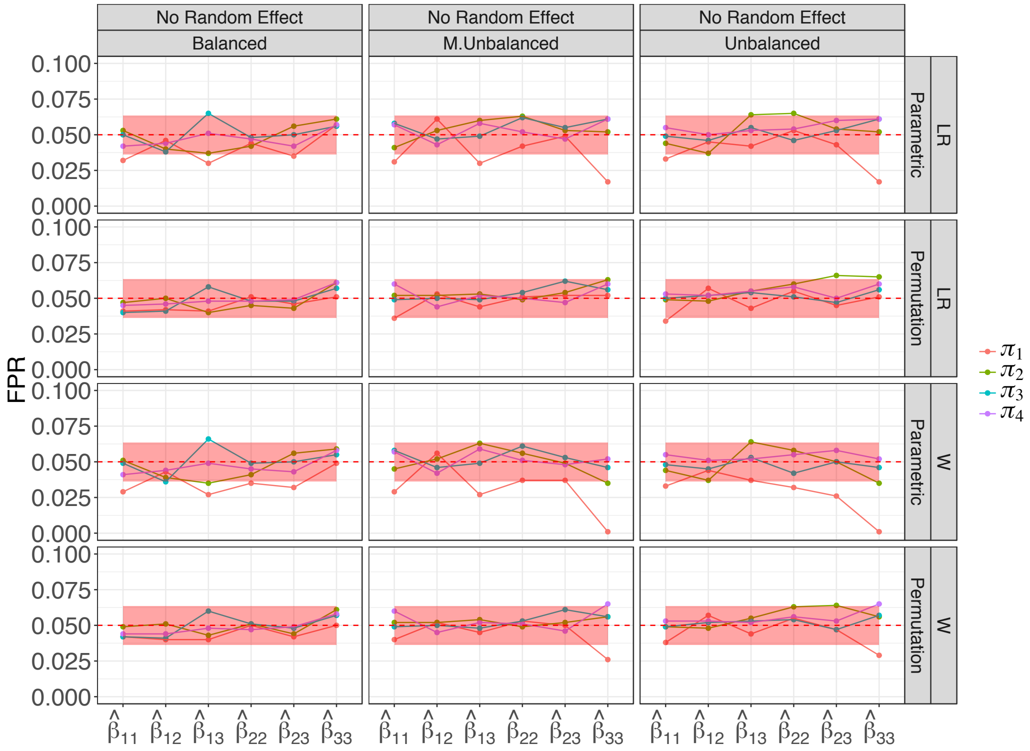


Figure S24: **Observed FPR for the simulated networks with 60 nodes, 10 subjects and no random effect.** The $x$-axis represents the block specific fitted regression coefficients. The observed FPRs are colour coded according to 4 different connectivity matrices. The columns represent 3 different block designs and the rows denote the likelihood ratio (LR) and Wald (W) scores based on the parametric and permutation tests. The red shaded strip represents the 95% Monte Carlo binomial proportion confidence intervals.

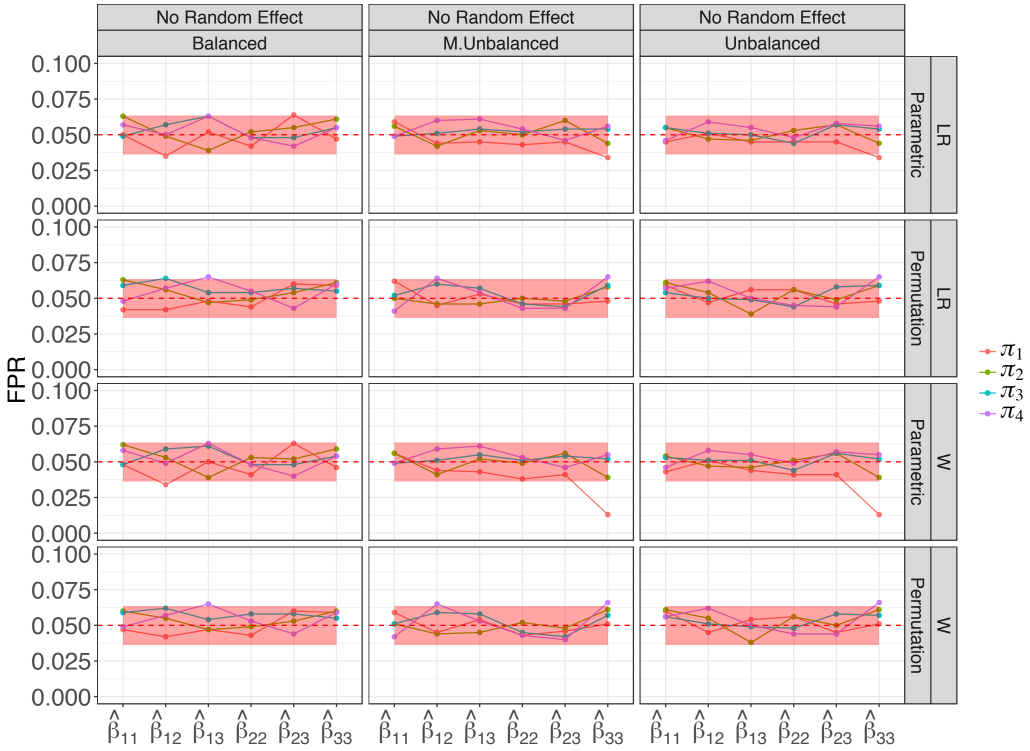


Figure S25: **Observed FPR for the simulated networks with 120 nodes, 10 subjects and no random effect.** The $x$-axis represents the block specific fitted regression coefficients. The observed FPRs are colour coded according to 4 different connectivity matrices. The columns represent 3 different block designs and the rows denote the likelihood ratio (LR) and Wald (W) scores based on the parametric and permutation tests. The red shaded strip represents the 95% Monte Carlo binomial proportion confidence intervals.

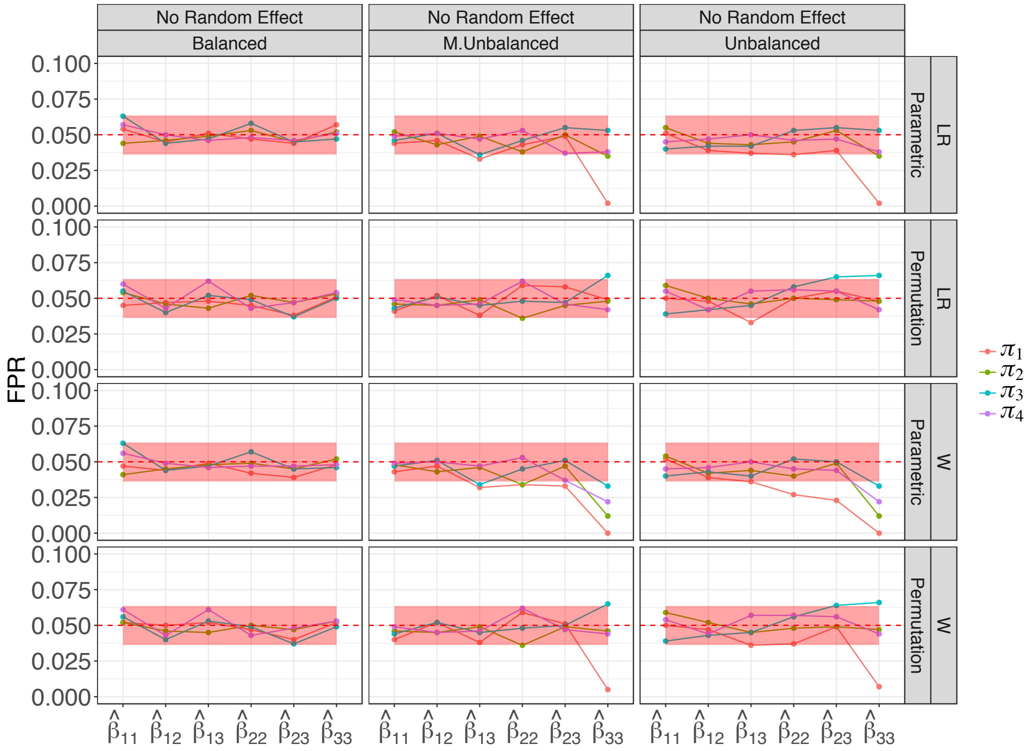


Figure S26: **Observed FPR for the simulated networks with 30 nodes, 20 subjects and no random effect.** The $x$-axis represents the block specific fitted regression coefficients. The observed FPRs are colour coded according to 4 different connectivity matrices. The columns represent 3 different block designs and the rows denote the likelihood ratio (LR) and Wald (W) scores based on the parametric and permutation tests. The red shaded strip represents the 95% Monte Carlo binomial proportion confidence intervals.

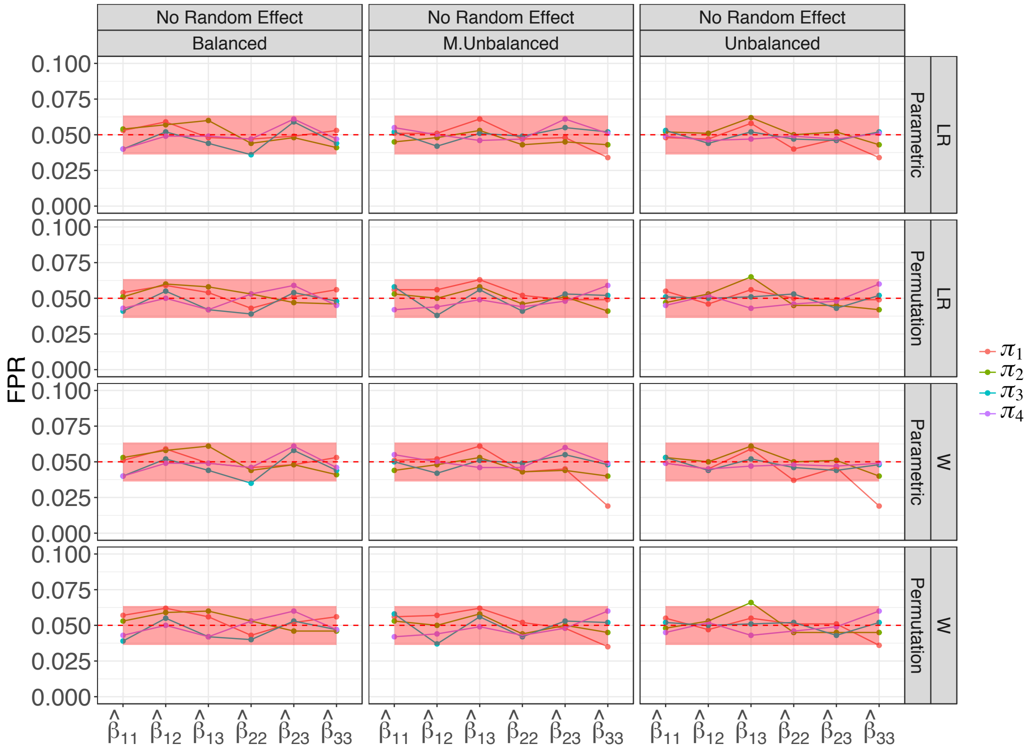


Figure S27: **Observed FPR for the simulated networks with 60 nodes, 20 subjects and no random effect.** The $x$-axis represents the block specific fitted regression coefficients. The observed FPRs are colour coded according to 4 different connectivity matrices. The columns represent 3 different block designs and the rows denote the likelihood ratio (LR) and Wald (W) scores based on the parametric and permutation tests. The red shaded strip represents the 95% Monte Carlo binomial proportion confidence intervals.

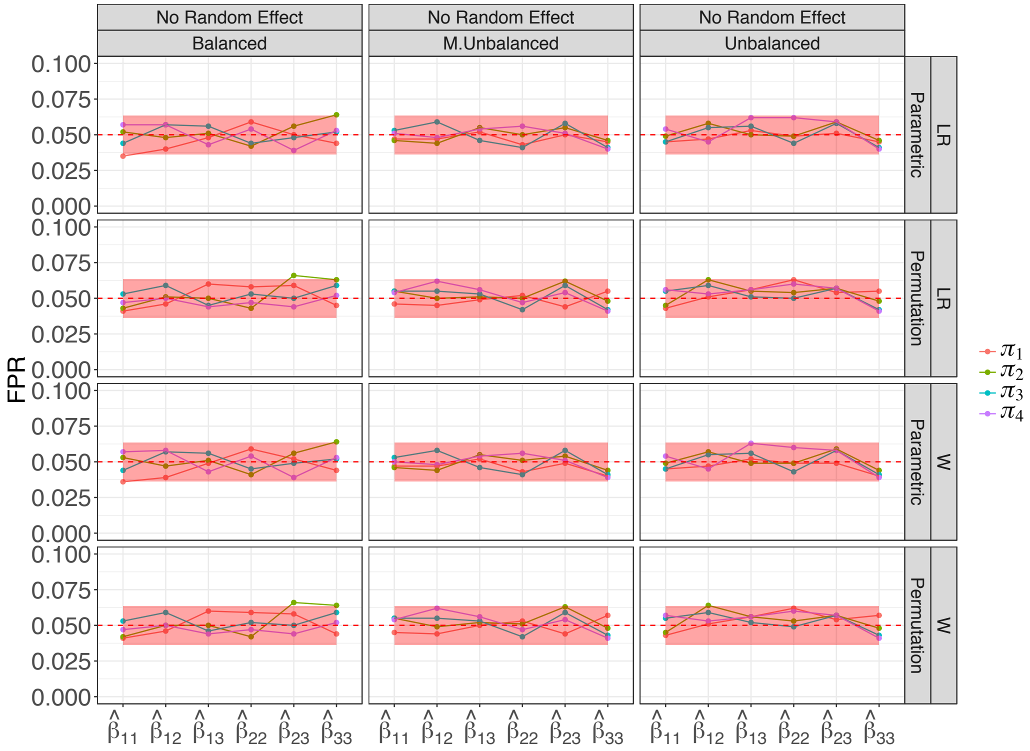


Figure S28: **Observed FPR for the simulated networks with 120 nodes, 20 subjects and no random effect.** The $x$-axis represents the block specific fitted regression coefficients. The observed FPRs are colour coded according to 4 different connectivity matrices. The columns represent 3 different block designs and the rows denote the likelihood ratio (LR) and Wald (W) scores based on the parametric and permutation tests. The red shaded strip represents the 95% Monte Carlo binomial proportion confidence intervals.

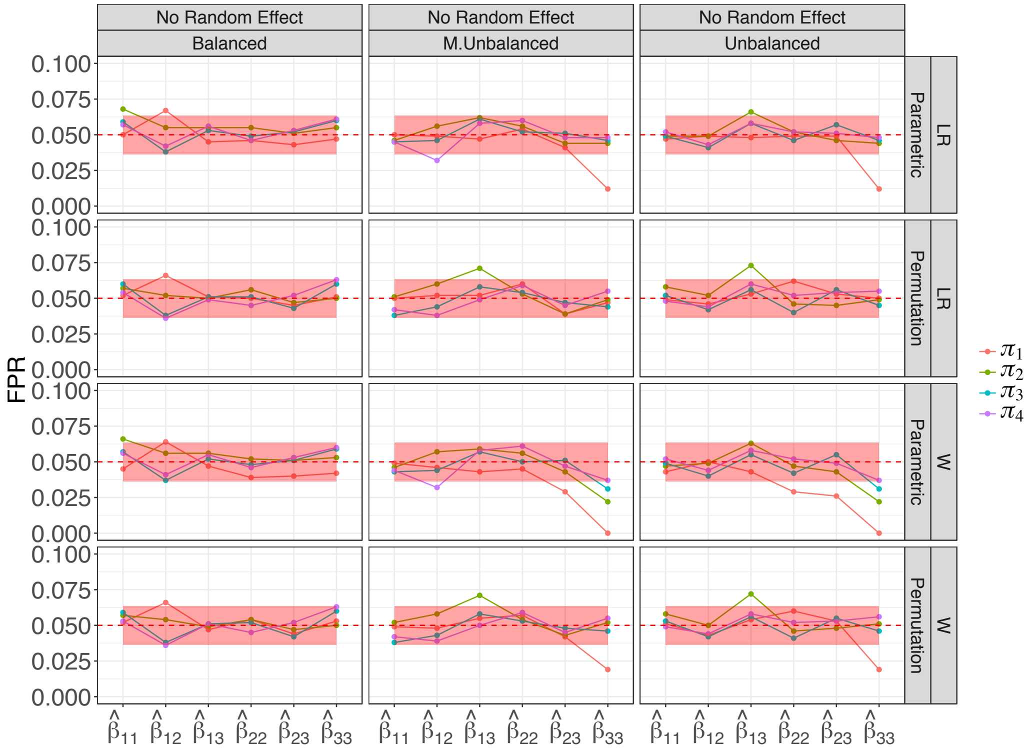


Figure S29: **Observed FPR for the simulated networks with 30 nodes, 40 subjects and no random effect.** The $x$-axis represents the block specific fitted regression coefficients. The observed FPRs are colour coded according to 4 different connectivity matrices. The columns represent 3 different block designs and the rows denote the likelihood ratio (LR) and Wald (W) scores based on the parametric and permutation tests. The red shaded strip represents the 95% Monte Carlo binomial proportion confidence intervals.

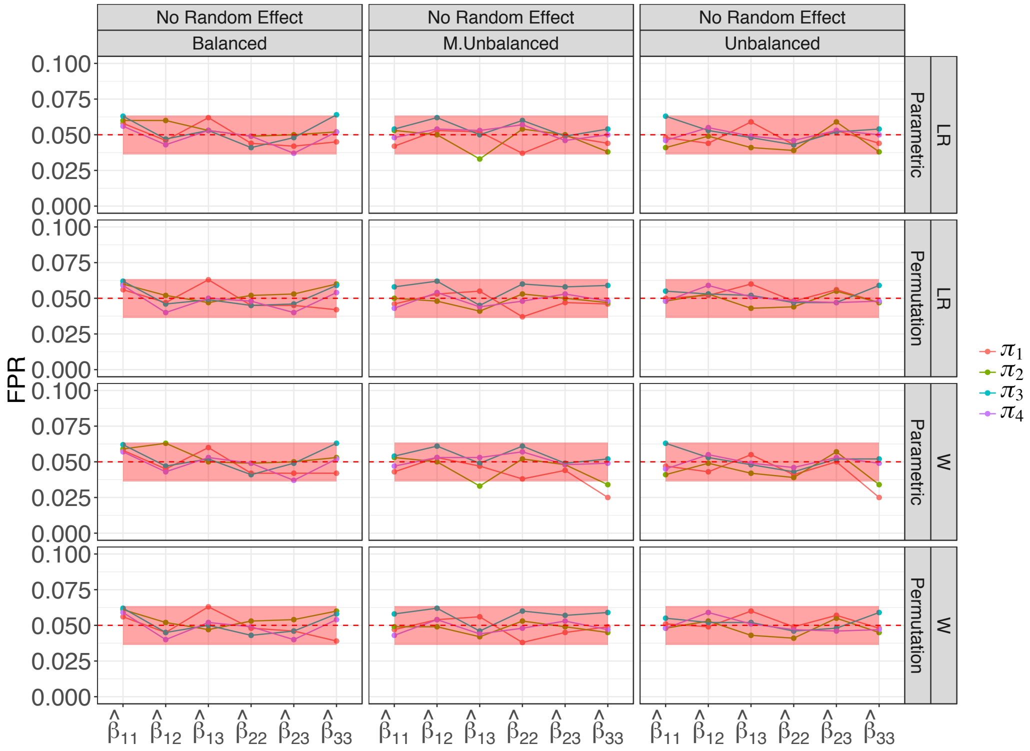


Figure S30: **Observed FPR for the simulated networks with 60 nodes, 40 subjects and no random effect.** The $x$-axis represents the block specific fitted regression coefficients. The observed FPRs are colour coded according to 4 different connectivity matrices. The columns represent 3 different block designs and the rows denote the likelihood ratio (LR) and Wald (W) scores based on the parametric and permutation tests. The red shaded strip represents the 95% Monte Carlo binomial proportion confidence intervals.

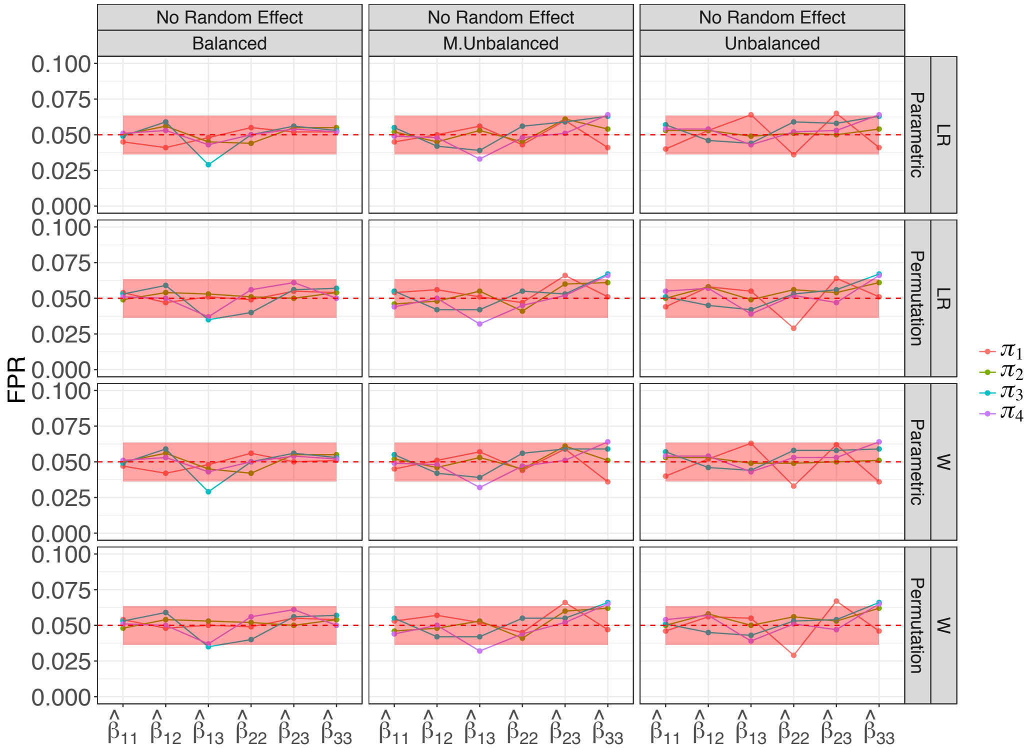


Figure S31: **Observed FPR for the simulated networks with 120 nodes, 40 subjects and no random effect.** The $x$-axis represents the block specific fitted regression coefficients. The observed FPRs are colour coded according to 4 different connectivity matrices. The columns represent 3 different block designs and the rows denote the likelihood ratio (LR) and Wald (W) scores based on the parametric and permutation tests. The red shaded strip represents the 95% Monte Carlo binomial proportion confidence intervals.

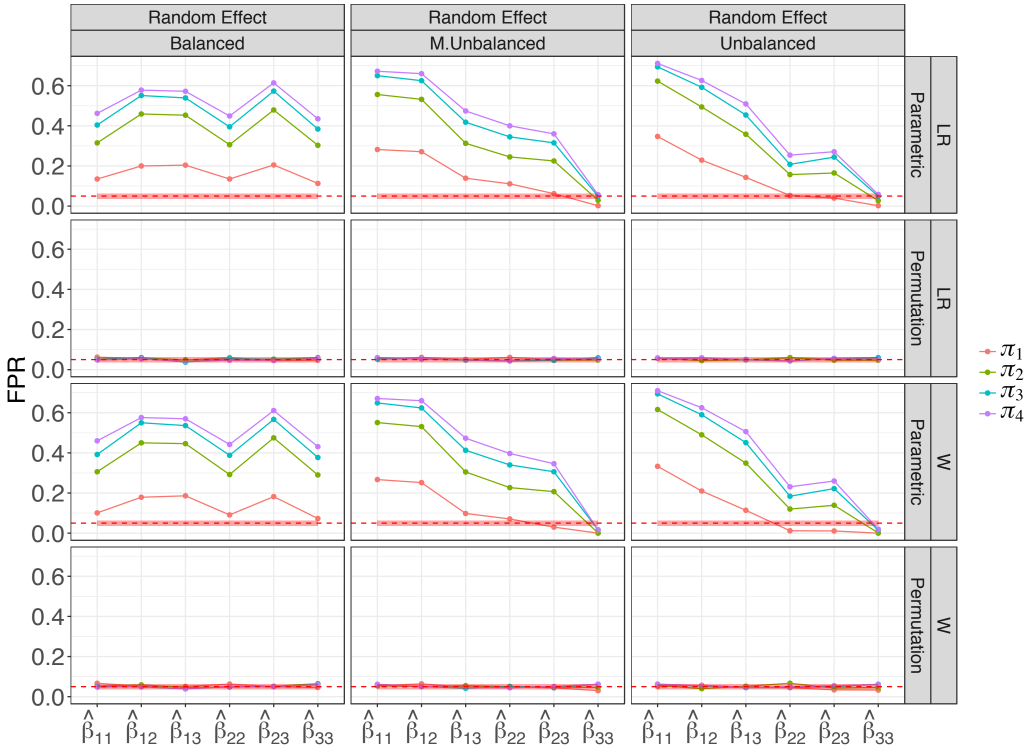


Figure S32: **Observed FPR for the simulated networks with 60 nodes, 10 subjects and random effect.** The $x$-axis represents the block specific fitted regression coefficients. The observed FPRs are colour coded according to 4 different connectivity matrices. The columns represent 3 different block designs and the rows denote the likelihood ratio (LR) and Wald (W) scores based on the parametric and permutation tests. The red shaded strip represents the 95% Monte Carlo binomial proportion confidence intervals.

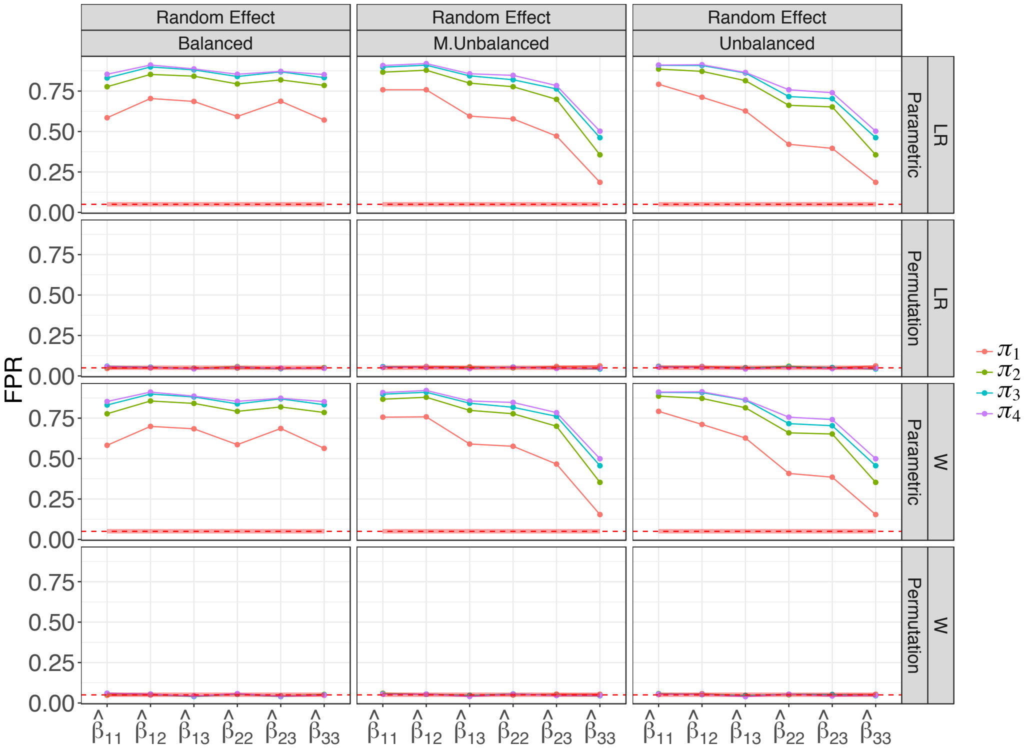


Figure S33: **Observed FPR for the simulated networks with 120 nodes, 10 subjects and random effect.** The $x$-axis represents the block specific fitted regression coefficients. The observed FPRs are colour coded according to 4 different connectivity matrices. The columns represent 3 different block designs and the rows denote the likelihood ratio (LR) and Wald (W) scores based on the parametric and permutation tests. The red shaded strip represents the 95% Monte Carlo binomial proportion confidence intervals.

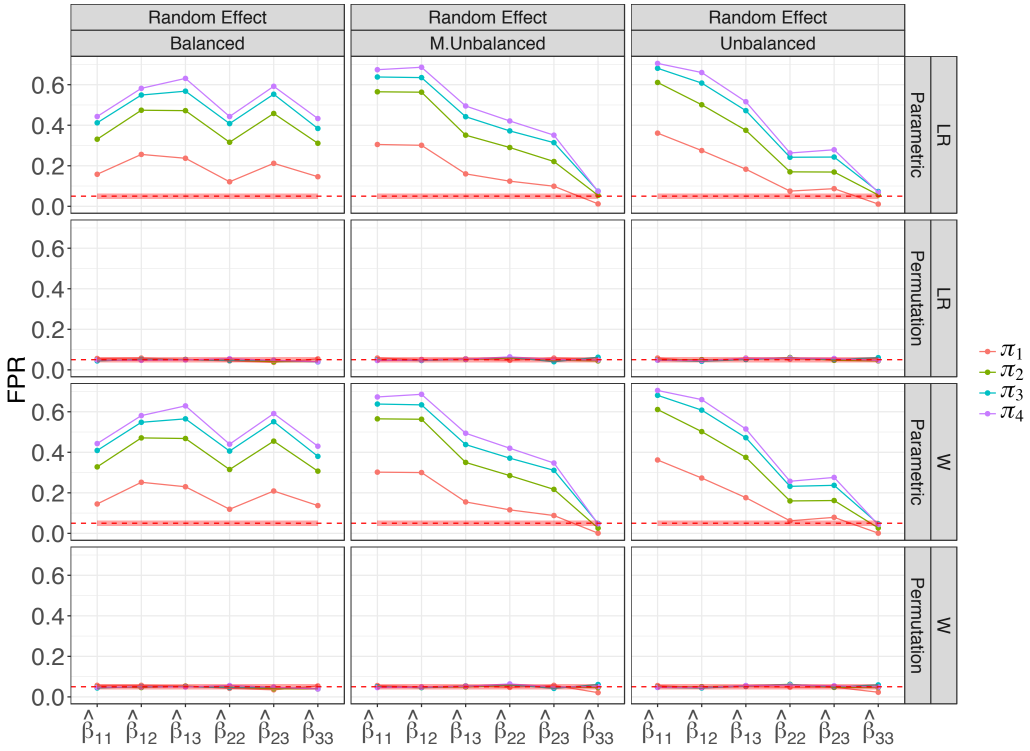


Figure S34: **Observed FPR for the simulated networks with 30 nodes, 20 subjects and random effect.** The $x$-axis represents the block specific fitted regression coefficients. The observed FPRs are colour coded according to 4 different connectivity matrices. The columns represent 3 different block designs and the rows denote the likelihood ratio (LR) and Wald (W) scores based on the parametric and permutation tests. The red shaded strip represents the 95% Monte Carlo binomial proportion confidence intervals.

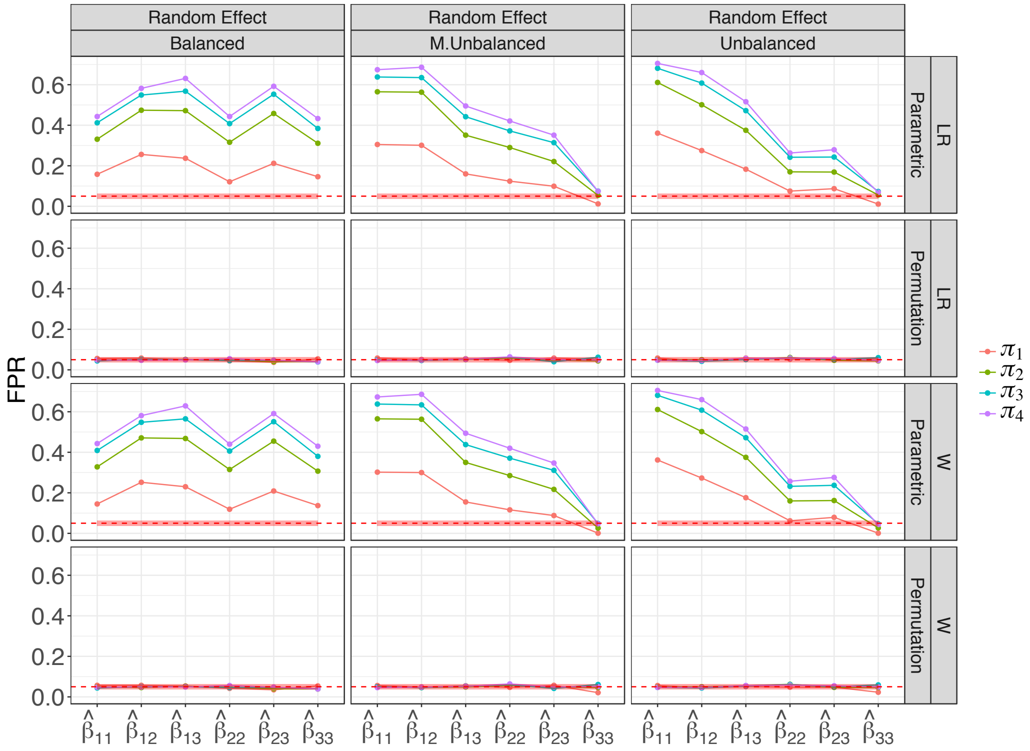


Figure S35: **Observed FPR for the simulated networks with 60 nodes, 20 subjects and random effect.** The $x$-axis represents the block specific fitted regression coefficients. The observed FPRs are colour coded according to 4 different connectivity matrices. The columns represent 3 different block designs and the rows denote the likelihood ratio (LR) and Wald (W) scores based on the parametric and permutation tests. The red shaded strip represents the 95% Monte Carlo binomial proportion confidence intervals.

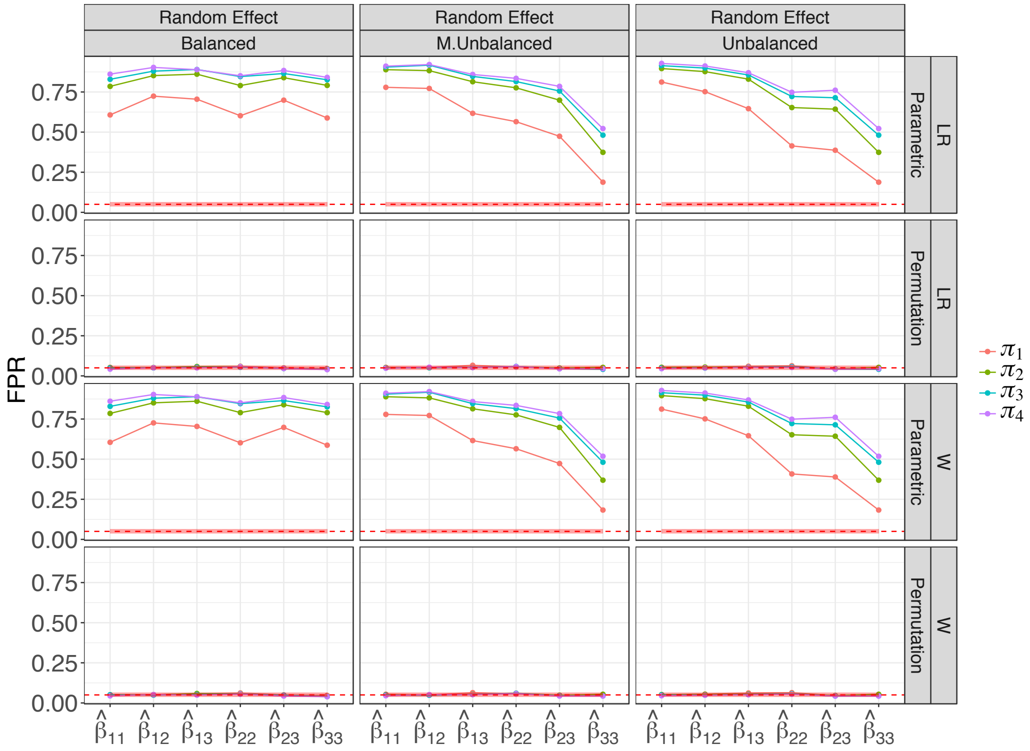


Figure S36: **Observed FPR for the simulated networks with 120 nodes, 20 subjects and random effect.** The $x$-axis represents the block specific fitted regression coefficients. The observed FPRs are colour coded according to 4 different connectivity matrices. The columns represent 3 different block designs and the rows denote the likelihood ratio (LR) and Wald (W) scores based on the parametric and permutation tests. The red shaded strip represents the 95% Monte Carlo binomial proportion confidence intervals.

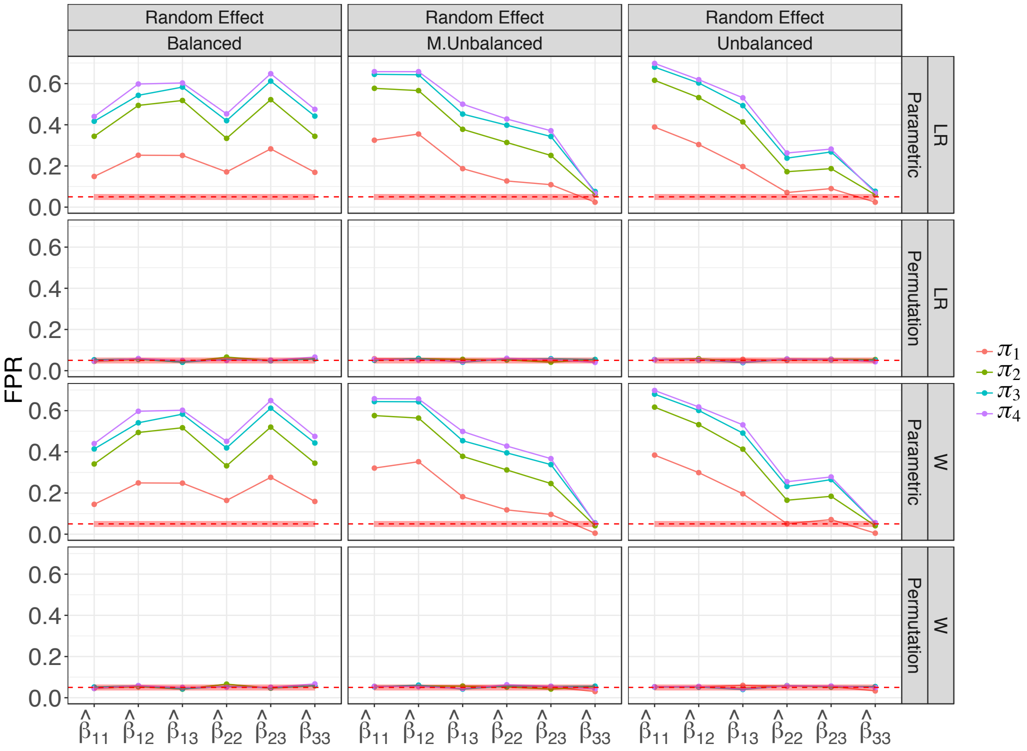


Figure S37: **Observed FPR for the simulated networks with 30 nodes, 40 subjects and random effect.** The $x$-axis represents the block specific fitted regression coefficients. The observed FPRs are colour coded according to 4 different connectivity matrices. The columns represent 3 different block designs and the rows denote the likelihood ratio (LR) and Wald (W) scores based on the parametric and permutation tests. The red shaded strip represents the 95% Monte Carlo binomial proportion confidence intervals.

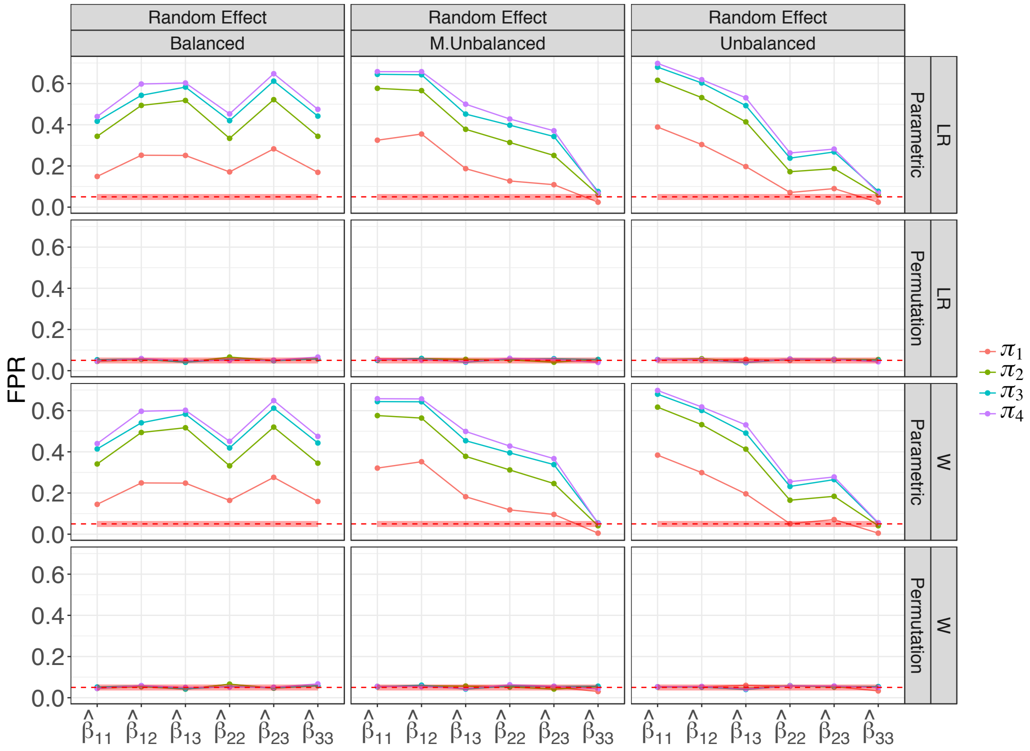


Figure S38: **Observed FPR for the simulated networks with 60 nodes, 40 subjects and random effect.** The $x$-axis represents the block specific fitted regression coefficients. The observed FPRs are colour coded according to 4 different connectivity matrices. The columns represent 3 different block designs and the rows denote the likelihood ratio (LR) and Wald (W) scores based on the parametric and permutation tests. The red shaded strip represents the 95% Monte Carlo binomial proportion confidence intervals.

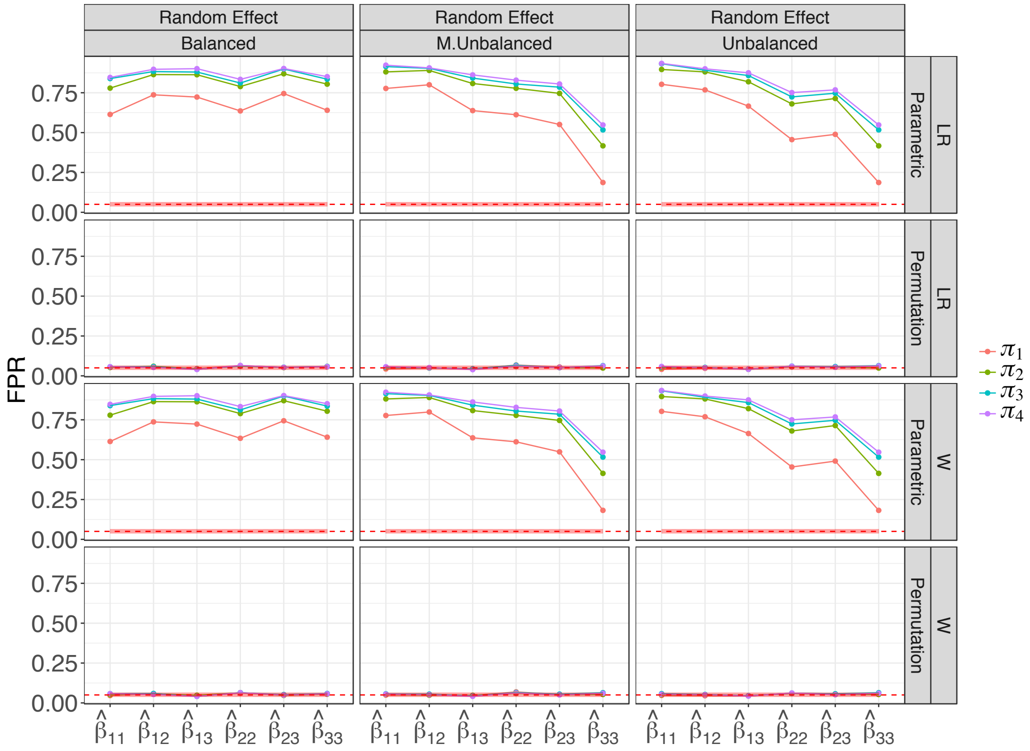


Figure S39: **Observed FPR for the simulated networks with 120 nodes, 40 subjects and random effect.** The $x$-axis represents the block specific fitted regression coefficients. The observed FPRs are colour coded according to 4 different connectivity matrices. The columns represent 3 different block designs and the rows denote the likelihood ratio (LR) and Wald (W) scores based on the parametric and permutation tests. The red shaded strip represents the 95% Monte Carlo binomial proportion confidence intervals.

#### M. Real Data Analysis

In this Section, we present additional information for pre-processing and the multi-subject data fit. During pre-processing, the list of ROIs removed due to the missing values for some subjects is given in Figure S40. Since, these are mostly located in cerebellum, the remaining ROIs of cerebellum were also removed.

| ROIs removed due to missing data | Remaining ROIs of the cerebellum removed |
|---|---|
| Right Superior Temporal Gyrus | Right Cerebellar Tonsil |
| Left Inferior Frontal Gyrus | Right Culmen |
| Left Cerebellar Tonsil | Left Pyramis |
| Right Pyramis | Right Declive |
| Left Inferior Frontal Gyrus | Right Tuber |
| Right Cerebellar Tonsil | Right Uvula |
| Left Parahippocampal Gyrus | Right Tuber |
| Right Cerebellar Tonsil | Left Culmen |
| Right Culmen | Left Uvula |
| Right Uvula | Left Cerebellar Tonsil |
| Left Cerebellar Tonsil | Right Culmen |
| Left Inferior Semi−Lunar Lobule | Left Declive |
| Left Precentral Gyrus | Right Culmen |
| Left Cerebellar Tonsil | Left Declive |
| Right Superior Temporal Gyrus | Right Culmen |
| Left Superior Temporal Gyrus | Left Culmen |
| Right Cerebellar Tonsil | Left Declive |
| Right Culmen | Left Pyramis |
| Right Cerebellar Tonsil | Right Culmen |
| Right Fusiform Gyrus | Left Culmen |
| Left Cerebellar Tonsil | Left Nodule |
| Right Cerebellar Tonsil | Right Declive |
| Left Cerebellar Tonsil | Left Culmen |
| Right Cerebellar Tonsil | Right Declive |
| Left Culmen | Left Culmen |
| Right Inferior Frontal Gyrus | Right Pyramis |
| Right Cerebellar Tonsil | Left Culmen |
| Right Cerebellar Tonsil | Left Culmen |
| | Right Tuber |

Figure S40: **Regions of the brain in both left and right hemispheres excluded from the study.**

##### *M.1. Het-SBM Inference Results*

To investigate if there is a random effect per each block, we extracted the data of each block across all subjects. Then, for each such block, we fit two models, a logistic model (`glm` function from the `stats` R package) with only fixed effects (the same as in the original Het-SBM fit) as the null model and a logistic mixed effect model(`glmer` function from the `lme4` R package) with the same fixed effects and a random intercept. For inference, we use a parametric bootstrap test for nested model using the R package `pbnm` with the LR score as statistic and 1000 bootstraps. To account for the multiple comparison problem (i.e. the total of 230 tests), we use the same bootstraps for each block and compute the maximum LR score for each of the 1000 bootstraps (one of them was the maximum of the original LR scores) and computed a FWER-corrected $P$-value at each block by counting the number of times a maximum LR score was greater than or equal to each of the original LR scores. We discarded Block (21,21) from the analysis as it did not have enough data to fit the models. Only 2 blocks had a FWER-corrected $P$-value greater than 5%, strongly suggesting some form of dependence between the block edges of each subject.

#### N. Additional Results for Schizophrenia Study

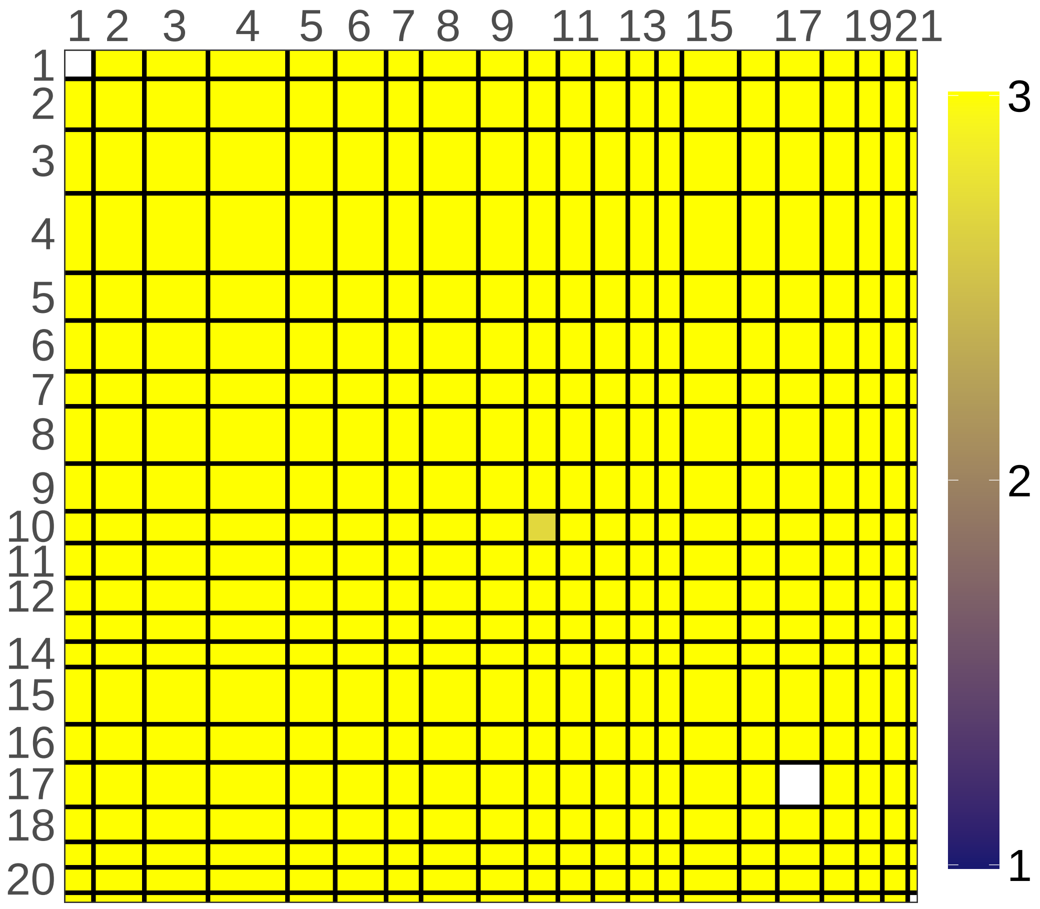


Figure S41: **Parametric bootstrap test with the LR score as statistic for the test of significance of random effect.** Significant FWER-corrected $P$-value are given for each block and they are stated in terms of negative logarithm with base 10 (e.g., the legend score of 3 corresponds to the $P$-value of $10^{-3}$). Non-significant blocks are given in white including Block (21,21) which was excluded form the analysis due to having a small number of data points. With the exception of Block (1,1) and Block (17,17), there is an overall strong evidence of random effect in the remaining block structure.

Block 1

Left Cingulate Gyrus
Left Precuneus
Right Precuneus
Left Cingulate Gyrus
Right Posterior Cingulate
Left Posterior Cingulate
Left Middle Temporal Gyrus
Right Middle Temporal Gyrus
Right Cingulate Gyrus

Block 2

Right Paracentral Lobule
Left Precentral Gyrus
Right Precentral Gyrus
Left Postcentral Gyrus
Left Cingulate Gyrus
Left Paracentral Lobule
Right Superior Frontal Gyrus
Left Precentral Gyrus
Right Postcentral Gyrus
Right Precentral Gyrus
Right Medial Frontal Gyrus
Left Cingulate Gyrus
Right Cingulate Gyrus
Left Inferior Parietal Lobule
Left Postcentral Gyrus
Right Precentral Gyrus

Block 3

Left Angular Gyrus
Left Superior Frontal Gyrus
Left Middle Frontal Gyrus
Right Medial Frontal Gyrus
Left Medial Frontal Gyrus
Left Supramarginal Gyrus
Left Superior Frontal Gyrus
Right Medial Frontal Gyrus
Left Superior Frontal Gyrus
Right Superior Frontal Gyrus
Right Inferior Parietal Lobule
Left Superior Frontal Gyrus
Right Superior Frontal Gyrus
Right Superior Frontal Gyrus
Right Superior Frontal Gyrus
Left Medial Frontal Gyrus
Right Middle Frontal Gyrus
Right Inferior Parietal Lobule
Right Middle Temporal Gyrus
Left Medial Frontal Gyrus

Block 4

Right Middle Occipital Gyrus
Right Lingual Gyrus
Left Fusiform Gyrus
Left Precuneus
Right Posterior Cingulate
Right Precuneus
Right Lingual Gyrus
Left Precuneus
Left Middle Occipital Gyrus
Right Middle Temporal Gyrus
Left Middle Occipital Gyrus
Right Cuneus
Left Cuneus
Left Fusiform Gyrus
Left Cuneus
Right Posterior Cingulate
Left Cuneus
Left Middle Occipital Gyrus
Right Cuneus
Right Fusiform Gyrus
Left Cuneus
Left Cuneus
Left Middle Occipital Gyrus
Left Cuneus
Right Inferior Temporal Gyrus

Block 5

Right Superior Temporal Gyrus
Right Parahippocampal Gyrus
Left Inferior Temporal Gyrus
Left Insula
Left Parahippocampal Gyrus
Right Middle Temporal Gyrus
Right Parahippocampal Gyrus
Right Superior Temporal Gyrus
Right Parahippocampal Gyrus
Right Insula
Left Superior Temporal Gyrus
Left Inferior Frontal Gyrus
Right Parahippocampal Gyrus
Left Parahippocampal Gyrus
Left Middle Temporal Gyrus

Block 6

Right Lentiform Nucleus
Left Caudate
Right Inferior Frontal Gyrus
Left Insula
Left Middle Frontal Gyrus
Left Inferior Frontal
Right Inferior Frontal Gyrus
Right Thalamus
Right Inferior Frontal Gyrus
Left Precentral Gyrus
Right Inferior Frontal Gyrus
Left Inferior Frontal Gyrus
Left Inferior Frontal Gyrus
Left Thalamus
Right Inferior Frontal Gyrus
Left Inferior Frontal Gyrus

Block 7

Left Superior Temporal Gyrus
Left Middle Frontal Gyrus
Right Middle Temporal Gyrus
Right Middle Temporal Gyrus
Left Middle Temporal Gyrus
Left Middle Temporal Gyrus
Right Inferior Frontal Gyrus
Left Middle Temporal Gyrus
Left Precentral Gyrus
Right Middle Temporal Gyrus
Right Inferior Frontal Gyrus

Block 8

Right Cingulate Gyrus
Right Middle Frontal Gyrus
Right Middle Frontal Gyrus
Left Supramarginal Gyrus
Right Supramarginal Gyrus
Right Middle Frontal Gyrus
Right Middle Frontal Gyrus
Left Superior Frontal Gyrus
Left Middle Frontal Gyrus
Left Cingulate Gyrus
Right Superior Frontal Gyrus
Right Middle Frontal Gyrus
Right Middle Frontal Gyrus
Right Cingulate Gyrus
Left Middle Frontal Gyrus
Left Middle Frontal Gyrus
Right Middle Frontal Gyrus
Left Precentral Gyrus

Block 9

Left Postcentral Gyrus
Right Postcentral Gyrus
Right Middle Frontal Gyrus
Left Medial Frontal Gyrus
Left Precentral Gyrus
Left Postcentral Gyrus
Left Middle Frontal Gyrus
Left Postcentral Gyrus
Right Postcentral Gyrus
Right Precuneus
Right Precentral Gyrus
Left Inferior Parietal Lobule
Left Medial Frontal Gyrus
Right Middle Frontal Gyrus
Right Postcentral Gyrus

Block 10

Right Parahippocampal Gyrus
Left Fusiform Gyrus
Right Parahippocampal Gyrus
Left Thalamus
Right Parahippocampal
Right Fusiform Gyrus
Left Lingual Gyrus
Left Fusiform Gyrus
Left Parahippocampal Gyrus
Left Lingual Gyrus

Figure S42: **Anatomical Labels Het-SBM**.

##### Block 11

Right Lingual Gyrus
Left Medial Frontal Gyrus
Left Precuneus
Right Superior Parietal Lobule
Left Superior Parietal Lobule
Left Paracentral Lobule
Right Superior Frontal Gyrus
Right Superior Frontal Gyrus
Left Postcentral Gyrus
Left Precentral Gyrus
Right Paracentral Lobule

##### Block 12

Left Middle Temporal Gyrus
Right Middle Temporal Gyrus
Right Middle Temporal Gyrus
Left Superior Temporal Gyrus
Left Middle Temporal Gyrus
Right Superior Temporal Gyrus
Left Superior Temporal Gyrus
Right Superior Temporal Gyrus
Right Superior Temporal Gyrus
Right Superior Temporal Gyrus
Left Insula

##### Block 13

Right Superior Parietal Lobule
Right Superior Frontal Gyrus
Right Inferior Temporal Gyrus
Right Middle Frontal Gyrus
Left Middle Frontal Gyrus
Left Middle Temporal Gyrus
Right Superior Parietal Lobule
Left Inferior Temporal Gyrus
Left Superior Frontal Gyrus

##### Block 14

Right Medial Frontal Gyrus
Left Medial Frontal Gyrus
Right Medial Frontal Gyrus
Right Superior Frontal Gyrus
Left Anterior Cingulate
Left Medial Frontal Gyrus
Left Medial Frontal Gyrus
Left Superior Frontal Gyrus

##### Block 15

Left Insula
Right Insula
Right Insula
Right Inferior Parietal Lobule
Left Postcentral Gyrus
Right Postcentral Gyrus
Right Postcentral Gyrus
Right Insula
Left Superior Temporal Gyrus
Left Inferior Parietal Lobule
Left Insula
Right Precentral Gyrus
Left Precentral Gyrus
Left Insula
Right Inferior Parietal Lobule
Left Insula
Right Postcentral Gyrus
Left Precentral Gyrus

##### Block 16

Right Caudate
Right Anterior Cingulate
Left Caudate
Right Middle Temporal Gyrus
Right Medial Frontal Gyrus
Left Subcallosal Gyrus
Left Superior Frontal
Right Middle Frontal Gyrus
Right Middle Temporal Gyrus
Left Medial Frontal Gyrus
Right Middle Frontal Gyrus
Right Middle Frontal Gyrus

##### Block 17

Right Precuneus
Right Precuneus
Left Precuneus
Right Precuneus
Right Angular Gyrus
Right Precuneus
Left Inferior Parietal Lobule
Right Precuneus
Left Precuneus
Right Inferior Parietal Lobule
Right Precuneus
Right Inferior Parietal Lobule
Left Inferior Parietal Lobule
Left Precuneus

##### Block 18

Right Insula
Right Parahippocampal Gyrus
Right Thalamus
Left Parahippocampal Gyrus
Right Lentiform Nucleus
Left Insula
Left Thalamus
Left Subcallosal Gyrus
Right Inferior Frontal Gyrus
Left Lentiform Nucleus
Left Superior Temporal Gyrus

##### Block 19

Right Superior Frontal Gyrus
Left Superior Frontal Gyrus
Right Middle Frontal Gyrus
Right Lentiform Nucleus
Right Cingulate Gyrus
Left Caudate
Right Middle Frontal Gyrus
Left Middle Frontal Gyrus

##### Block 20

Right Cuneus
Left Middle Occipital
Right Inferior Occipital Gyrus
Right Cuneus
Left Cuneus
Left Inferior Occipital Gyrus
Right Cuneus
Left Lingual Gyrus

##### Block 21

Left Anterior Cingulate
Right Anterior Cingulate
Right Anterior Cingulate

Figure S43: **Anatomical Labels Het-SBM**.
